# Genome-wide coancestry reveals details of ancient and recent male-driven reticulation in baboons

**DOI:** 10.1101/2023.05.02.539112

**Authors:** Erik F. Sørensen, R. Alan Harris, Liye Zhang, Muthuswamy Raveendran, Lukas F. K. Kuderna, Jerilyn A. Walker, Jessica M. Storer, Martin Kuhlwilm, Claudia Fontsere, Lakshmi Seshadri, Christina M. Bergey, Andrew S. Burrell, Juraj Bergmann, Jane E. Phillips-Conroy, Fekadu Shiferaw, Kenneth L. Chiou, Idrissa S. Chuma, Julius D. Keyyu, Julia Fischer, Marie-Claude Gingras, Sejal Salvi, Harshavardhan Doddapaneni, Mikkel H. Schierup, Mark A. Batzer, Clifford J. Jolly, Sascha Knauf, Dietmar Zinner, Kyle K.-H. Farh, Tomas Marques-Bonet, Kasper Munch, Christian Roos, Jeffrey Rogers

## Abstract

Baboons (genus *Papio*) are a morphologically and behaviorally diverse clade of catarrhine monkeys that have experienced hybridization between phenotypically and genetically distinct phylogenetic species. We used high coverage whole genome sequences from 225 wild baboons representing 19 geographic localities to investigate population genomics and inter-species gene flow. Our analyses provide an expanded picture of evolutionary reticulation among species and reveal novel patterns of population structure within and among species, including differential admixture among conspecific populations. We describe the first example of a baboon population with a genetic composition that is derived from three distinct lineages. The results reveal processes, both ancient and recent, that produced the observed mismatch between phylogenetic relationships based on matrilineal, patrilineal, and biparental inheritance. We also identified several candidate genes that may contribute to species-specific phenotypes.

**One-Sentence Summary:** Genomic data for 225 baboons reveal novel sites of inter-species gene flow and local effects due to differences in admixture.

---

Our understanding of the evolutionary processes involved in the origin of biological diversity has changed significantly over the past two decades. Genetic analyses have demonstrated that hybridization and inter-species gene flow between closely related mammalian species occur more often than previously assumed *(1, 2)*. Traditional studies of natural hybridization among populations and species have relied on phenotypic variation and a few informative genetic markers *(3, 4)*. However, access to large-scale genomic datasets now allows more extensive analyses *(5–7)* demonstrating that in some cases complex reticulations rather than dichotomously branching phylogenetic trees more accurately represent evolutionary histories.

Among primates, including humans, the number of genera found to exhibit complex histories of interspecific reticulation has recently grown considerably *(2, 8–12)*. Baboons (genus *Papio*) have long been recognized as a prime example of inter-species gene flow with several hybrid zones between the six currently recognized parapatric species (Guinea baboons *P. papio*; hamadryas baboons *P. hamadryas*; olive baboons *P. anubis*; yellow baboons *P. cynocephalus*; Kinda baboons *P. kindae*; chacma baboons *P. ursinus*; Fig. 1; for the rationale behind the classification of these major forms as species rather than subspecies see *(13)*) *(14–17)*. Previous analyses have identified substantial discrepancies in species-level phylogenies based on nuclear DNA, mitochondrial DNA (mtDNA), and phenotypes, indicating para-and polyphyletic relationships, and suggesting a complex history of differentiation and admixture *(18–21)*. Recent comparisons of whole-genome sequence (WGS) data across *Papio* species illustrated the extent of genetic exchange between phenotypically distinct species *(22–25)*. These studies were, however, restricted to one or two populations per species, and therefore unable to analyze wider geographic patterns of genetic diversity or compare the local effects of interspecific contact.

**Fig. 1.**
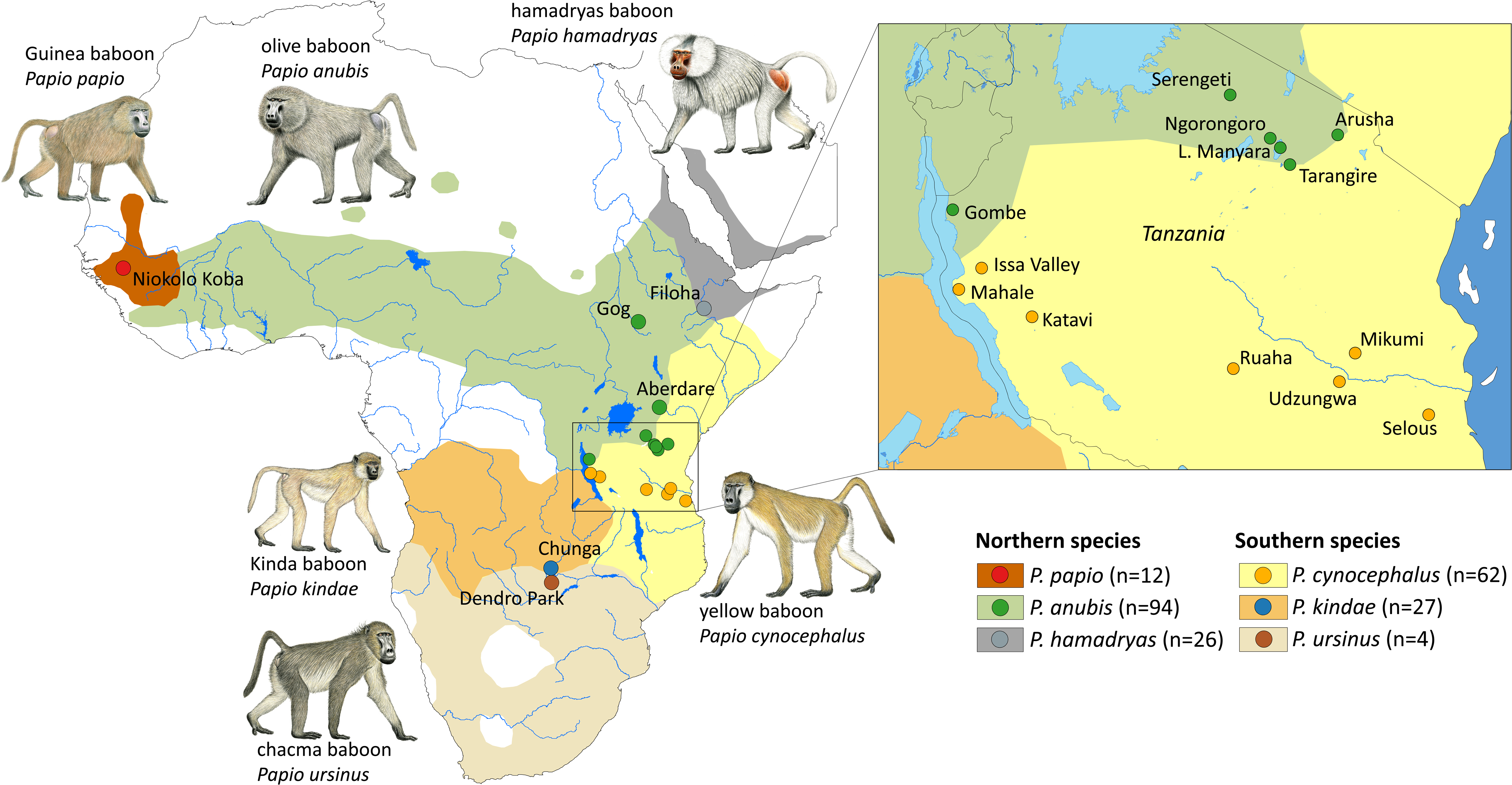
Distribution of the six baboon species and sampling sites. Species distributions are modified from *(20)*. The insert map shows sampling sites in Tanzania. Drawings of male baboons by Stephen Nash, used with permission. Numbers of samples per species are given in parentheses.

This study provides the first detailed WGS-based analysis of coancestry and genomic exchange across all six baboon species, including multiple populations within olive and yellow baboons. We generated deep (>30x; table S1, *(13)*) WGS data from 225 wild baboons representing 19 localities (Fig. 1, table S2), describing variation within and among localities for autosomes, X-and Y-chromosomes, mtDNA, and other genetic features such as insertions of *Alu* repeats and long interspersed elements (LINEs). In addition to population structure using autosomal single nucleotide variants (SNVs) and repetitive elements, we contrast coancestry inferred from autosomal and X-chromosomal data to reveal sex-biased effects on genetic population structure. Our results provide the most extensive analysis of genetic diversity in baboons to date and reveal processes, both ancient and recent, that produced the observed mismatch between phylogenetic relationships based on matrilineal, patrilineal, and biparental inheritance. The evidence indicates the radiation that produced the six extant species began more than one million years ago. The lineages that diverged around that time have since experienced extensive admixture, as reflected in their current genetic composition. We suggest that these findings inform predictions for similar systems such as hominin and early human evolution, for which baboons have long been recognized as a model *(26–29)*.

## RESULTS

WGS analysis across multiple populations of baboons provides a fine-grained picture of present-day population structure and the evolutionary history that generated it. Results of this analysis also document additional locations of ongoing admixture among genetically distinct lineages. Our analyses of SNVs strongly support the existence of differentiated clades including the six recognized species, despite well-known hybrid zones between parapatric species. The initial divergence of evolutionary lineages separates the three northern species (hamadryas, olive, and Guinea baboons) from the three southern species (Kinda, yellow, and chacma baboons). Analyses of population structure (Fig. 2, A to C, figs. S1 to S4) and phylogenomic maximum-likelihood (ML) trees using autosomal, X-and Y-chromosomal, and mtDNA data (figs. S5 to S8) are consistent with the initial north-south split, and with greater overall divergence among southern than northern baboons (see also *(23)*). Principal component analyses (PCAs) and ML trees of autosomal and X-chromosomal data separate the western Tanzanian yellow baboons located at Mahale and Katavi into their own cluster distinct from eastern Tanzanian yellow baboons from Mikumi, Selous, Ruaha and Udzungwa as well as from Kinda baboons. However, the Y-chromosomal phylogenies, including one based on *Alu* insertions (fig. S9), show six main clusters largely corresponding to the six species and place most western yellow baboons with Kinda baboons. Other western yellow baboons cluster in that analysis with eastern yellow and one olive baboon, providing a clear example of admixture processes not revealed by the whole-genome phylogeny.

**Fig. 2.**
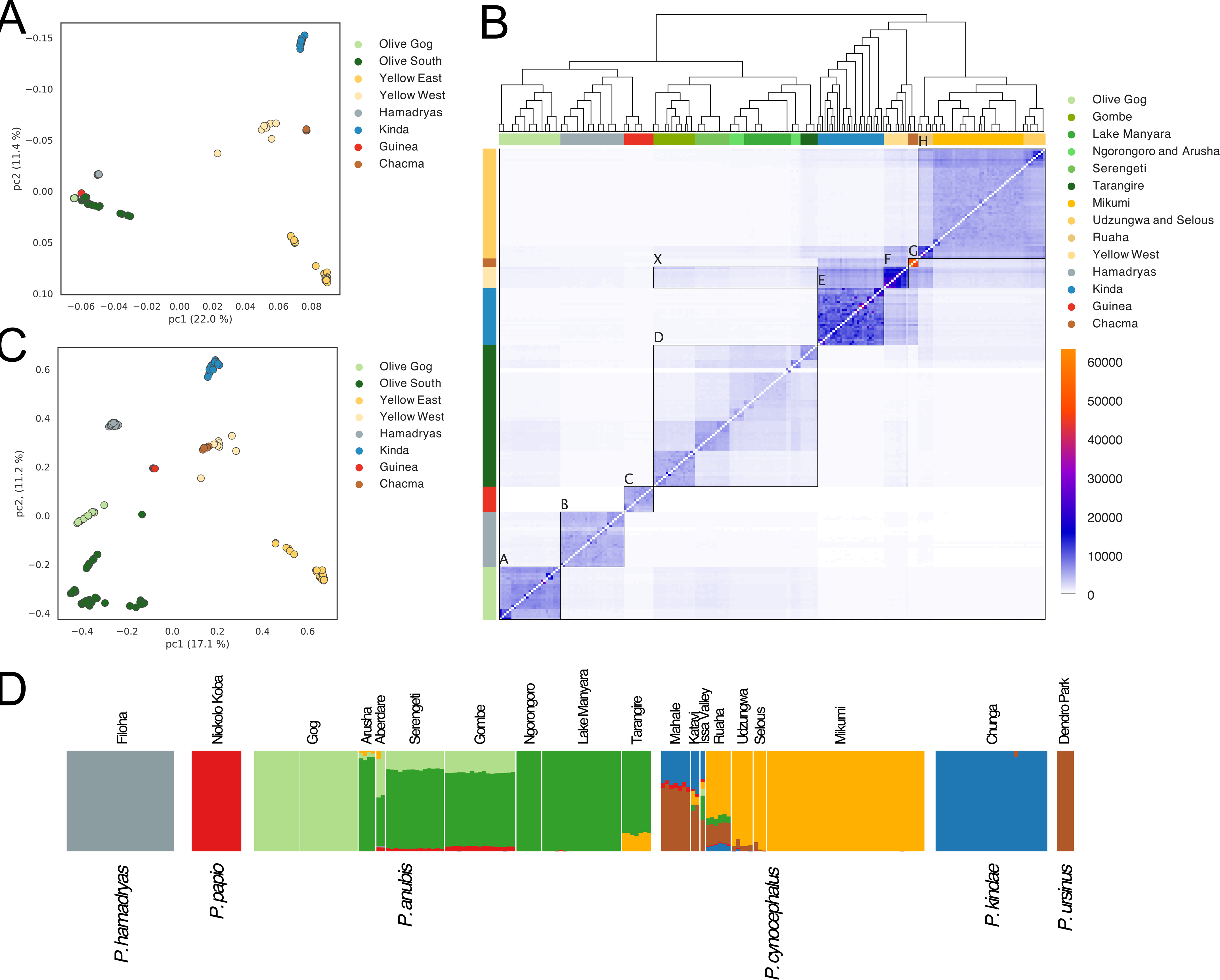
Population structure and coancestry of the six baboon species. (**A**) PCA of autosomal SNVs. (**B**) ChromoPainter coancestry matrix with fineSTRUCTURE dendrogram. Each row in the coancestry matrix represents an individual and illustrates how its most recent common ancestry is distributed across all other sampled individuals. The ordering of individuals is the same for rows and columns. The row color labels are the same as in A and correspond to clusters shown for eight populations labeled with boxes: A: Gog olive (Ethiopia), B: hamadryas, C: Guinea, D: southern olive (Kenya and Tanzania), E: Kinda, F: western yellow, G: chacma, H: eastern yellow; X: olive coancestry in western yellows suggesting admixture (see alternate fineSTRUCTURE figure fig. S4). Colors labels below the dendrogram represent the 14 groups named in the figure legend. (**C**) PCA of the coancestry matrix. (**D**) ADMIXTURE plot with the preferred grouping of baboons into seven clusters (*K* = 7; for *K* = 2-10 see fig. S12).

Across the genome of each individual, we identified the most recent coancestry among all other sampled individuals (ChromoPainter *(30)*). The corresponding first two principal components (Fig. 2C) show extensive variation among yellow baboons and confirm the primary north-south split. This north-south split is also apparent in the clustering using fineSTRUCTURE *(30)* (Fig. 2B). ML trees for autosomes and X-and Y-chromosomes (figs. S5 to S7) all support the conclusions based on PCA, with two individuals falling outside their expected species clades (samples PD0266 and PD0662, also anomalous in the PCAs; figs. S1, S2, S10 and S11, *(13)*). As discussed below, the Y-chromosomal phylogeny places Kinda baboons basal to all others (fig. S7).

Unsupervised cluster algorithms group individuals largely by species (see ADMIXTURE analysis; Fig. 2D, fig. S12) with *K* = 7 as the preferred number of clusters. However, in species for which we sampled more than one population (olive and yellow baboons), we find local genetic differences and evidence for a complex evolutionary history (detailed discussion below). These results are also supported by an analysis of LINE-1 (L1) insertions (fig. S13), an independent class of genetic marker that is less prone to parallel mutations. The pelage phenotypes on which taxonomy was traditionally based are generally very consistent within species over wide geographic ranges *(31)*. Yet, we find high genomic variation within and among conspecific populations. Heterozygosity ranges from 0.0006 to 0.0026 (average 0.0018) per base pair across the six species, and from 0.0006 to 0.0029 across the 19 localities, with the lowest values in Guinea baboons (table S3, figs. S14 to S17). The coancestry matrix and its PCA (Fig. 2, B and C) differentiates the various sampling localities and is therefore consistent with the ADMIXTURE analysis (Fig. 2D) showing that the sampled populations within both yellow and olive baboons can be distinguished genetically. The yellow baboons in Mikumi (Fig. 2B, box H) share pelage and morphological phenotypes with those in Ruaha, although they are genetically distinct. Western yellow baboons from Mahale and Katavi (Fig. 2B, box F) exhibit phenotypic traits (somewhat smaller body size than Mikumi baboons, especially cranial metrics; aspects of coat color with some individuals having pink skin around the eyes, and sporadic occurrence of white-furred infants) in which they resemble Kinda baboons *(32)*. The coancestry matrix (Fig. 2B) further shows that yellow baboons from Mahale and Katavi (box F) exhibit greater genetic similarity with Kinda (box E) and chacma baboons (box G) than with their supposed conspecifics from eastern Tanzania (box H). Similarly, all olive baboons (with exception of those from Tarangire) share a very consistent pelage and external phenotype. However, ADMIXTURE (Fig. 2D) and ChromoPainter (Fig. 2B) analyses identify clear evidence of genetic differences between the Ethiopian Gog olive baboons and the Tanzanian olive baboons of Lake Manyara and Ngorongoro. Furthermore, the Serengeti population is more similar genetically to both the Gombe and Aberdare populations than to the Ngorongoro or Lake Manyara populations which are geographically much closer.

We used the SNV data to reconstruct the history of population size for each baboon locality (Fig. 3A, figs. S18 to S21). The estimated effective population sizes (N_e_) are all essentially the same and on the order of 100,000 until about 1.0-1.2 million years ago, which is consistent with the prior dating of the initial north-south divergence *(23)*. At the separation, the N_e_ of northern populations fell below that of the southern populations, supporting the idea that the genus arose in southern Africa, and a daughter population from this basal stock spread to the north, then to the west, losing genetic diversity in serial founding events. The suggestion that Guinea baboons represent the descendants of those groups that were at the leading edge of that dispersal for the longest distance and time *(33)* is supported by the lower heterozygosity in that sample relative to all other baboon species (table S3). Also, whole-genome *Alu* and L1 insertion-based phylogenies place western yellow baboons with Kinda baboons, while Guinea baboons are basal among baboons, and hamadryas baboons are the sister taxon to olive and southern baboons (figs. S22 and S23). These findings may result from Guinea baboons, and to a lesser extent hamadryas baboons, losing polymorphic derived *Alu* and L1 insertions through drift as they dispersed north from the southern geographic origin *(34)*.

**Fig. 3.**
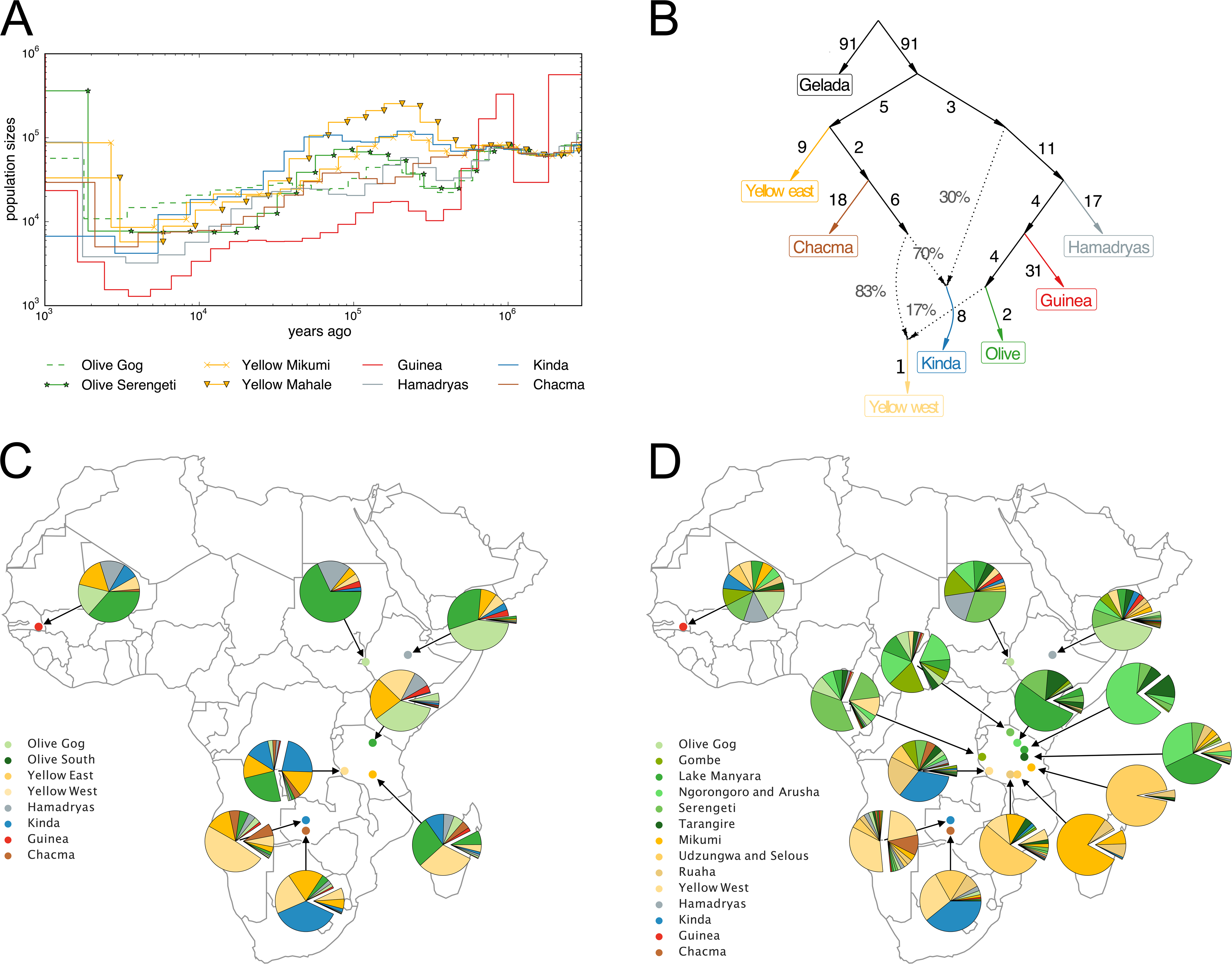
Population history and complex reticulation between baboon populations. (**A**) MSMC2 plots using a mutation rate of 0.9×10^-8^ and a generation time of 11 years *(23)*. (**B**) Admixture graph of the populations used in this study, based on 48,730,011 SNPs with data for all individuals, and a predefined number of two admixture events. Numbers on solid branches correspond to the estimated drift in f2 units of squared frequency difference; labels on dotted edges give admixture proportions. (**C**) Globetrotter analysis of the eight major regional populations. The pie chart for each cluster shows ancestry contributions from other clusters. Expanded wedges represent ancestry that can be attributed to recent admixture (< 56 generations, bootstrap p-values < 0.05). (**D**) Same as C but for 14 populations separating each major sampling location (here, expanded wedges represent ancestry that can be attributed to admixture more recent than 95 generations, bootstrap p-values < 0.05).

Earlier studies provided clear evidence for hybridization and gene flow across the contact zones between pairs of parapatric species *(15–17, 24, 25, 35)*. In this study, we present new evidence for additional ancient and recent arenas for gene flow between species pairs. Species tree reconstruction (ASTRAL *(36)*) using window-based ML trees (50kb and 500kb window size) produced inconsistent branching patterns among datasets and only 58-70% of gene trees fit the species tree at the quartet level (figs. S24 and S25). Both incomplete lineage sorting (ILS) and gene flow are likely contributing to this discordance which is expected to be larger for smaller windows. In addition, a qualitative visualization of these trees (figs. S24 and S25) shows a network-like pattern, again indicating complexity. There is greater shared genetic drift (measured by f3 outgroup statistics) among eastern yellow baboon localities (Udzungwa, Selous, Mikumi, Ruaha), while western yellow baboons tend to cluster with Kinda baboons (fig. S26). In admixture graphs (Fig. 3B), Kinda baboons are, similarly to the description in *(23)*, represented as a fusion product of populations from southern and ancestral northern clades, while the western yellow baboons share ancestry with both Kinda and olive baboons. More complex graphs (tables S4 and S5, figs. S27 to S29) might be supported, but failed to give replicable results, likely due to complex reticulation and multiple gene flow events at different times and between different local populations which now obscure the processes involved.

Taken as a whole, this expanded dataset does not support the previous suggestion that Kinda baboons result from a recent fusion event *(23)* as shown in Fig. 3B. In PCA plots using genome wide SNVs, Kinda baboons do not fall intermediate between northern and southern clades but in fact are quite distinct (Fig. 2A, figs. S1 and S2). Some ML trees (i.e., Y-chromosome data; fig. S7) place Kinda baboons as sister clade to all other baboons whereas other trees (autosomes and X-chromosome data, figs. S5 and S6) lump them together with yellow and chacma baboons into the southern clade. These results are more consistent with the idea that Kinda baboons show substantial genetic similarity to both northern and southern clade baboons because they are basal and phenotypically resemble the ancestral form from which all extant species are derived. Fossil evidence suggests a southern African origin for baboons *(34)*, and the mtDNA haplotypes of Kinda and western yellow baboons (Fig. 4, fig. S8 *(21)*) suggest their range in tropical southern Africa may include the area of origin of both northern and southern primary branches. Broader aspects of Y-chromosome data also do not support Kinda baboons as a fusion product; Kinda baboon Y-haplotypes are found in western yellow baboons but not in olive baboons, and no olive baboon mtDNA has been observed in any Kinda baboon to date. Finally, Kinda baboons share more polymorphic *Alu* insertions with geladas than do other *Papio* species, possibly the result of a period of co-existence and hybridization between their ancestors *(37)*.

**Fig. 4.**
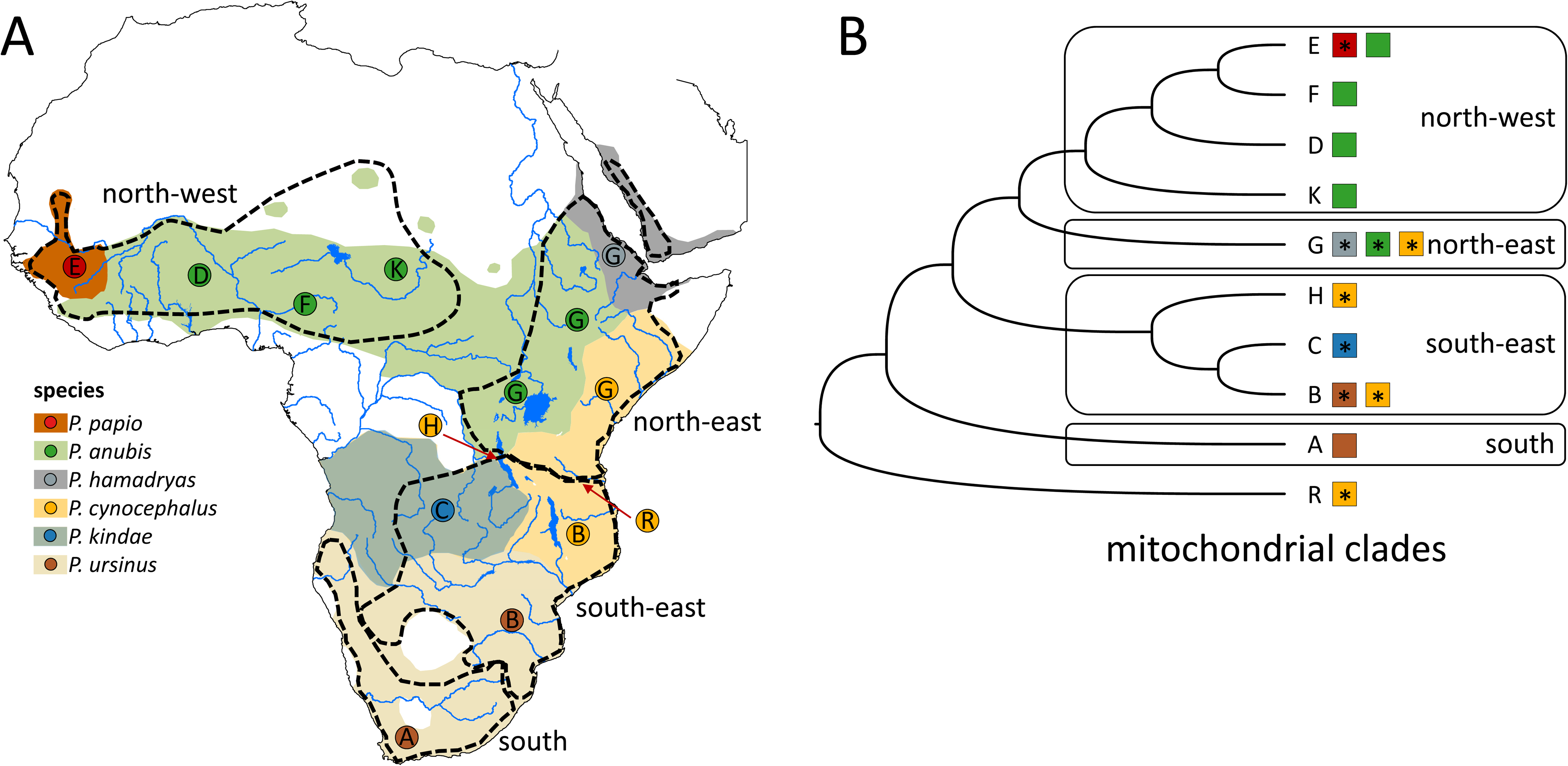
Geographic distribution of mtDNA clades and mtDNA phylogeny. (**A**) Distribution ranges of baboon species and the four main mtDNA clades (south, south-east, north-east, north-west, dashed lines) including major mitochondrial lineages (A-R). (**B**) Phylogeny based on complete mtDNA genomes (see also fig. S8). Clade designation follows *(20, 21)*, asterisks indicate lineages from which mtDNA genomes have been generated in this study. For identical haplotypes see table S7.

We analyzed the genetic relationships among the eight major regional baboon populations that constitute our samples, i.e., the four single-locality populations of chacma, Kinda, hamadryas and Guinea baboons as well as two groups each of yellow (western and eastern) and olive (Gog and southern) baboons. By modeling the recent ancestry along the chromosomes of individual baboons (Globetrotter *(38)*) we can represent each group as a mixture of recent ancestry with the remaining seven groups (Fig. 3C). In most of the groups, we can identify a contribution from recent admixture events (the oldest identifiable event estimated at 56 generations; table S6) separate from contributions of older admixture and retention of ancestral polymorphism (bootstrap p-values < 0.01 unless otherwise noted). In Fig. 3, C and D, we distinguish the recent admixture from more ancient shared ancestry by showing the recent admixture estimates as expanded (exploded) wedges.

We identified a large amount of shared ancestry between southern olive and eastern yellow baboons not concordant with the overall phylogeny (Fig. 3C). This is also expressed in the coancestry matrix (Fig. 2B, box X) and is additional evidence of persistent admixture between both species *(15, 17, 22, 25)*. Furthermore, western yellow baboons from Mahale and Katavi share substantial ancestry with eastern yellow, Kinda, and southern olive baboons. This cannot be explained as a retention of ancient shared variation present prior to the origin of the six major branches, because there is no equivalent sharing with chacma, hamadryas or Guinea baboons. This is, therefore, the first evidence that a single population (western yellow baboons) contains measurable admixture contributions from more than two distinct lineages. Comparing the ancestry of recently admixing populations (expanded wedges in Fig. 3C) to that of each other group identifies recent admixture from Gog into southern olive baboons, between western and eastern yellow baboons, from southern olive baboons into eastern yellow baboons (p-value 0.04), between Kinda and chacma baboons (p-value 0.02), and between Kinda and western yellow baboons. Repeating the Globetrotter analysis assuming 14 populations representing all major sampling locations differentiates olive and yellow baboon populations (Fig. 3D) and reveals a complex system of recent gene flow (all events < 95 generations) between: i) olive baboon populations, ii) yellow baboon populations, iii) yellow and Tarangire olive baboons, iv) western yellow and Gombe olive baboons, and v) Tarangire olive baboons and Ruaha yellow baboons. These results do not imply direct migration of males (e.g., individual males moving from Gog to Serengeti), but more plausibly the overall consequences of many incremental gene flow events distributing alleles long distances over multiple generations.

This is not the first study to suggest that the history of genetic differentiation and reticulation among baboons is complex. Previous studies *(10, 18–21, 33, 39, 40)* showing widespread phenotype-mitochondrial discordance strongly suggest that nuclear swamping (i.e. the immigration of males into a phenotypically different population, largely or completely displacing the nuclear DNA composition and phenotype of the invaded population, without changing its mtDNA composition) has been a major contributing process. The present study found a similar discordance between the expanded mtDNA phylogeny (Fig. 4, fig. S8) on the one hand and the new autosomal and Y-chromosomal phylogenies on the other (figs. S5 and S7). Thus, our WGS findings strongly support previous suggestions based only on mtDNA and phenotype data that nuclear swamping has been a major factor generating the current pattern of baboon genetic and phenotypic variation.

The dense sampling of mtDNA provides important information about matrilineal ancestry. However, as a single locus, mtDNA represents only one of many possible genealogies generated by ILS and admixture. To test the hypothesis that nuclear swamping produced the discord observed between mtDNA phylogenies and relationships based on phenotype, we contrasted ancestry proportions across the X-chromosome and the similar-sized chromosome 8, each contributing thousands of individual genealogies. Admixture by hemizygous males introduces disproportionately more autosomal than X-chromosomal sequence, rendering shared X-chromosome ancestry a better representation of deep species relationships prior to admixture. We found that the X-chromosome of our chacma baboons derives more ancestry from yellow baboons than their chromosome 8 does (0.47 vs 0.62, paired t-test value 0.005; Fig. 5A), suggesting that male-biased admixture from the ancestors of chacma baboons into the southern range of yellow baboons produced northern chacma baboons, including the grayfooted chacma baboons (*P. ursinus grisiepes*) that we analyze here. This observation is consistent with the close relationship between mtDNA found in southern-most yellow and northern chacma baboons (clade B in Fig. 4 *(19, 40)*). The most compelling evidence of male-biased admixture is the relationship between western yellow and Kinda baboons. The ancestry profile of western yellow baboons (Fig. 5B) is very different from eastern yellow baboons (Fig. 5C). Western yellow baboons share more ancestry with Kinda baboons on the X-chromosome than on chromosome 8 (0.27 vs 0.44, paired t-test p-value 0.025) while Kinda baboons contain twice as much western yellow baboon ancestry on the X-chromosome as on chromosome 8 (0.23 vs 0.55, paired t-test p-value 1.8e-13; Fig. 5D). Furthermore, eastern yellow baboons share more X-chromosomal ancestry with western yellow baboons than chromosome 8 ancestry (0.16 vs 0.20, paired t-test p-value 3.1e-9; Fig. 5B). Together these observations indicate that western yellow baboons were produced mainly from males carrying haplotypes that originated among eastern yellow and southern olive baboons migrating into the ancestral range of Kinda baboons, replacing Kinda baboon autosomes more than they replaced Kinda baboon X-chromosomes. As a result, western yellow baboons carry genetic input from three distinct lineages.

**Fig. 5.**
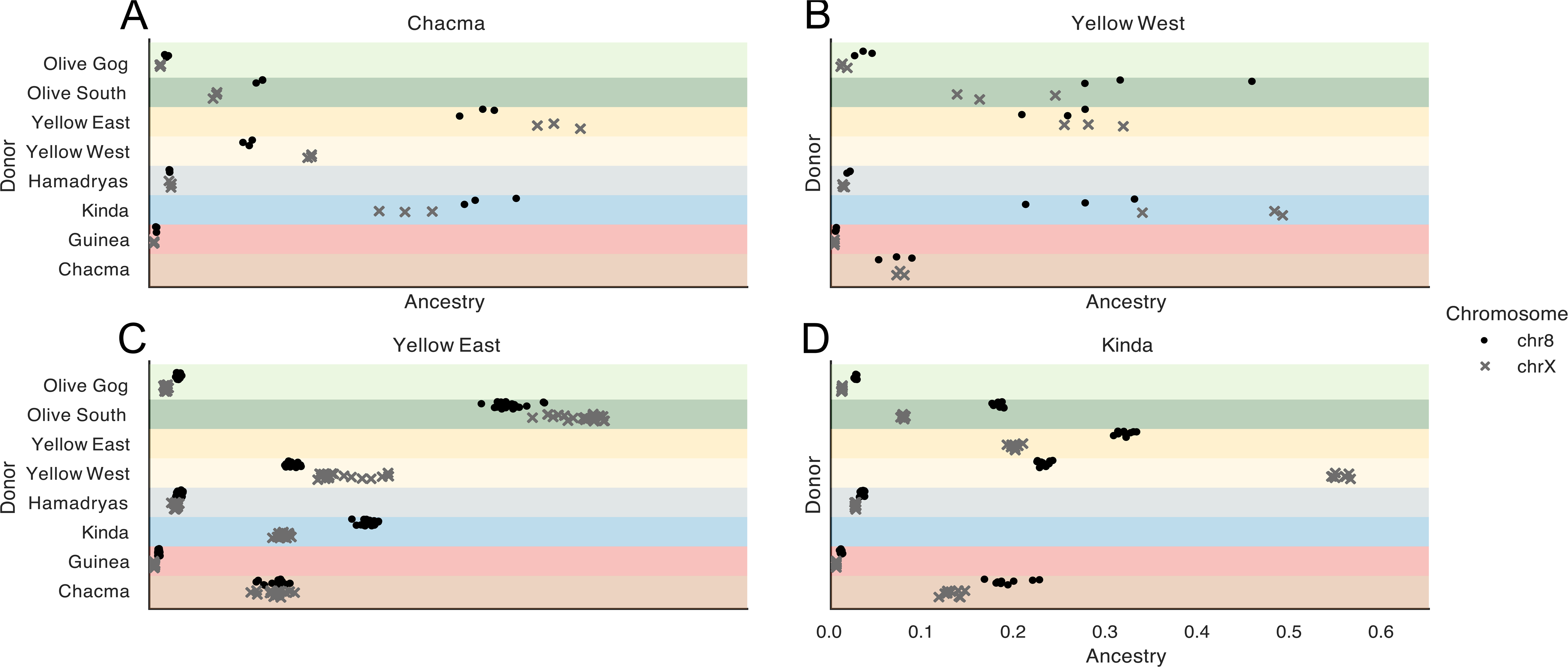
Differential ancestry profiles on the X-chromosome and an autosome. **(A**) Ancestry proportions of female chacma baboons. Each marker represents the fraction of total chromosome ancestry of one individual that is assigned to each of the remaining donor populations. Black dots and grey crosses represent ancestry proportions of chromosomes 8 and X, respectively. (**B**) Same as A but for female western yellow baboons. (**C**) Same as A but for female eastern yellow baboons. (**D**) Same as A but for female Kinda baboons. For additional profiles see figs. S10, S30 and S31.

In addition to patterns of shared ancestry among populations and species, we used two strategies to seek preliminary evidence for species-specific genetic adaptations in baboons. First, we used PLINK *(41)* to identify SNVs enriched in one species relative to all others (table S8). Genes containing possibly functional SNVs enriched in a given taxon were correlated with species phenotypes using Gene Ontology (GO) *(42)* terms and literature searches. We also used OmegaPlus *(43)* to test those gene regions for evidence of selective sweeps. Across all species, 1,342,371 SNVs met the criteria for being enriched in one particular species, including 4,337 missense and 76 stop gained SNVs (table S8). We next searched this list of candidates for genes annotated as influencing known traits of that species. Among them, SNV_1 (Table 1, fig. S32), a missense variant in Serine Protease 8 (*PRSS8*) has a 0.96 allele frequency (AF) in hamadryas baboons and a 0.02 AF in the geographically adjacent Gog olive baboons (absent in other species). *PRSS8* increases epithelial sodium channel activity and mediates sodium reabsorption through the kidneys *(44)*. *PRSS8* is under positive selection in the desert-adapted canyon mouse (*Peromyscus crinitus*) *(45)*, and hamadryas baboons inhabit the most arid environment of all baboons *(46)*. SNV_2 (Table 1, fig. S33) has a 1.0 AF in both hamadryas and Guinea baboons and is absent from other species. This is a missense variant in Neurexin 1 (*NRXN1*) which is associated with the GO term “social behavior”. *NRXN1* knockout mice exhibit changes in male aggression *(47)*. Guinea and hamadryas baboons differ from others in the genus in exhibiting a multi-level male-philopatric social organization with substantial male-male tolerance *(29, 48)*. This contrasts with the matrilineal, male-dispersing social organization typical and likely ancestral for the genus. This observation is compatible with the speculation that until “swamped” by males from olive and yellow baboon populations, male-philopatric “pre-Guinea” and “pre-hamadryas” baboon populations occupied the northern savanna-woodland belt and much of the East African savanna-woodland corridor *(33)*. SNV_3 (Table 1, fig. S34) has a 1.0 AF in Kinda baboons and a 0.05 AF in yellow baboons (western yellow baboons and Ruaha) and one Serengeti olive baboon. This is a missense variant in the pigmentation-associated Agouti Signaling Protein (*ASIP*). In mice, this gene affects melanin synthesis, shifting eumelanin production (black/brown hair) to phaeomelanin (red/yellow hair) *(49)*. Kinda baboons display several unique coat color traits, including a substantial proportion of infants with white natal coats *(16)*.

**Table 1.** Species enriched SNV statistics. Cluster and OmegaPlus statistics for the hamadryas and Guinea baboon shared SNV_2 are shown for hamadryas baboons. CADD and REVEL scores from human annotations predict functional impact of mutations (see Supplementary Materials).

| SNV ID | SNV | PLINK p value | Cluster Length (bp) | SNVs in Cluster | OmegaPlus (percentile) | CADD PHRED | REVEL |
| --- | --- | --- | --- | --- | --- | --- | --- |
| SNV_1 | 20:27347531:G:T | $1.40 \times 10^{-78}$ | 64,284 | 24 | 5.59 (1.8%) | 0.001 | 0.351 |
| SNV_2 | 13:49896439:G:C | $9.72 \times 10^{-101}$ | 126,701 | 96 | 11.01 (0.3%) | 7.266 | NA |
| SNV_3 | 10:30107617:T:C | $2.52 \times 10^{-69}$ | 39,912 | 58 | 4.92 (0.7%) | 19.140 | 0.080 |

In our second approach to functional variation, we searched for genomic regions of elevated differentiation between pairs of closely related species (for details see *(13)*). We asked whether regions with the strongest evidence of differentiation (windows in the top 0.1%) were enriched for genes with particular GO terms. Genomic regions most distinct between Kinda and yellow baboons were enriched for genes linked to skeletal development and morphogenesis (p-value adjusted for false discovery rate, p = 1.77×10^-4^); tables S9 to S11; fig. S35) including limb development (e.g., embryonic forelimb morphogenesis, adj. p = 0.02). This enrichment was driven by one region on chromosome 3 containing a HOXA gene cluster (fig. S36) and may influence the distinctively small size and gracile, long-limbed build of Kinda baboons *(16)*. Genes linked to male sexual differentiation were also increased in regions highly differentiated between Kinda and yellow baboons (adj. p = 0.0484), possibly related to the reduced sexual dimorphism in Kinda baboons *(50)*.

## DISCUSSION

Our expanded whole-genome dataset provides several novel insights into genetic reticulation and the evolutionary history of multiple local populations of baboons. Previous work showed that gene flow occurs among phenotypically and genetically distinct baboon species and pointed to nuclear swamping as a major contributing process. Our study extends and adds higher resolution to this picture, using genetic data to confirm hybrid zones that were previously suspected from field observation of phenotypic variation alone. We also identify the first local population (western Tanzanian yellow baboons) that has clear evidence for genetic contributions from three genetically distinct lineages.

While our results significantly extend our knowledge of baboon evolutionary history, some gaps remain. The richness of evolutionary detail to be derived from denser sampling is indicated by our results from East African populations. More materials are needed to document other regions with complex biogeographic and evolutionary history, including the olive-Guinea baboon interface in West Africa *(21)*, and regions of southern Africa where chacma baboons have experienced both ancient and recent periods of genetic divergence and reticulation *(39, 40)*. Other geographic regions, e.g., the northern savanna-woodland belt west of our Gog population have not been studied and would likely provide further information, especially regarding the origins and history of olive and Guinea baboons. Nevertheless, our dense sampling in East Africa clearly identifies new arenas of gene flow and documents the complexity of the evolutionary history of baboons in this region.

Our results lead to several substantive conclusions. With regard to methods, we find that while comparison of mtDNA and phenotypic variation are effective in detecting nuclear swamping, analyses comparing levels of shared ancestry across the X-chromosome to that across autosomes provides a more quantitative assessment of demographic processes and genetic history. Second, we conclude that Kinda baboons are not the product of a recent fusion event. Instead, they are more likely close to the basal ancestor of all extant baboons. Next, we find additional support for the prior observation that the primary separation of northern and southern baboon species is the result of dispersal from the south to the north, with Guinea baboons recognized as the most recent occupants of the leading edge of that dispersal. Despite the sharp gradient of phenotypes that is characteristic of baboon inter-species contact zones, gene flow distributes the introgressed alleles far from the regions of obvious hybridization. Last, we report that extant western yellow baboons carry genetic contributions from three genetically different baboon lineages.

The patterns of local, regional and species-level genetic structure in baboons are likely a valuable model for population structure in other primate clades that consist of multiple closely related species, such as African green monkeys (genus *Chlorocebus (51)*) and macaques (genus *Macaca (52)*). Clades in other mammalian orders are also revealing complex, often reticulated, evolutionary histories like those of baboons (e.g., polar bears *(53, 54)*, giraffes *(7)*, and deer *(55)*). The results for baboons also provide informative parallels and contrasts to the evolutionary differentiation and relationships among early human ancestors that arose, differentiated and admixed over a timespan remarkably similar to that of baboon cladogenesis *(56)*.

## MATERIALS AND METHODS SUMMARY

Extended materials and methods are available in the supplementary materials.

### Samples and DNA Sequencing

Blood samples from 225 baboons and two geladas were gathered in accordance with local regulations. Genomic DNA was extracted from blood and libraries were prepared for sequencing on the NovaSeq 6000 platform (Illumina).

### Variant Calling and Phasing

We used BWA-MEM to map reads to the Panu_3.0 baboon and the Mmul_10 rhesus assemblies. GATK was used to call variants following best practices. Panu_3.0 SNVs were phased using WhatsHap and SHAPEIT.

### Population Structure and Phylogenetic Analyses

Population structure based on SNVs was examined using PCA, ADMIXTURE, and fastSTRUCTURE. Phylogenetic trees based on autosomal and sex chromosome SNVs and Geneious assembled mitochondrial genomes were generated using IQ-TREE and visualized with FigTree. Polymorphic mobile elements were identified using DELLY and MELT. STRUCTURE and MELT were used to analyze population structure of L1 and *Alu* elements. PAUP was used to generate maximum parsimony trees from *Alu* and L1 elements. We used MSMC2 to infer baboon demographic history and population structure through time. Admixture graphs and F3 outgroup statistics were generated using ADMIXTOOLS 2.

### Inference of most recent coancestry along each chromosome

ChromoPainter was used to infer the most recent coancestry along chromosomes and fineSTRUCTURE was used to identify relationships between individuals based on their most recent coancestry. We used Globetrotter to compute p-values for a coancestry contribution from recent admixture.

### Functional Variation

Functional variation was examined using PLINK for association analyses, OmegaPlus for selective sweep identification, and differentiation-based scans for selection using windowed F_ST_ values.

## ACKNOWLEDGEMENTS

We thank all countries and their respective governmental and non-governmental institutions that supported sampling and sample analysis. Specifically, we thank the Government of the United Republic of Tanzania, Ministry for Education and Vocational Training, Commission for Science and Technology, Ministry for Natural Resources and Tourism, Ministry for Agriculture, Natural Resources, Livestock and Fisheries, Department of Forestry and Non-renewable Natural Resources, Tanzania Wildlife Authority, Tanzania Wildlife Research Institute, Tanzania National Parks (Yustina A. Kiwango, Rehema Kaitila, Inyasi A. V. Lejora), Ngorongoro Conservation Area Authority, Sokoine University of Agriculture (Rudovick R. Kazwala), National Institute for Medical Research (Clara C. Lubinza, Sayoki G. M. Mfinanga), Jane Goodall Institute (Iddi F. Lippende, D. Anthony Collins), and the Greater Mahale Ecosystem Research and Conservation Project (Alexander Piel, Fiona A. Stewart). For Zambia, we thank the Government of Zambia, the Zambia Wildlife Authority (Jack Chulu, Edwin Matokwani; Chilanga), the staff of Kafue National Park, and the Department of Veterinary and Livestock Development (Yona Sinkala, Lusaka). For Ethiopia, we thank the Government of Ethiopia, the Ethiopian Public Health Institute (Ebba Abate, Addis Ababa), the Guinea Work Eradication Program of The Carter Center (Jim Zingeser, Ernesto Ruiz-Tiben, Zerihun Tadesse, Addis Ababa) as well as James Else, Harry Marshall and the logistics team in Gog Woreda. For Senegal, we thank the Diréction des Parcs Nationaux and Ministère de l’Environnement et de la Protéction de la Nature de la République du Sénégal for permission to work in the Niokolo Koba National Park. We particularly thank the former conservateurs of the park Colonel Ousmane Kane and Commandant Mallé Gueye for their cooperation and logistical support during the study period, and all the staff and field assistants of the CRP Simenti, in particular Mustapha Faye, Armél Louis Nyafouna, Elhadji Dansokho, Lamine Diedhiou, Moustapha Dieng and Touradou Sonko for their support in the field.

## Author contributions

Conceptualization: KK-HF, TM-B, CR, JR; Data curation: EFS, RAH, LZ, MR, LFKK; Formal analysis: EFS, RAH, LZ, MR, JAW, JMS, MK, CF, LS, CMB, ASB, JB, KM, CR; Funding acquisition: KK-HF, TM-B, KM, CR, JR; Investigation: EFS, RAH, LZ, MR, LFKK, JAW, JMS, MK, CF, LS, CMB, ASB, JB, MHS, MAB, CJJ, KM, CR; Methodology: MK, CF, CMB, M-CG, SS, HD, MAB, SK, DZ, TM-B, KM, CR, JR; Project administration: KM, CR, JR; Resources/Sample acquisition: CMB, ASB, JEP-C, FS, KLC, ISC, JDK, JF, CJJ, SK, DZ, CR, JR; Supervision: TM-B, KM, CR, JR; Visualization: EFS, RAH, LZ, JAW, JMS, MK, CF, CMB, DZ, KM, CR; Writing – original draft: EFS, RAH, LZ, JAW, MK, CF, CMB, CJJ, DZ, TM-B, KM, CR, JR; Writing – review & editing: all authors

## Data and materials availability

The sequencing data used in these analyses are available through the Short Read Archive under BioProject accession PRJEB49549. Additional data are available in the Supplementary Materials.

## SUPPLEMENTARY MATERIALS

Materials and Methods

Supplementary Text S1 to S5

Figs. S1 to S36

Tables S1 to S11

References and Notes *(57–114)*

