## Supplementary Material for "Genome-wide coancestry reveals details of ancient and recent male-driven reticulation in baboons"

**This PDF file includes:**

Materials and Methods
Supplementary Text S1 to S5
Figs. S1 to S36
Tables S3 to S5, S8

**Other Supplementary Materials for this manuscript include the following:**

Tables S1, S2, S6, S7, S9 to S11 (.xlsx)

#### Materials and Methods

##### Sampling

All samples were gathered in accordance with and abiding by local laws and regulations. Most samples from Tanzania originated from studies on *Treponema pallidum* infection in nonhuman primates (*57–59*). For this aspect of sampling, baboons were chemically immobilized using 10.0 mg ketamine/kg body mass in combination with 0.2 mg/kg medetomidine. Anesthetics were intramuscularly injected using a cold-gas immobilization rifle (MOD JM) and appropriate projectiles. Immobilized baboons were continuously observed for vital parameters such as respiration, pulse frequency, and internal body temperature. We collected whole blood from the femoral vein using an S-Monovette EDTA closed blood collection system (Sarstedt) mounted with a 20G needle. After sedimentation, buffy coat was collected and samples were frozen at -80°C until further use. We allowed animals to recover under close supervision. All procedures were performed by trained veterinarians and animals were handled applying highest animal welfare standards.

Other samples from Tanzania taken between 1984 and 1986 were collected by remote distance immobilization or capture followed by anesthesia in custom-designed falling door traps (*60*). Baboons were administered ketamine (10mg/kg body mass) and blood was drawn from the femoral vein into Vacutainer tubes with EDTA anticoagulant. Animals were released back to their social groups after they fully recovered from anesthesia. Samples from Zambia and Ethiopia were collected through capture in falling door traps. Baboons in Zambia and Ethiopia were immobilized with ketamine (10mg/kg body weight) and blood was drawn from the femoral vein using EDTA vacutainer tubes. Baboons in Senegal belong to a habituated study group. Anesthetics were, therefore, applied directly using a blowpipe. Blood was collected as described previously.

##### Dataset

In this study, we investigated a total of 225 baboon individuals and two geladas, the latter were used for outgroup purposes. Whole-genome sequence (WGS) data for 217 baboons and one gelada have been newly generated in this study, while data from another gelada and eight geo-referenced baboons were previously published (*23*). Baboon samples were derived from 19 geographic locations with 1 to 38 samples per location (n=1: Issa Valley; n=2: Aberdare and Katavi; n=3: Selous; n=4: Arusha and Dendro Park; n=5: Udzungwa; n=6: Ngorongoro and Ruaha; n=7: Mahale and Tarangire; n=12: Niokolo Koba; n=14: Serengeti; n=17: Gombe; n=19: Lake Manyara; n=25: Gog; n=26: Filoha; n=27: Chunga; n=38: Mikumi; average 11.8; table S2).

##### DNA extraction and sequencing

Genomic DNA was processed and sequenced by two different groups. For 116 of the samples, DNA from blood samples was extracted with the QIAamp DNA Mini Kit (Qiagen). The short-insert paired-end libraries for WGS were prepared with PCR-free protocol using KAPA HyperPrep kit (Roche), with some modifications. In short, genomic DNA was sheared on a Covaris™ LE220-Plus (Covaris) to reach the average fragment size of ~300bp. The fragmented

DNA was size-selected for the fragment size of 220-550bp with AMPure XP beads (Agencourt, Beckman Coulter). The size selected genomic DNA fragments were end-repaired, adenylated, and Illumina platform compatible adaptors with unique dual indexes and unique molecular identifiers (UMI, integrated DNA Technologies) were ligated. The libraries were quality controlled on an Agilent 2100 Bioanalyzer with the DNA 7500 assay (Agilent) for size and quantified by Kapa Library Quantification Kit for Illumina platforms (Roche). Library with final molarity below 3nM underwent PCR amplification of 6 - 10 cycles using KAPA Library Amp Primer Mix (Roche) and KAPA HiFi HotStart ReadyMix PCR Kit (Roche). The libraries were sequenced on NovaSeq6000 (Illumina) in paired-end mode with a read length of 2x151 bp following the manufacturer's protocol for dual indexing.

For the remainder of samples (primarily legacy samples collected in Ethiopia, Tanzania, and Zambia several years ago), DNA was extracted either through standard phenol-chloroform extraction procedures or by using QIAamp DNA extraction kits (Qiagen) or GenFind3 (Beckman Coulter). Following quality control to check purity and DNA concentration, we produced PCR-free KAPA Hyper paired-end sequencing libraries. Total genomic DNA (500-750 nanograms) was sheared to fragments 200-600 bp in length using a Covaris E220 system (96-well format) and purified using AMPure XP beads. The sheared DNA was then subjected to double size selection with two different ratios of AMPure XP beads. This is followed by DNA end-repair and 3' adenylation before ligation of barcoded adapters. The quality of library preparations was evaluated by fragment analysis using a Fragment Analyzer (Advanced Analytical Technologies) and tested via qPCR assay using KAPA Library Quantification Kit and their SYBER FAST qPCR Master Mix. These whole-genome libraries were then sequenced on Illumina NovaSeq 6000 instruments using S4 reagents, producing 2 x 150 bp paired-end reads. The final concentration of the libraries loaded on the flowcells is 400-450 pM. Briefly, the libraries are diluted in an elution buffer and denatured in sodium hydroxide. The denatured libraries are pooled to maximize use of flowcells and loaded into each lane of the S4 flowcell using NovaSeq Xp Flowcell Dock. Each lane also includes ~1% mixture of PhiX control library for run quality control.

#### Variant calling and phasing

We used BWA-MEM version 0.7.12 (*61*) to map sequence reads to the Panu_3.0 baboon genome assembly GenBank Accession GCA_000264685.2 and the Mmul_10 rhesus assembly GenBank Accession GCA_003339765.3. To identify reads potentially originating from a single fragment of DNA and mark them in the binary alignment map (BAM) files, we used Picard MarkDuplicates version 1.128 of the Picard package (https://broadinstitute.github.io/picard/). Variants were then called using the Genome Analysis Toolkit (GATK) version 4.1.6.0 (*62*) following best practices (https://gatk.broadinstitute.org/hc/en-us/articles/360035535932-Germline-short-variant-discovery-SNPs-Indels-) and variant call format (VCF) files were generated. The hard filters suggested by the developers of GATK were applied to the SNVs and indels (https://software.broadinstitute.org/gatk/documentation/article?id=11097) and all failing variants were removed. We then used GATK VariantAnnotator to annotate SNVs applying AlleleBalance. Variants with an allelic balance for heterozygous calls (ABHet=ref/(ref+alt)) ABHet < 0.2 or ABHet > 0.8 were removed. Variant Effect Predictor (VEP) version 102 was used to annotate the variants in the VCF files based on combined Ensembl and RefSeq gene annotations.

For SNV calls on Panu_3.0, WhatsHap version 1.1 (*63*) with default settings was used to perform read-based phasing of SNVs individually for each sample. SHAPEIT version 4.2.1 (*64*) with --use-PS 0.0001 was applied to the WhatsHap phasing results to generate longer phased haplotypes.

#### Principal component analysis (PCA)

We performed PCAs based on biallelic SNV data (autosomes and X-chromosome: all 225 individuals, mapped to the rhesus macaque (Mmul_10) genome, filtering parameters --maf 0.05, Y-chromosome: all 122 males, mapped to the Mmul_10 genome, --maf < 0.05, all heterozygous sites removed) using the smartPCA program from the EIGENSOFT package version 6.1.4 (*65*) with default settings. The genotype input files were transformed by PLINK version 1.90 (*41*) and convertf program (EIGENSOFT package) from the population VCF files. To determine the significance level of different principal components, we performed the Tracy-Widom test using the twstats program (EIGENSOFT package). Two- and three-dimensional PCA plots were generated by R based on the evec results.

#### ADMIXTURE analysis of autosomal SNVs

We inferred the population genetic structure using a maximum likelihood (ML) estimation of individual ancestries from autosomal SNV genotype data (input files were generated by vcftools and PLINK with filtering parameters --geno 0.05 --maf 0.05 --hwe 0.0001) in ADMIXTURE version 1.3.0 (*66*) using all 225 baboon individuals and investigating population structuring into *K* = 2-10 clusters. The best number for *K* was determined using cross-validation error methods (-cv commands, the lowest cross-validation error was chosen as the best *K* value). The cross-validation error for different *Ks* were *K*=2: 0.39229, *K*=3: 0.36916, *K*=4: 0.34398, *K*=5: 0.34548, *K*=6: 0.33424, *K*=7: 0.32151, *K*=8: 0.33457, *K*=9: 0.32455, and *K*=10: 0.33261. Accordingly, *K*=7 represents the most suitable population structure. We note that the cross-validation errors for several *K* values in our study are highly similar and do not show a clear unimodal distribution, suggesting high complexity in our dataset and similarly likely numbers of clusters. The analysis using *K*=8 rejoins hamadryas and Guinea baboons, first split at *K*=6, into the same cluster. As this could indicate that the likelihood optimization finds a local maximum, we redid the analysis using different random seeds and replicated the analysis using fastSTRUCTURE. However, although each analysis reproduces the same clustering, this could reflect a shared difficulty in finding a global optimum. For this reason we report the clustering for, *K*=7 in the main text.

#### Autosomal and X- and Y-chromosomal phylogenies

To reconstruct phylogenetic trees, we first used all high-quality SNV datasets (see PCA) for autosomes and X- and Y-chromosomes with gelada as outgroup (2 geladas for autosomes and X-chromosome, 1 male gelada for Y-chromosome). For the X- and Y-chromosome phylogenies, SNVs of all 225 (and both geladas) and 122 male baboon individuals (and one gelada), respectively were aligned across the rhesus macaque (Mmul_10) X- and Y-chromosome assemblies to generate input files in Phylip format by Perl scripts. For the autosomal phylogeny, we aligned all 225 baboon individuals and both geladas using all autosomal SNVs, then used window-based methods to extract 5% of SNVs randomly as the input for the phylogenetic reconstruction. To ensure the accuracy of the obtained tree topology, the phylogenetic tree reconstruction was repeated with another randomly selected 5% SNV dataset. ML trees were generated in IQ-TREE version 2.1.3 (*67*) with 1000 ultrafast bootstrap replicates (*68*) and 16 threads to obtain statistical node support, and applying the respective best-fit substitution model as automatically calculated with ModelFinder (*69*) in IQ-TREE according to the Bayesian information criterion. FigTree version 1.4.0 (http://tree.bio.ed.ac.uk/software/figtree/) was used to visualize the SNV-based trees.

Next, we used sequence data to reconstruct phylogenetic trees. For the Y-chromosome, we used all 122 baboon males and a male gelada. Haplotype sequences were extracted with a perl script from population VCF files and transferred to fasta format using FastaAlternateReferenceMaker of the GATK package version 4.1.8.1, then all sequences were aligned to the Y-chromosome of rhesus macaque (Mmul_10). A ML tree was generated with IQ-TREE with 1000 ultrafast bootstrap replications (-bb 1000) and the best-fit substitution model as described above. To obtain divergence times, we re-ran IQ-TREE using the phylogenetic dating option (*70*). Therefore, we applied a relaxed clock model and constrained the split between *Papio* and *Theropithecus* at 4.2 million years ago (Mya) based on the oldest known *Theropithecus* fossil (*71, 72*). To obtain confidence intervals we resampled branch lengths 100 times and used the default setting of 0.2 for the standard deviation of the lognormal distribution. The trees were visualized and edited with FigTree. For autosomes and the X-chromosome, we applied a window-based approach using 50kb and 500kb windows and one individual per species and location, selected based on results from the ADMIXTURE analysis (olive baboons from Ngorongoro and Lake Manyara combined). Gelada was used as the outgroup. To obtain putatively neutral regions, we masked exons and respective 10kb flanking regions as well as repeat regions according to the annotation files (rheMac10.ensGene.gtf). We retained only windows with at least 20% unmasked sequence data, resulting in 28,700 windows with average unmasked 20,646bp (range: 10,001-49,180bp) and 3129 windows with average unmasked 175,845bp (range: 100,003-338,873bp) for autosomes, and 1553 windows with average unmasked 17,954bp (range: 10,001-38,438bp) and 165 windows with average unmasked 154,953bp (range: 100,654-244,170bp) for the X-chromosome. We then generated ML trees in IQ-TREE with 1000 ultrafast bootstrap replications and the respective best-fit substitution models as described above. A species tree under the multi-species coalescent model was generated in ASTRAL version 5.7.7 (*36*) using standard settings and visualized in FigTree. To obtain divergence times, we re-ran IQ-TREE using the phylogenetic dating option, applying a relaxed clock model and constraining the split between *Papio* and *Theropithecus* at 4.2 Mya. Generated window-trees were visualized in DensiTree version 2.6.6 of the BEAST package version 2.6.0 (*73*).

#### Mitochondrial genome assembly and phylogeny

Complete mitochondrial genomes were generated via mapping the cleaned sequence reads to the following baboon reference mitogenomes: *P. anubis*: NC_020006, *P. hamadryas*: JX946201, *P. kindae*: NC_020008, *P. papio*: NC_020009, *P. ursinus*: NC_020010, *P. cynocephalus*: JX946200, MT279060, or MT279069. Mapped sequence reads were then re-mapped to the phylogenetically closest available baboon mitogenome with the Geneious assembler of the

Geneious package version 11.1.3 (https://www.geneious.com) using default settings. Generated mitogenomes were manually checked and annotated in Geneious.

For phylogenetic tree reconstructions, we expanded our dataset with 21 additional baboon mitochondrial genomes (*20, 21*); gelada was used as outgroup. A total of 247 mitochondrial genomes were aligned with Muscle version 3.8.31 (*74*) in AliView version 1.18 (*75*) and identical haplotypes (table S7) were subsequently removed. A ML tree, treating the mitogenome as a single partition, was generated with IQ-TREE using 1000 ultrafast bootstrap replications and the best-fit substitution model as determined by ModelFinder. A dated phylogeny was calculated as described above, constraining the split between *Papio* and *Theropithecus* at 4.2 Mya. The trees were visualized and edited with FigTree.

**Detection of polymorphic mobile element insertions (MEIs)**

Two strategies were implemented to identify mobile element insertions, an SV caller, DELLY version 0.8.1 (*76*), and MELT (Mobile Element Locator Tool) version 2.2.2 (*77*). A local installation of RepeatMasker (http://www.repeatmasker.org) was used to ascertain LINE-1 (L1) MEIs from the Panu_3.0 genome assembly for an olive baboon. L1 elements >5500bp were considered full-length. RepeatMasker-based coordinates for 3884 L1 elements provided the input sequence reads for DELLY genotypes. A local installation of MELT was used to ascertain full-length *Alu* elements from WGS. Full-length *Alu* elements are defined as possessing a start position no less than 4 bp and an end position not shorter than 267 bp. BAM files for all available samples exceeded 11TB of data. Due to our computational server limitations, a subset of 43 individuals representing each population were included for the initial MELT analysis. Read files were aligned to the outgroup rhesus macaque assembly (Mmul_10) to determine lineage-specific and polymorphic insertions.

**STRUCTURE analysis for L1 elements**

Analyses of population structure were performed using STRUCTURE version 2.3.4 (*78*). Binomial genotype data for 261 L1 elements and 222 baboons were arranged in an excel spreadsheet using consecutive columns as "1, 1" for an insertion that is fixed present in an individual, "1, 0" for an insertion that is heterozygous in an individual, and "0, 0" for the absence of the L1 element in an individual. Missing genotypes were given a value "-9". This file was converted to a tab-delimited text file suitable for upload into STRUCTURE. All analyses were performed under the admixture model which assumes that individuals may have mixed ancestry. No information about sample origin was included. To determine the number of population clusters ($K$), our initial simulations tested $K$ = 2-10. These analyses used 20,000 burn-in, followed by 200,000 Markov chain Monte Carlo (MCMC) iterations with three replicates for each $K$ value. The Simulation Summary data were exported and LnP(D) values were analyzed for the most likely value of $K$. The output run files were then analyzed using the delta-K method (*79*) implemented in the program Structure Harvester (*80*) to determine the most appropriate $K$ value. Following these procedures, the most likely value of $K$ was determined to be eight and was graphed in Microsoft Excel (fig. S13). The LnP(D) value for different $K$s are $K$=2: -40165, $K$=3: -37022, $K$=4: -35092, $K$=5: -33816, $K$=6: -33984, $K$=7: -38757, $K$=8: -33229, $K$=9: -33527, $K$=10: -33936.

#### <u>MELT analyses of *Alu* and L1 elements</u>

MELT was used to detect and genotype lineage-specific baboon MEIs from a subset of 43 individuals (figs. S22 and S23). Briefly, BAM files of baboon sequencing reads aligned to the rhesus macaque genome assembly (Mmul_10) served as input for MELT analysis to detect and genotype MEIs present in the selected baboon samples, but absent in the rhesus genome. The *Alu*Y family and associated subfamilies are only active in Old World monkeys and apes. Therefore, for *Alu* insertions, the *Alu*Y consensus sequence was used to detect *Alu* insertions within the BAM files, allowing for 6 mutations per 100 bp. The MELT output for the *Alu* analysis was filtered to exclude *Alu*J and *Alu*S families, include only those insertions with evidence for target site duplications, and to exclusively analyze full-length insertions.

MELT detection of LINE insertions was performed with consensus sequences for L1PA6, L1RS25, and L1RS10, derived from full-length insertions obtained from the olive baboon (Panu_3.0) genome assembly. Full-length LINE insertions were defined as having a length of >= 5500 bp. LINE consensus sequences were derived using the alignAndCallConsensus.pl program (*81*). For the detection of L1PA6, L1RS25, and L1RS10 sequences, respectively, eight mutations per 100 bp in the BAM files were permitted. The aforementioned L1 consensus sequences were chosen for MELT analysis to represent a range of ages within the baboon lineage. The L1PA6 subfamily was active after the split of Catarrhini from Platyrrhini but shared between Cercopithecoidea and Hominoidea, and was thus chosen as the oldest L1 subfamily for MELT analysis to determine ancestral relationships among baboon populations. The L1RS subfamilies are derived from L1PA5, a younger family than L1PA6. Based on the full-length L1RS distribution in both the rhesus and baboon genomes, L1RS25 displayed a spike in the number of insertions, and could therefore be indicative of a speciation event within Cercopithecoidea (*82*). A previous analysis has indicated that L1RS2 and L1RS1 are the youngest elements within the rhesus macaque genome (*83*). In addition, a speciation event, and associated rhesus macaque lineage-specificity of the L1RS2 group, is supported by a large increase in L1RS2 insertions in the rhesus macaque assembly, but not the baboon genome assembly. Therefore, L1RS10 is likely the youngest defined subfamily in the baboon genome, as L1RS2 and L1RS1 are expected to be rhesus-specific. For both *Alu* and L1, custom python scripts were used to reformat the MELT output into a nexus format for PAUP analysis.

#### <u>Maximum-parsimony tree reconstruction for *Alu* and L1 elements</u>

A heuristics search was performed using PAUP* version 4.0a169 (*84*). Because it is assumed that the absence of a retrotransposable element is the ancestral state of each locus, Dollo's law of irreversibility was used in all parsimony analyses with all loci set to Dollo.up. From the L1 genotype data derived from the DELLY output, a nexus file was generated using a custom python script. A deletion compared to the reference genome (1/1) in the DELLY output was scored as a "0", while a shared insertion (0/0) was scored as a "1". Ambiguous loci were indicated with "?". One thousand bootstrap replicates were performed with a maximum tree space set to 100. An artificial individual with all loci set to "0" was used as the outgroup with which to root the phylogenetic tree generated by FigTree version 1.4.4 (<u>http://tree.bio.ed.ac.uk/software/figtree/</u>). Several similar maximum parsimony analyses were performed using MELT generated *Alu* genotype data. These included 42 baboons representing each wild population and one gelada. One thousand to 10,000 bootstrap replicates were performed for three independent nexus files. First, 293,506 *Alu* elements distributed across all chromosomes. These data were filtered for full-length *Alu* insertions from subfamily *Alu*Y only resulting in 246,219 insertions (fig. S22). For the L1PA6 data, 35,913 L1PA6 insertions are distributed across all chromosomes (fig. S23). A male Y-chromosome *Alu*-based phylogeny was generated using 355 elements and 24 male baboons (fig. S9). Using MELT to align all samples against rhesus macaque (Mmul_10) places equal evolutionary distance between each sample and the reference outgroup, thus minimizing directional bias, whereas the L1 DELLY analysis was subject to sampling bias towards olive baboons (Panu_3.0). Although the phylogenetic trees generated from MELT data had robust bootstrap support and were quite consistent with sampling localities, the homoplasy index parsimony tree score was still about 0.85 or higher, indicating extensive reticulation in these populations.

#### Multiple Sequentially Markovian Coalescent (MSMC) analyses

We used MSMC2 version 2.1.1 (*85*) to infer baboon demographic history and population structure through time. Using BWA-mem mappings to Panu_3.0 autosomes, GATK version 4.1.6.0 GenotypeGVCFs was run to include all sites variant or not using the -all-sites flag. The resulting SNVs were phased using WhatsHap version 1.1 and SHAPEIT version 4.2.1. The vcfAllSiteParser.py script from msmc-tools was used to generate VCF and mask.bed files from the phased GATK VCF. The script generate_multihetsep.py was used to generate files for input to MSMC2 using four phased individuals from each population or species. MSMC2 was run using default settings. MSMC2 plots were generated using a mutation rate of 0.9x10-8 and a generation of 11 years (*23*).

#### Admixture graphs

We applied an individual filtering approach to obtain a dataset for calculating population statistics like f3 or f4 and for creating admixture graphs (*86*). We first obtained only bi-allelic segregating sites on the autosomes from genotypes mapped to the rhesus macaque (Mmul_10) genome, removing sites where no genotype was obtained for any individual. We then obtained the genotype, sequencing depth, and alternative depth fields using BCFTools (*87*) to remove positions where sequencing coverage was below 1/3-fold or above 2-fold the mode coverage of each individual, and positions where the minor allele was observed in less than 20% of the reads (allele imbalance). Given the better quality, here we used only newly sequenced individuals. Genotype data were converted to the “eigenstrat” format required for ADMIXTOOLS version 5.1. We first created admixture graphs using the R package admixtools 2 (https://github.com/uqrmaie1/admixtools). We calculated f2-statistics using the function extract_f2. We then used *find_graphs* with a specified number of admixture edges from 1 to 10 to determine the best fitting graph, defining gelada as the outgroup. We obtained the likelihood scores for each graph and calculated the out-of-sample scores, as described in the package vignette. Admixture graph topologies were also studied using f4-statistics, using the previously described admixturegraph package (*88*), to model the fit of all possible f4-statistics between the tested populations (*i.e.* all configurations of four populations). These statistics were obtained using the admixr package (*89*). The possible space of graphs with one admixture edge was explored using the function *add_an_admixture2*.

#### Population structure using F3 statistics

F3 outgroup statistics were obtained with ADMIXTOOLS (*86*) to estimate shared drift between different baboon populations, here defined as the sampling localities. We ran qp3Pop using gelada as the outgroup and visualized the shared drift in a pairwise matrix of the F3 values, plotted in R.

#### Inference of most recent coancestry along each chromosome - ChromoPainter

Chromosome painting requires a phased genome, as well as an estimate of recombination rate between every variable position. The VCF files for every individual were phased using SHAPEIT version 4.2.1 as described in the "Variant Calling and Phasing" section. For the autosomes, every individual was used, while the female X-chromosome was phased using only other females as males are haploid on the X-chromosome. The input file format needed for ChromoPainter version 2 (*30*) is a phase file, so the phased VCF was transformed using "plink --vcf with the commands --recode12 --const-fid 0" and a companion script supplied by ChromoPainter (plink2chromopainter).

We created a recombination map based on 20 female olive baboons from Tanzania using ldhat with standard interval settings. Choosing olive females for the analysis, we first excluded individuals with large runs of homozygosity to avoid biases from recent inbreeding and subsequently picked the 20 females with highest heterozygosity. ldhat estimates rho ($4*N_e*r$), which is scaled by the effective population size. Since ChromoPainter requires a map with Morgans/Base, we therefore need an estimate of the effective population size ($N_e$). We ran SMC++ on the same 20 olive baboons used to create the recombination map per chromosome. When calculating the harmonic mean of the $N_e$ until the most recent bottleneck, population sizes of 4500-7500 were found on the autosomes. Meanwhile, the harmonic mean of effective population size for the last $N_e$ generations was 12,000-17,000. A recent bottleneck will tend to reduce rho estimates, so an intermediate $N_e$ between these two extremes (8000) was used for the recombination map of the autosomes, and 6000 for chromosome X, as chromosome X in both measures was lower than the autosomes. This results in cM/Mb rates of 0.9 cM/Mb to 1.2 cM/Mb on the chromosomes, except for chromosome 19, which had a mean cM/Mb of 2.05. This rate is slightly lower than in humans but higher than in macaques. The recombination map was also transformed by use of a companion script supplied by ChromoPainter (convertrecfile).

ChromoPainter uses a hidden Markov model (HMM), in which one sample (the "recipient") is modeled as a mosaic of other individuals (the "donors"). As part of an expectation-maximization step, ChromoPainter estimates the mutation rate and the switch rate between chromosomal segments contributed from different donors. For ChromoPainter, these parameters are regarded as nuisance parameters and only used to optimize the fit of the underlying model. The analysis thus performs its own scaling of the supplied recombination map. The model parameters and the phased variant data are then used to identify the most likely donor at each position along each chromosome. The alternating donors along chromosomes each represent the most recent common ancestry among all donor individuals.

#### Clustering of individuals – fineSTRUCTURE

We ran fineSTRUCTURE version 4.1.1 (*30*) on the full autosomal dataset to identify relationships between individuals based on their most recent ancestry. To achieve this, fineSTRUCTURE analyzes chromosome paintings (see above) where all individuals can be donors to every other individual. It then uses MCMC sampling of possible clusters and creates a tree based on mean coincidence of pairs of individuals. fineSTRUCTURE recaptures all sampling locations, and also shows that all of the locations, even though they are of the same species, are genetically distinct. The only exception is locations with only 1-2 samples, which are grouped with samples from the most related population because no donors from their own population are available. The sampling locations in western Tanzania, which originally have been assumed to be yellow baboons, are identified as closer to Kinda and chacma baboon populations than to eastern yellow baboon populations. An especially noteworthy sample from western Tanzania is the one from Issa Valley, which seems to be a recent hybrid between the resident baboons in this location and invading olive baboons. Kinda baboons from Zambia cluster as one group, although this group is more heterogeneous than the groups representing the other species. There is a strong spatial component in the larger fineSTRUCTURE clusters, with olive baboons from Ethiopia clustering with hamadryas/Guinea baboons before they cluster with southern olive baboons, as well as the western yellow baboons clustering with Kinda and chacma baboons. The north/south split of the six species is also supported by the fineSTRUCTURE analysis. Fig. S4 depicts the fineSTRUCTURE results with a reduced heatmap ceiling.

The ChromoPainter results for each individual, produced for fineSTRUCTURE, can be summarized by the number of segments contributed by each donor individual. This relative coancestry is represented as a row in the coancestry matrix (Fig. 2B). PCA on this matrix (Fig. 2C). reveals the north/south along PC1, while PC2 distinguishes between the populations after the north/south split.

#### Population coancestry - Globetrotter

We ran Globetrotter (version 1) on the autosomal data set. Due to computational limitations of Globetrotter, filtering through PLINK is employed so that there are between 221,442 and 79,283 SNVs per chromosome (This is still more data than was used in analyzing human populations in the original Globetrotter publication (*38*). The Globetrotter analysis requires two sets of chromosome paintings as input: One set allowing for copying between all groups of individuals as for fineSTRUCTURE, and one set in which the focal group only can copy from other groups. We ran two Globetrotter analyses on the eight and 14 groups described in the main text. For each group, we produced 100 bootstraps using Globetrotter to compute p-values for a coancestry contribution from recent admixture. For computational efficiency, we used the newest implementation of Globetrotter, fastGlobetrotter (version 1) (*90*), which implements the same analysis, but with computational optimizations. The results reported uses the "null individual" functionality of Globetrotter, which corrects for changes in historical population size.

#### Contrasting ancestry on chromosome X and an autosome

To investigate the genetic evidence of sex-biased admixture, we contrast the ancestry proportions along the X chromosome and the similar-sized chromosome 8. We further use only females, as they are diploid on both chromosomes. We quantify ancestry as the sequence proportion that each chromosome derives from a donor group in the ChromoPainter analysis. The ancestral relationship of baboons is subject to incomplete lineage sorting (ILS) because of the rapid succession of speciation events producing the six species. The X-chromosome, hemizygous in males, has a smaller effective population size and is thus subject to less ILS. The prior expectation is therefore that the X-chromosome ancestry adheres more closely to the species tree if no admixture has occurred. Most importantly, however, admixture by hemizygous males introduces disproportionately more autosomal content than X-chromosomes. This is revealed as ancestry contributions on the autosomes not similarly reflected on the X-chromosome in which case the X-chromosome ancestry better represents the species relationship prior to admixture.

We ran ChromoPainter on chromosome X and the similar-sized chromosome 8 for all females in the dataset. We exclude males from this analysis to avoid any biases related to phasing of the two chromosomes and to ensure that the numbers of possible donors are the same for both chromosomes. No data was filtered in this analysis. For each phased chromosome, 10 paintings were generated using iterbi sampling. For each individual, inferred ancestry is assigned to either Olive Gog (Ethiopia), Olive south (Kenya and Tanzania), Yellow west, Yellow east, Hamadryas, Kinda, Guinea or Chacma. Separate ancestry proportions for chromosomes X and 8 were calculated by summing the total length of ancestry from each of the eight donor groups and dividing by the sum of all inferred ancestry. The significance of differences in ancestry contributions between chromosomes X and 8 across individuals was evaluated using a paired t-test.

Given the amount of independent genealogical data for each chromosome, chromosome 8 is expected to faithfully represent all autosomes. We repeated the analysis using chromosome 3 (fig. S31) and found conclusions to be consistent with chromosome 8.

#### Functional Variation

##### <u>Genetic association analyses of species-specific functional variants</u>

PLINK version 1.90b2t (*41*) genome-wide case/control association analyses were performed with each target species as the case and all the other species as controls. SNVs with a p-value < $5x10^{-8}$, a threshold commonly used for genome-wide association significance (*91*), were filtered for those with an allele frequency > 0.9 in the target species and allele frequency < 0.1 in the other species generating a list of species enriched SNVs. The species enriched SNVs were clustered using BEDTools version 2.26.0-19-g6bf23c4 (*92*) with a window of 10kb. OmegaPlus version 3.0.3 (*43*) was used to identify selective sweeps in each species using the parameters -grid 100000 -minwin 10000 -maxwin 100000. Identified species enriched missense SNVs were then examined in the context of clusters and OmegaPlus window scores. SNVs were reciprocally lifted over to the orthologous position on the human hg38 assembly using Picard version 2.18.25 LiftoverVcf. The lifted over SNVs were then scored using CADD version 1.6 (*93*) and REVEL version 1.3 (*94*). Species enriched missense SNVs were examined for relevance to species phenotypes using Gene Ontology (GO) terms as reported by the Database for Annotation, Visualization and Integrated Discovery (DAVID) (*95*) and literature searches.

**Differentiation-based scans for selection**

Differences in allelic frequency between closely related species at a particular genomic region may indicate the region has been the target of positive selection in one or both lineages. To identify such putative targets of selection, we searched the genomes for outlier regions of elevated differentiation between pairs of closely related species. We compared allele frequencies between Kinda versus yellow baboons, olive versus hamadryas baboons, and olive versus Guinea baboons, restricting our analyses to pairs of closely related species with higher sample sizes (n>12). Additionally, given the deep divergence identified within yellow baboons, we additionally examined genome-wide differentiation metrics for the eastern (Mikumi, Ruaha, Selous, Udzungwa) versus western (Mahale, Katavi, Issa Valley) yellow baboons.

To assess differentiation between populations, SNVs were included if, in any taxon pair, the minor allele frequency (MAF) was greater than 0.05. We computed $F_{ST}$ (*96*) in sliding windows of size 100 kb across the genome, stepping 50 kb each time (VCFtools) (*97*). We considered a genomic window to be an outlier if the value of $F_{ST}$ of SNVs in the window was in the top 0.1% of the distribution across the genome for that taxon pair (*e.g.*, empirical p-value < 0.001). We intersected top windows with annotated genes to generate a list of genes in outlier $F_{ST}$ windows for each taxon pair. For all comparisons, we tested whether the regions with the strongest evidence of differentiation (windows in top 0.1%) were enriched for genes with particular functional annotations (GO terms; (*42*)). For this functional enrichment, we assessed significance using a hypergeometric test as implemented in g:Profiler (*98*). We corrected for multiple testing by controlling for Benjamini-Hochberg False Discovery Rate (FDR), and adjusted p-values are reported for the functional enrichments.

### Supplementary Text

#### S1: Rationale for baboon taxonomy and nomenclature

Since the 1960s, various taxonomic schemes have been proposed for the genus *Papio* (reviewed in (*31, 99*)), and the only conclusions that are universally accepted are that i) the complexity of phenotypic variation and population genetic structure (including potential or actual gene flow among phenotypically distinct populations) observed among baboons defies simple Linnaean binomial nomenclature, and ii) there is discordance between the relationships among populations based on external phenotype and equivalent relationships based on genetic (mtDNA or nuclear) similarities (*20, 99*). Much of the debate about how to express baboon diversity taxonomically arises from disagreements about the conception and definition of species in general, and the criteria for recognizing species taxa (for reviews see (*20, 26, 31, 99*)). In the case of the baboons, the relevant biological facts, namely the broad picture of intra-generic diversity and inter-populational gene flow, are not seriously disputed. As briefly described in the main text, at least six major geographical forms are distinguishable, differing consistently in pelage, body size, social behavior, social systems and other phenotypic characters, with only minor differences among subpopulations within them. The geographic distributions of the major forms do not overlap but often adjoin. At their boundaries, interbreeding and backcrossing form narrow zones occupied by hybrid swarms of phenotypically diverse individuals. Hybrids are viable and fertile, though there is some evidence for subtle abnormalities of behavior or development that might impede or filter gene flow (*25, 100*). Genetic evidence, including that presented here, suggests a long and complex history of widespread genetic introgression among populations that is not obvious at the phenotypic level.

Like most, though not all, contemporary baboon researchers (*16, 20, 99, 101–103*), we recognize the six major forms as distinct species (Main Text, Fig. 1), a taxonomy in accord with the Phylogenetic Species Concept (PSC) (*104, 105*). The PSC is widely accepted by taxonomists working on primates (*106, 107*) and other mammals (*108*) comparable to baboons in populational complexity. It conceptualizes a species as a minimal clade, a cluster of populations united by a common ancestry that is attested by a unique and consistent set of apomorphic states shared by descent (*105*). The PSC's cladistic conception of the species aligns it with other, higher ranks in the taxonomic hierarchy such as the genus and the family. The main alternative classification is based on the Biological Species Concept (BSC) (*109*). The BSC sees species principally not as clades but as genetic isolates, protected from invasion and dilution by barriers to genetic exchange. BSC species are often polytypic, including sub-populations ("subspecies") that are phenotypically distinguishable, but not genetically isolated from each other. According to BSC criteria, genetic continuity via frequent marginal interbreeding would unite all extant *Papio* baboons in a single, polytypic, species (*Papio hamadryas*), with the major forms named as well-defined subspecies (*110*). As is now generally recognized (*111*), and is evident in the present work, neither of the major species concepts can adequately represent the full complexity of population history and diversification, and the choice between them is largely aesthetic. We prefer the PSC for the reasons outlined elsewhere (*112–114*). We do, however, adopt Groves' slightly less stringent version that permits naming of minor geographic variants within the species as subspecies (*101*).

Thus, we find no simple model to be entirely satisfactory in this interesting but daunting taxonomic situation. Nevertheless, some system of nomenclature that facilitates unambiguous discussion concerning variation among the individual animals and populations under study is required. Taxonomic recognition at the level of species is a suitable framework for nomenclature and effective communication regarding these diverse populations.

#### S2: Homoplasy in the *Alu* and L1 insertion data

Maximum parsimony output statistics, particularly the homoplasy index (HI) is consistently about 0.85 for whole-genome *Alu* and L1-based baboon phylogenies, compared to near zero in human studies. Because MEIs are unidirectional insertions, the high HIs indicate extensive reticulation among the sampled populations such that their autosomal genomes have gradually become more similar. By contrast, male Y-chromosome *Alu* insertions are from a single genetic system (Y-chromosome), and the HI is considerably lower (0.311) compared to inheriting a copy from each parental lineage, often from different populations. The Y-chromosome *Alu*-based tree also supports the theory of active and ongoing male swamping by the placement of an olive baboon with a Y-chromosome of eastern yellow baboons. All MEI-based maximum parsimony trees place western yellow baboons from Mahale, Katavi, and Issa Valley with Kinda baboons rather than eastern yellow baboons. This provides additional support suggesting that the western yellow baboons derived from Kinda baboon ancestry. Using MELT to ascertain MEIs from individual BAM files places equal evolutionary distance among these baboon individuals compared to the ancestral reference genome, rhesus macaque, minimizing directional bias. This further suggests that the MEIs in these datasets originated from multiple source individuals and populations over the same time frame.

#### S3: Further explanation of functional analyses

Genome regions that were the most distinct between Kinda and yellow baboons were enriched for genes linked to skeletal development and morphogenesis (e.g., embryonic skeletal system development, adjusted p = $1.77\times10^{-4}$) including specifically limb development (e.g., embryonic forelimb morphogenesis, adj. p = 0.0202). This selection on skeletal development may be reflected in size and morphometric differences between the two species: Kinda baboons are the smallest baboon species with a long-limbed build (*16*). Genes linked to male sex differentiation were also more likely than expected by chance to be in outlier high-differentiation windows (adj. p = 0.0484), which may reflect the low degree of sexual dimorphism observed in Kinda baboons, hypothesized to have resulted from reduced direct male-male competition (*50*).

Somewhat surprisingly given their stark phenotypic, behavioral, and ecological differences, there were no significantly enriched (FDR=10%) GO terms among the genomic regions most differentiated between olive and hamadryas baboons. The top GO annotations included disparate terms, including negative regulation of the force of heart contraction by chemical signal (adj. p = 0.144) and regulation of protein catabolic process (adj. p = 0.144). Similarly, no significant differences existed for the olive versus Guinea comparison.

When eastern and western yellow baboons were compared, the over-represented GO terms were similar to those of the comparison between all yellow and Kinda baboons. The GO biological process with the strongest evidence of enrichment for high-differentiation windows, for instance, was embryonic skeletal system development (adj. p = $1.315\times10^{-2}$). Such differences may reflect genomic similarity between the closely related Kinda and western yellow baboons and a shared history of selection on body proportions and size. Several sex- and reproduction-linked functional annotations were also over-represented among regions most distinct between the two subgroups of yellow baboons. These included uterus development (adj. p = 0.0132) and sex differentiation (adj. p = 0.0477).

#### S4: Anomalous individuals PD0266 and PD0662

A male olive baboon from Tarangire (PD0266) showed a similar genomic composition in the autosomes (figs. S1 and S5), Y-chromosome (fig. S7), and mtDNA (fig. S8) as other baboons from Tarangire, but was clearly separated from olive baboons in the PCA (fig. S2) and ML tree (fig. S6) based on X-chromosomal data. The patterns of most recent coancestry with northern vs. southern species along chromosome 8 resemble that of other Tarangire individuals, but the patterns along chromosome X reveal very long stretches of admixed southern haplotypes, indicating very recent admixture (fig. S11).

The female yellow baboon from Issa Valley (PD0662) clustered with the geographically close western yellow baboons in X-chromosomal (figs. S2 and S6) and mtDNA datasets (fig. S8), but was clearly separated from western yellow baboons in the PCA (fig. S1) and represented a sister lineage to the northern clade in the ML tree (fig. S5) when autosomal data was used. This individual also exhibits more olive ancestry than the Mahale baboon females, showing that its underlying ancestry is different from the other western yellow baboons (figs. S11 and S30).

These two cases show that individuals within a population can be highly heterogeneous. These anomalous animals may indicate the presence of additional hybrid zones.

#### S5: Exploring admixturegraph topologies

We selected positions where all individuals pass individual-wise filters (Methods), resulting in a final set of 48,730,011 high-quality positions. In coherence with observations from other methods, we decided to stratify the population assigned to yellow baboons into two different groups, geographically observed in eastern Tanzania (Ruaha, Mikumi, Selous, Udzungwa) and western Tanzania (Mahale, Katavi). We excluded the individual from Issa Valley.

Based on f2-statistics, we used a specified number of admixture edges from 1 to 10 to determine the best fitting graph, defining Gelada as the outgroup (fig. S27). The likelihood and out-of-sample scores are shown in table S4: We observe decreasing values for both measures with increasing complexity, most likely due to the increasing complexity of the models rather than their accuracy. When using bootstrap-resampling to fit the graphs, we find that the more complex graph provides a significantly better fit in most cases ($p < 0.05$). The first time a more complex graph is not significantly better than the simpler one is when introducing 5 instead of 4 admixture edges. The graph with 4 edges adds several layers of ancient substructure within the southern branch of baboons.

We share the cautions and caveats described in detail by the authors of the program (https://github.com/uqrmaie1/admixtools), and consider the most complex models as unlikely since they don't recover the known species phylogeny. Hence, we explored whether the same graph was recovered when exploring the graph space multiple times, or whether different graphs were found to fit best (fig. S28). We find that only 16 (64%) graphs with one admixture event match the best graph inferred above (western yellow as a mixture of Kinda and olive baboons). Another 7 (28%) graphs infer the chacma baboon population to be a mixture of Kinda and yellow baboons, and 3 graphs show different topologies each. More complex graphs are not well recovered either, and we conclude that automatically inferred admixture graphs are difficult to interpret in the context of baboon genomics.

As an alternative approach, we studied possible admixture graph topologies using f4-statistics, which may reflect the history of admixture events better than f2- and f3-statistics, but are computationally more expensive. We modeled the fit of all possible f4-statistics between the tested populations (i.e., all configurations of four populations), testing different topologies: A basic model grouping the yellow baboon populations together, a model where the eastern yellow baboons were basal to the other southern populations, and the model from a previous study (*23*), based on a smaller number of genomes (fig. S29). Among these, the (*23*) model had a lower residual error when fitting the graph to the data (table S5). Since this model was also more complex, with two admixture edges, we explored the possible space of graphs with one admixture edge. We find the best support for an admixture event from olive (14%) into western yellow baboons. The fit of this model is considerably better than the (*23*) model (table S5). The second- and third-best fitting models place the source of the admixture to Guinea or the common ancestor of Guinea and olive baboons (fig. S29).

Exploring the addition of another admixture edge to the best fitting graphs suggests a substructure within the ancestral baboon populations. Such substructure in the ancestor of Kinda baboons, as in the best model with one more event, has been described previously (*23*), but might reflect ancient substructure or contributions of unknown lineages rather than recent events. The residual errors from f4-statistic-based fits and the likelihood scores for f3-statistic-based qpgraph-calculations on the same tree topologies differ between the different graphs in the same directions (table S5). The likelihood scores are significantly different ($p < 0.01$) between the best fit with one admixture edge and the (*23*) model, and between the best fits with one and two admixture edges. Finally, when testing the f3-statistic-based models with up to 10 admixture edges, we find a worse fit of these when using the f4-statistic, in comparison to the first best model with one more edge (residual error = 166,118, compared to 3,792,813-4,129,094). This implies that the genetic drift is well characterized by these models, while gene flow is less accurately represented.

We conclude that at the current stage, admixture graphs are not sufficient to fully model the complex population history of baboons. However, gene flow from olive baboons into western yellow baboons is well-supported, and more fine-scaled sampling of baboon populations from different geographic areas may lend further support to other scenarios. Furthermore, deep structure within southern baboons seems likely, and we hypothesize that this represents a reticulated pattern of population separation, rather than clear-cut speciation, in line with the results shown above.

#### Supplementary Figures

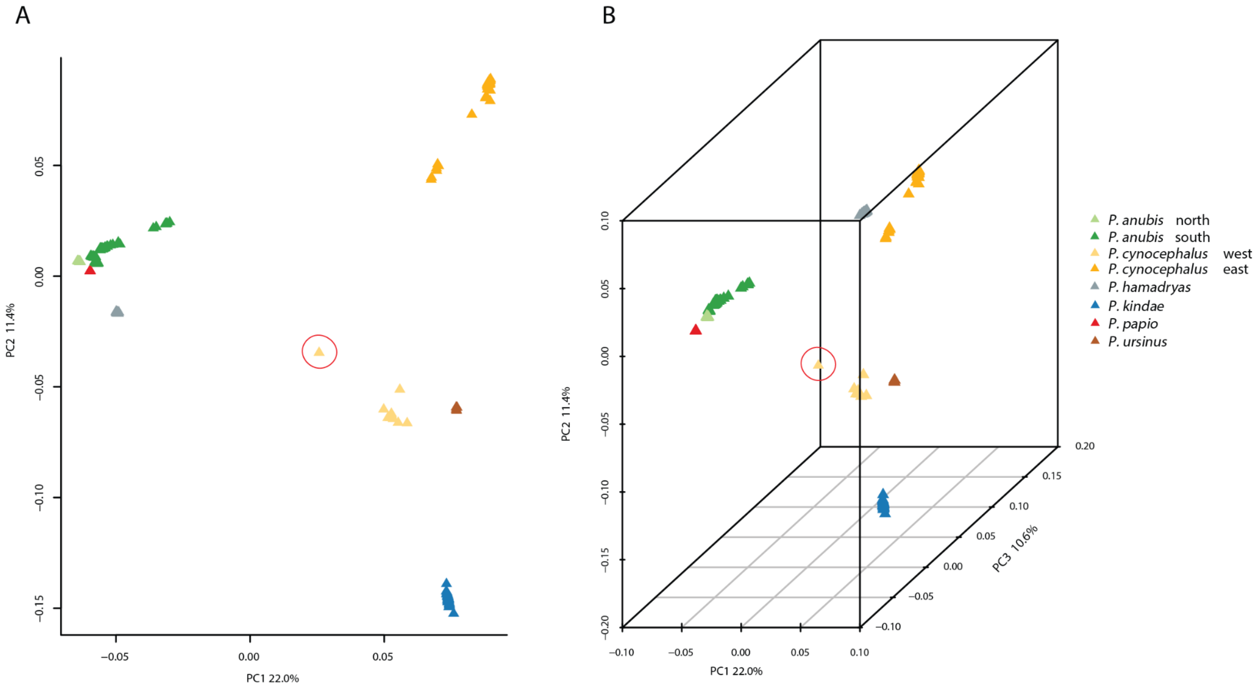


**Fig. S1.**
**Principal component analysis (PCA) among all 225 baboon individuals based on autosomal SNVs.** (**A**) Two- and (**B**) three-dimensional PCA plot. The fraction of the variance explained is 22.0% for PC1, 11.4% for PC2 and 10.6% for PC3 (for all PCs Tracy-Widom test $P < 0.01$). Yellow baboon individual PD0662 from Issa Valley is marked with a red circle.

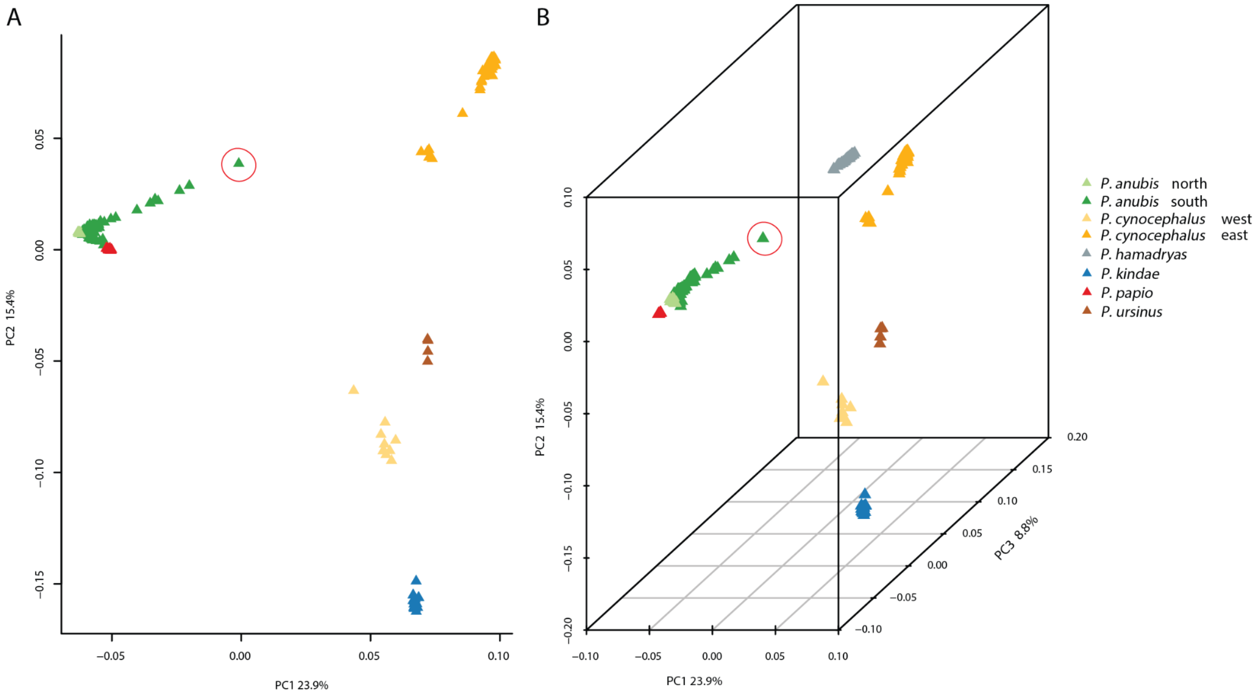


**Fig. S2.**

**Principal component analysis (PCA) among all 225 baboon individuals based on X-chromosomal SNVs.** (**A**) Two- and (**B**) three-dimensional PCA plot. The fraction of the variance explained is 23.8% for PC1, 15.4% for PC2 and 8.8% for PC3 (for all PCs Tracy-Widom test P < 0.01). Olive baboon individual PD0266 from Tarangire is marked with a red circle.

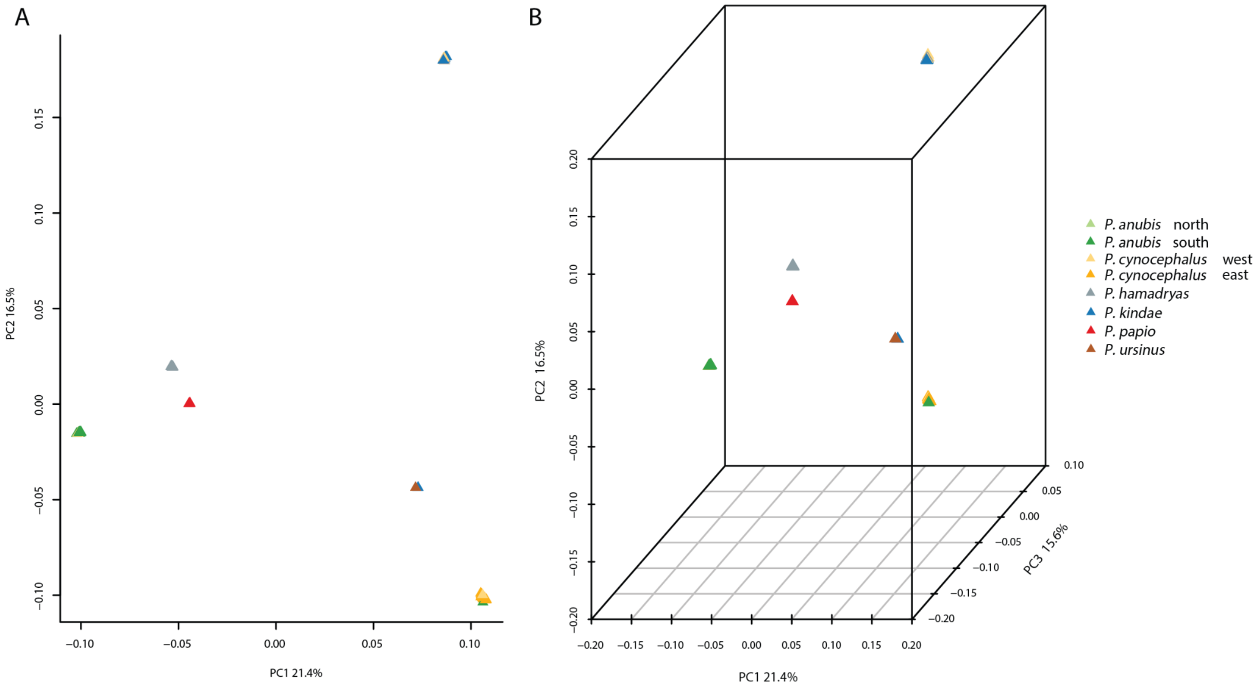


**Fig. S3.**

**Principal component analysis (PCA) among all 122 baboon males based on Y-chromosomal SNVs.** (**A**) Two- and (**B**) three-dimensional PCA plot. The fraction of the variance explained is 21.4% for PC1, 16.5% for PC2 and 15.6% for PC3 (for all PCs Tracy-Widom test P < 0.01).

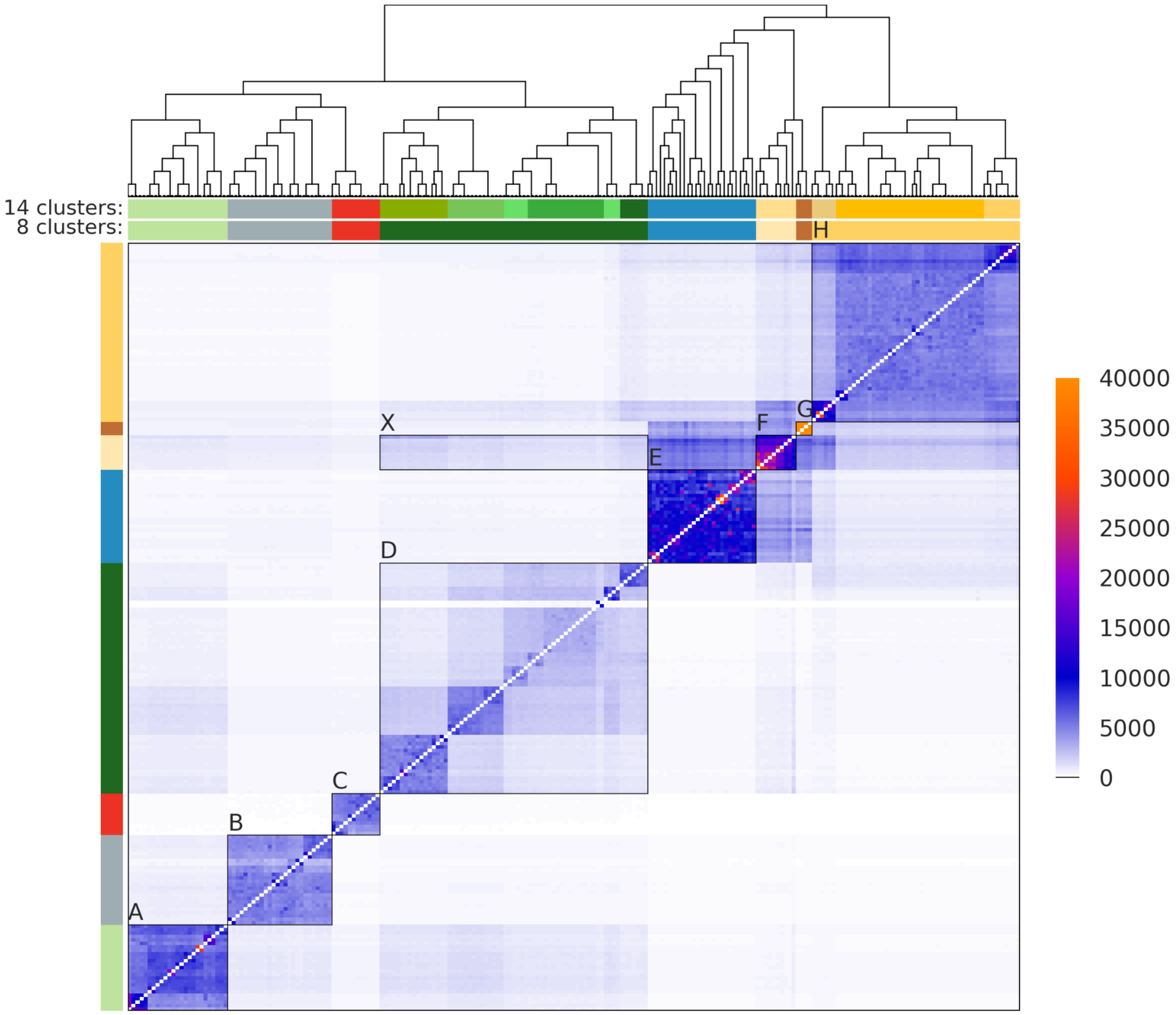


**Fig. S4.**
**Alternate fineSTRUCTURE dendrogram**. Each row represents the number of chunks copied from each other individual. In this version of the figure, the heatmap interval is set to a max of 40,000 to better represent values in the lower range, so finer details are easier to see compared to the main version.

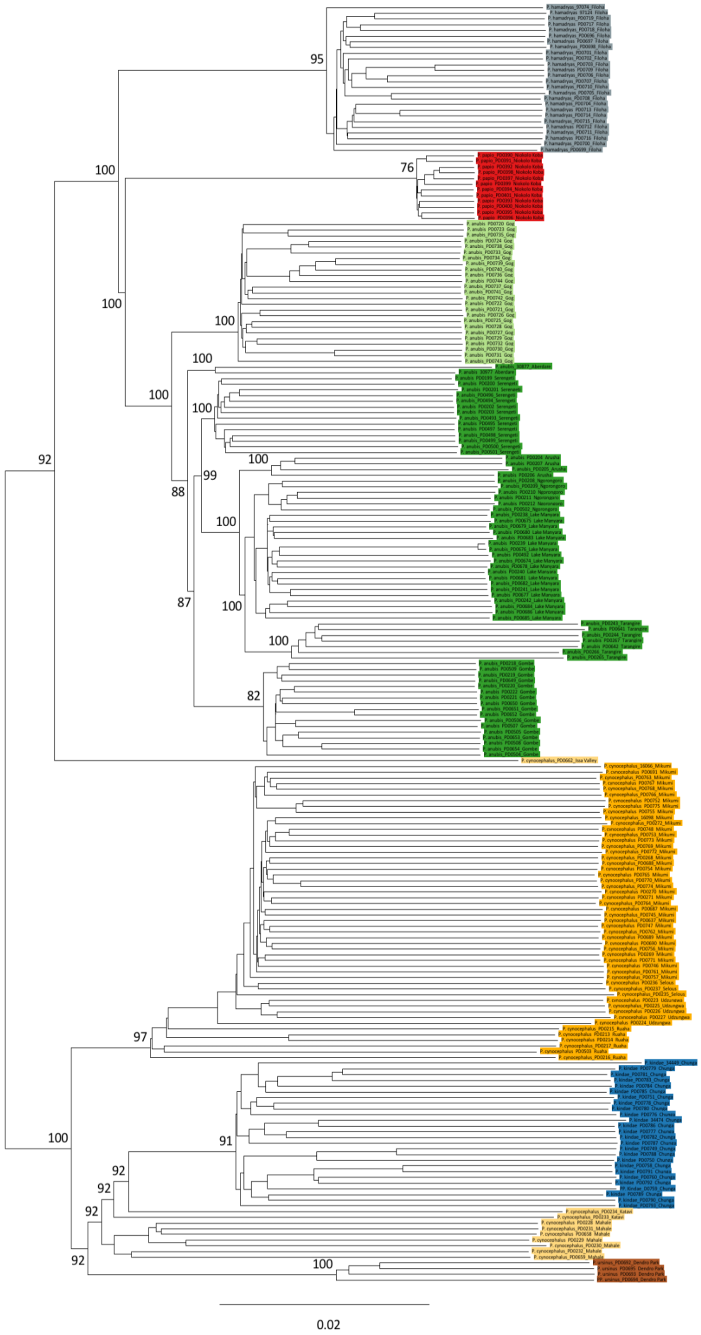


**Fig S5.**

**Maximum-likelihood tree showing phylogenetic relationships among the 225 baboon individuals based on autosomal SNVs**. Numbers at nodes refer to bootstrap values. The bar below indicates substitutions per site. Baboon individuals are colored according to phenotypical appearance. Note that yellow baboon individual PD0662 from Issa Valley forms a sister lineage to the northern clade, which is likely the result of recent introgression from olive baboons. Outgroup gelada is not shown.

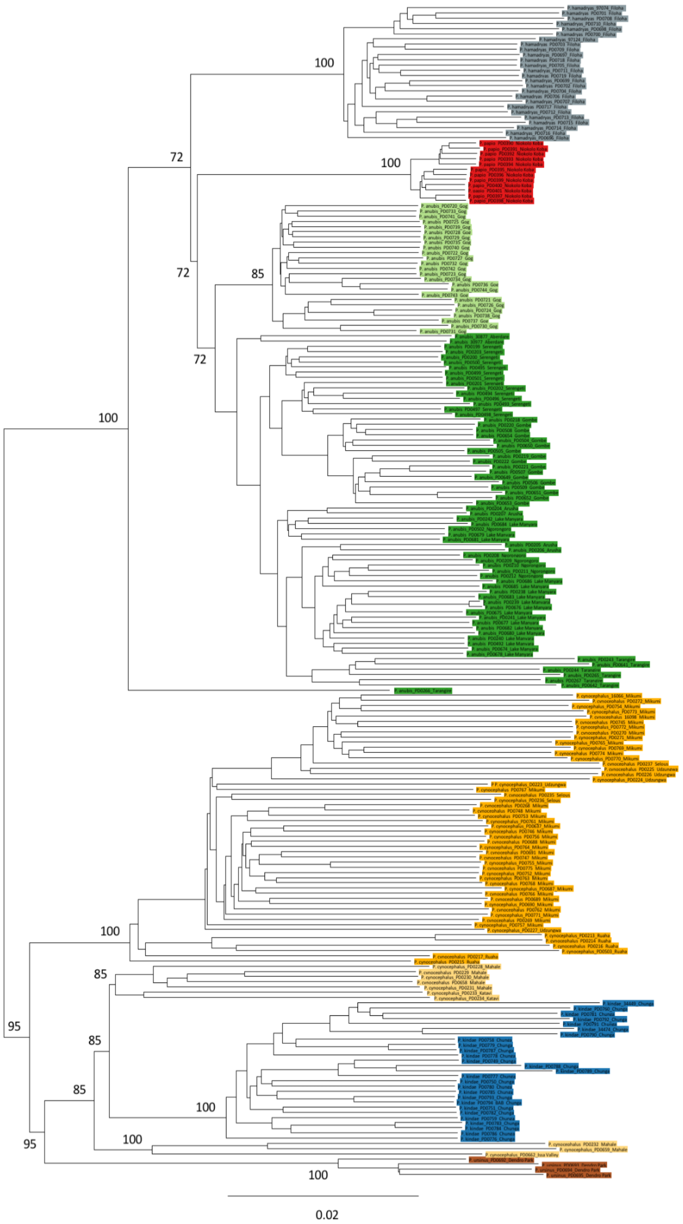


**Fig S6.**

**Maximum-likelihood tree showing phylogenetic relationships among all 225 baboon individuals based on X-chromosomal SNVs.** Numbers at nodes refer to bootstrap values. The bar below indicates substitutions per site. Baboon individuals are colored according to phenotypical appearance. Note that olive baboon individual PD0266 from Tarangire forms a distantly related lineage to the northern clade, which is likely the result of recent introgression from yellow baboons. Western yellow baboons from Mahale, Katavi, and Issa Valley are paraphyletic. Outgroup gelada is not shown.

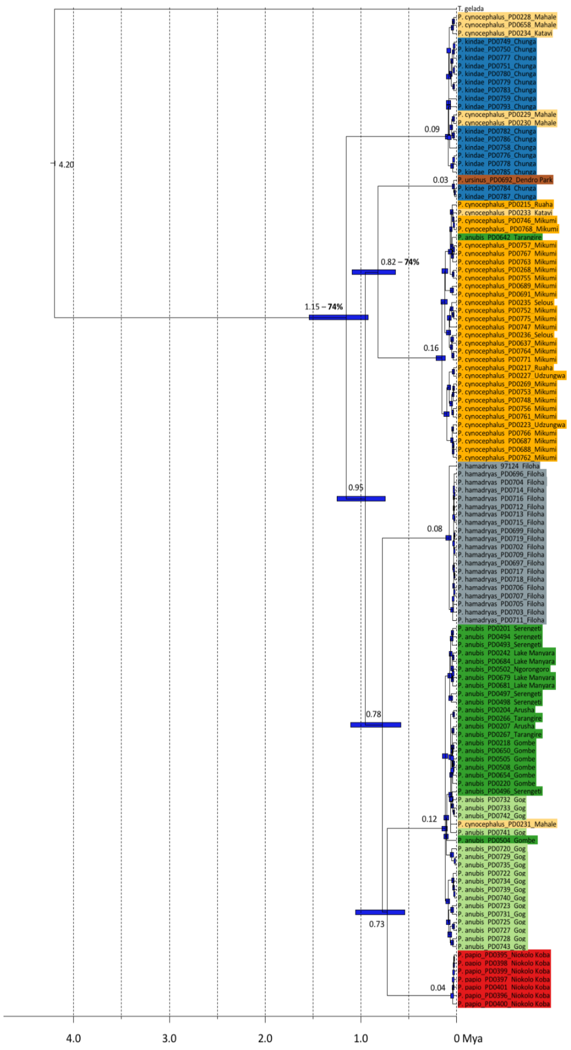


**Fig S7.**

**Maximum-likelihood time tree showing phylogenetic relationships among the 122 baboon males based on Y-chromosomal haplotypes.** Numbers at nodes refer to estimated divergence times in million years ago and blue bars indicate confidence intervals. Bootstrap values <100% are given at respective branches. Baboon individuals are colored according to phenotypical appearance. Overall, clustering of Y-chromosomal haplotypes is in agreement with phenotypical classification, with the following exceptions: western yellow baboon PD0231 clusters with olive baboons, western yellow baboon PD0233 clusters with eastern yellow baboons, southern olive baboon PD0642 clusters with eastern yellow baboons, Kinda baboons PD0784 and PD0787 cluster with chacma baboon, and western yellow baboons PD0228, PD0229, PD0230, PD0234 and PD0658 cluster with Kinda baboons.

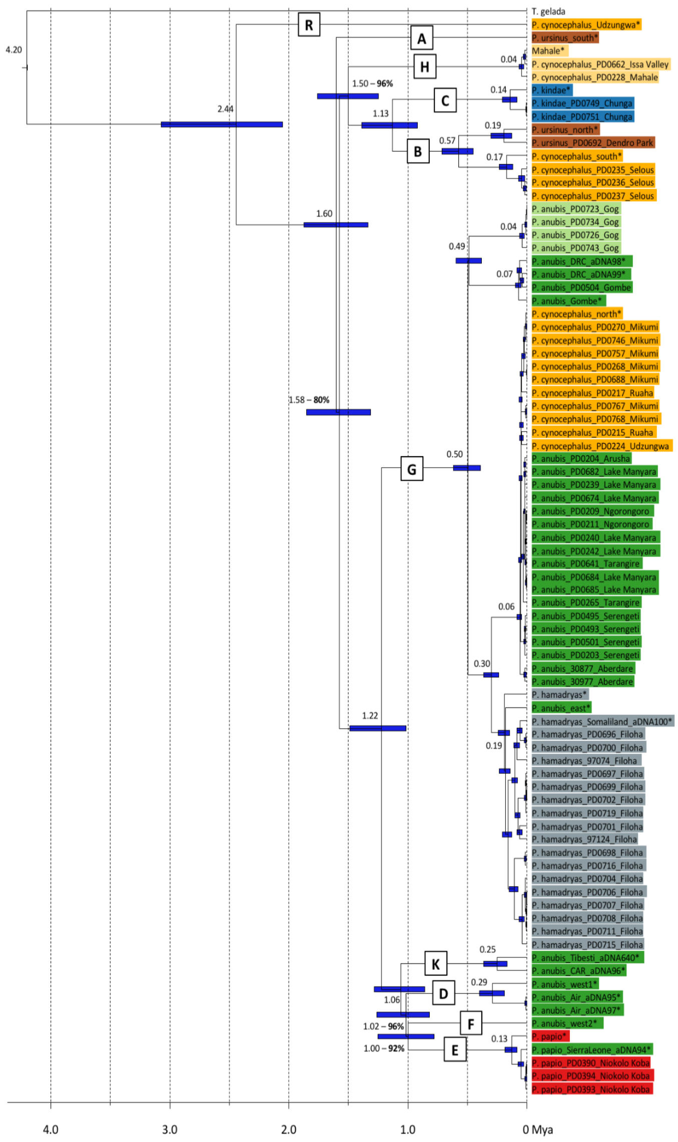


**Fig S8.**

**Maximum-likelihood time tree showing phylogenetic relationships among baboon mtDNA haplotypes.** Numbers at nodes refer to estimated divergence times in million years ago and blue bars indicate confidence intervals. Bootstrap values <100% are given at respective branches. Letters A-R on branches indicate clades as previously defined (*20, 21*). Mitochondrial genomes marked with an asterisk have been previously published (*20, 21*). Baboon individuals are colored according to phenotypical appearance. For identical haplotypes, see table S7.

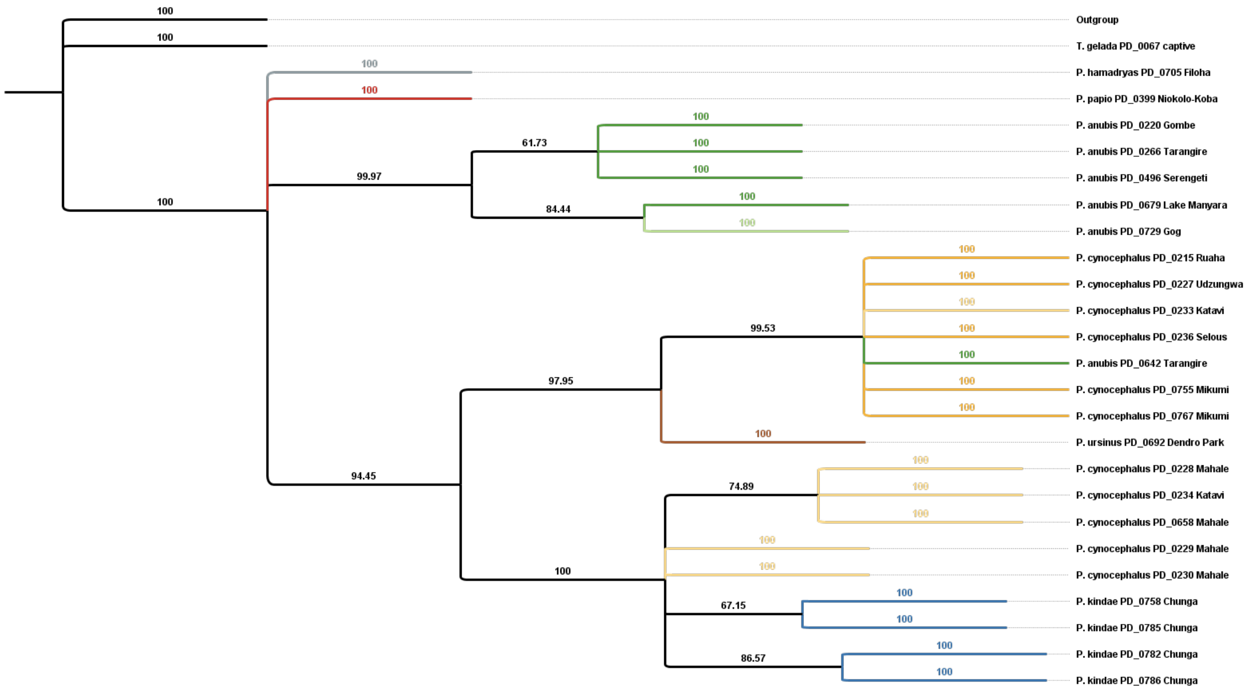


**Fig. S9.**

***Alu* insertion-based Y-chromosome tree.** A Dollo parsimony Y-chromosome tree was generated in PAUP for 24 male baboons and one gelada using 355 *Alu* insertions. All loci were set to Dollo.up for 10,000 bootstrap replicates. Numbers above branches are bootstrap values. The number of parsimony-informative characters was 169. *P. anubis,* Olive baboon individual PD0642 from Tarangire groups with eastern yellow baboons, as does western yellow baboon individual PD0233 from Katavi. Other western yellow baboons group with Kinda baboons. These results are consistent with the ML tree generated using Y-chromosomal sequence data (fig. S7). Consistency index (CI): 0.689; Retention index (RI): 0.911; Homoplasy index (HI): 0.311.

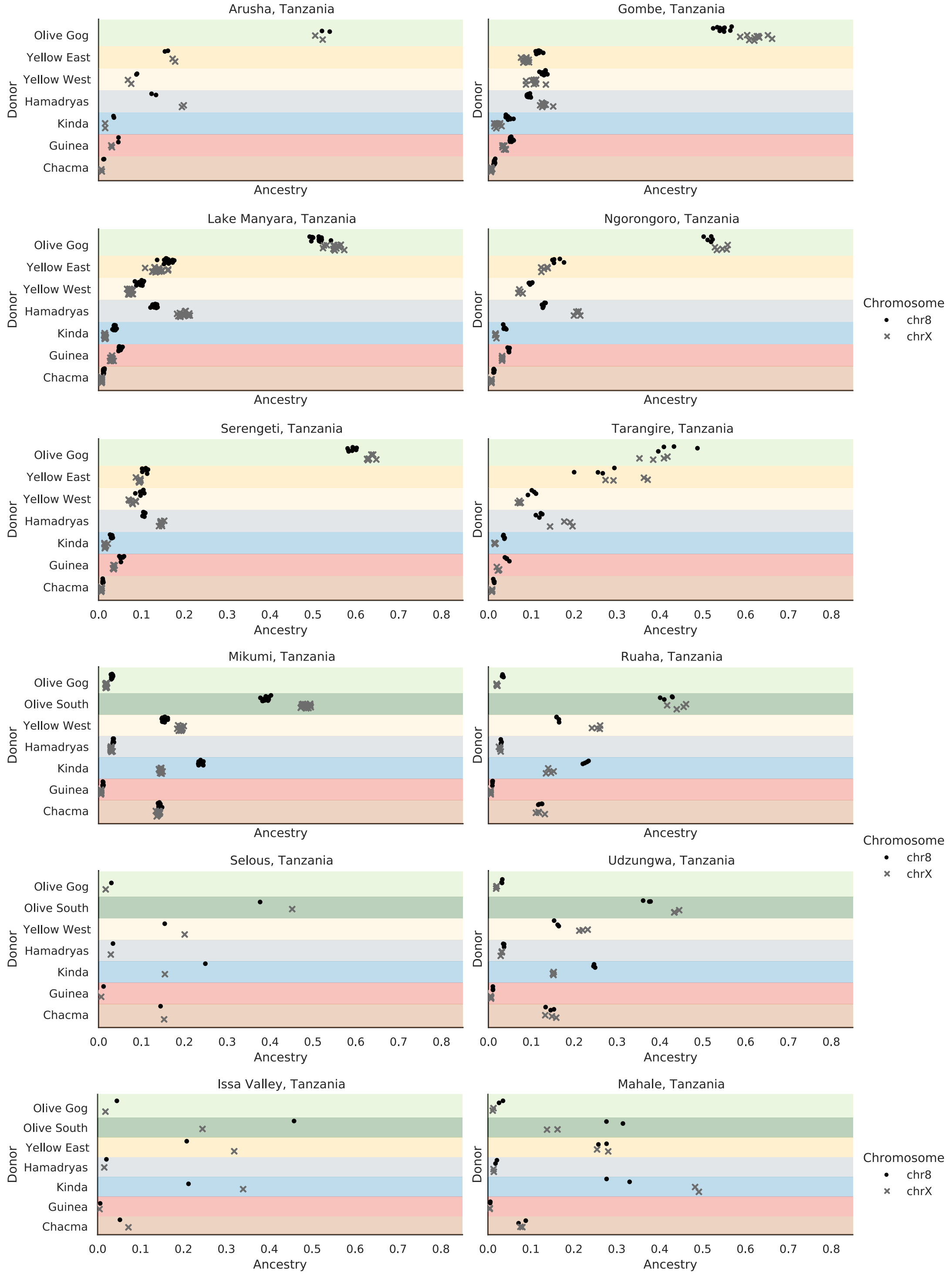


**Fig. S10.**

**Ancestry proportions of female baboons.** Depicting the Tanzanian individuals split into their individual sampling locations. Tarangire departs from the other olive baboons in their ancestry.

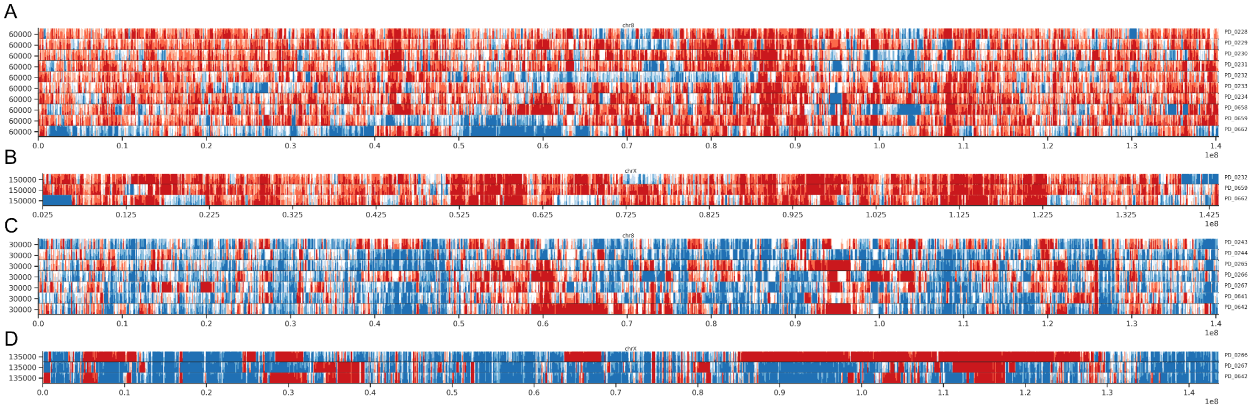

**Fig. S11.**

**Northern and southern species ancestry along chromosomes 8 and X of individuals PD0266 and PD0662.** (**A**) Most recent coancestry with northern and southern species along chromosome 8 of western yellow baboons (including individual PD0662). (**B**) Same as A but for chromosome X. (**C**) Chromosome 8 for Tarangire olive baboons (including individual PD266). (**D**) Same as B but for chromosome X. Red depicts southern ancestry, which consists of chacma, Kinda and yellow baboons. Blue depicts northern ancestry, which consists of Guinea, hamadryas and olive baboons. Horizontal panels in each subfigure are horizon plots showing the intensity of ancestry proportions of each individual in 100kb windows averaged over ten Viterbi samples by ChromoPainter. The intensity is calculated as total northern ancestry minus total ancestry divided by 2. This results in a normalized measure, with positive values (blue) showing more northern ancestry and negative values (red) showing more southern ancestry. Coancestry is defined as the most recent common ancestor not including individuals from the same population, as in the Globetrotter analysis. On chromosome X, only baboons of the same gender are included due to differences in ploidy.

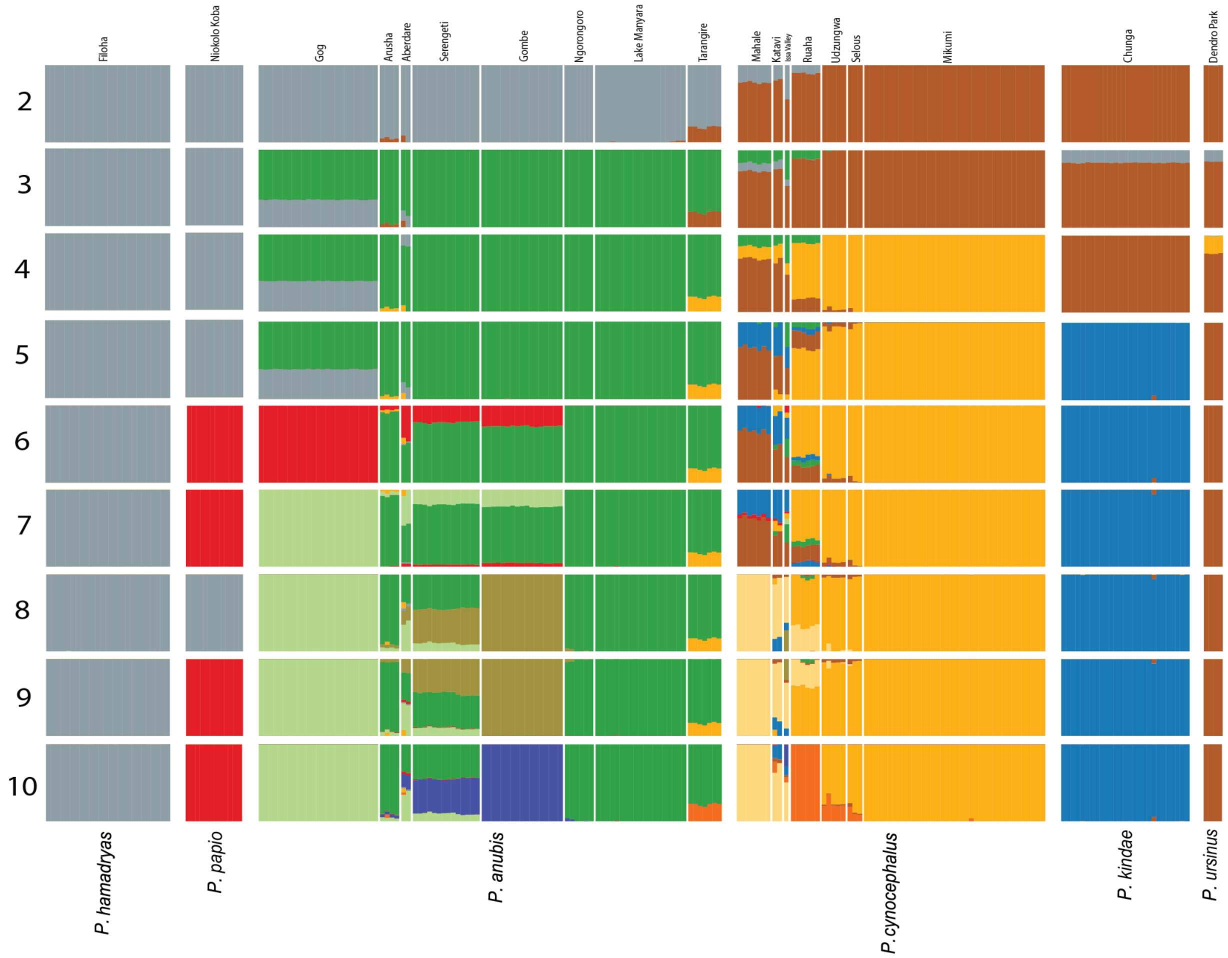


**Fig S12.**

**Population structure plots for all 225 baboon individuals with *K* = 2-10 based on autosomal SNVs**. The cross-validation error for different *Ks* were *K*=3: 0.36916, *K*=4: 0.34398, *K*=5: 0.34548, *K*=6: 0.33424, *K*=7: 0.32151, *K*=8: 0.33457, *K*=9: 0.32455, and *K*=10: 0.33261. Accordingly, a division of baboons into seven clusters is most suitable.

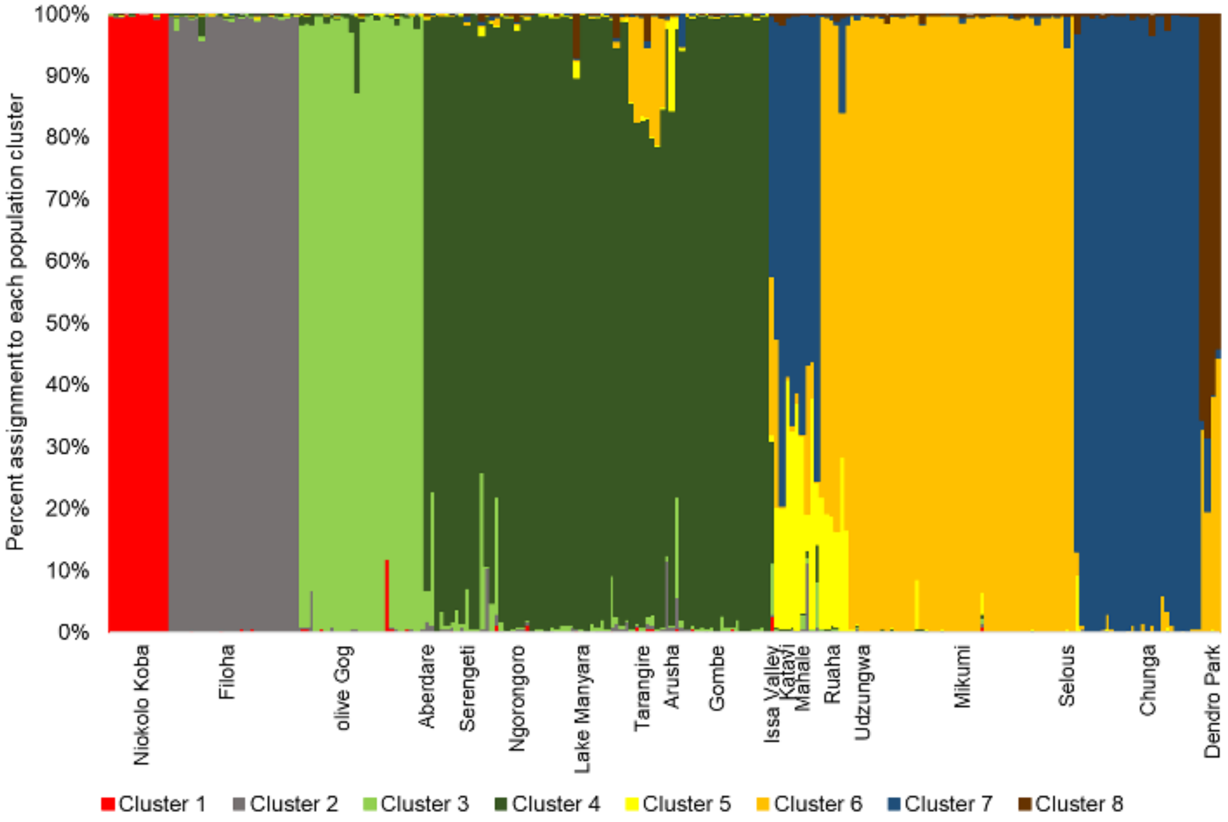


**Fig. S13.**
**Population structure using 261 L1 insertions for 222 baboons.** The optimal LnP(D) value for *K*=8 is shown. For K=2-10 the LnP(D) value for different *K*s are *K*=2: -40165, *K*=3: -37022, *K*=4: -35092, *K*=5: -33816, *K*=6: -33984, *K*=7: -38757, *K*=8: -33229, *K*=9: -63555, *K*=10: -33936.

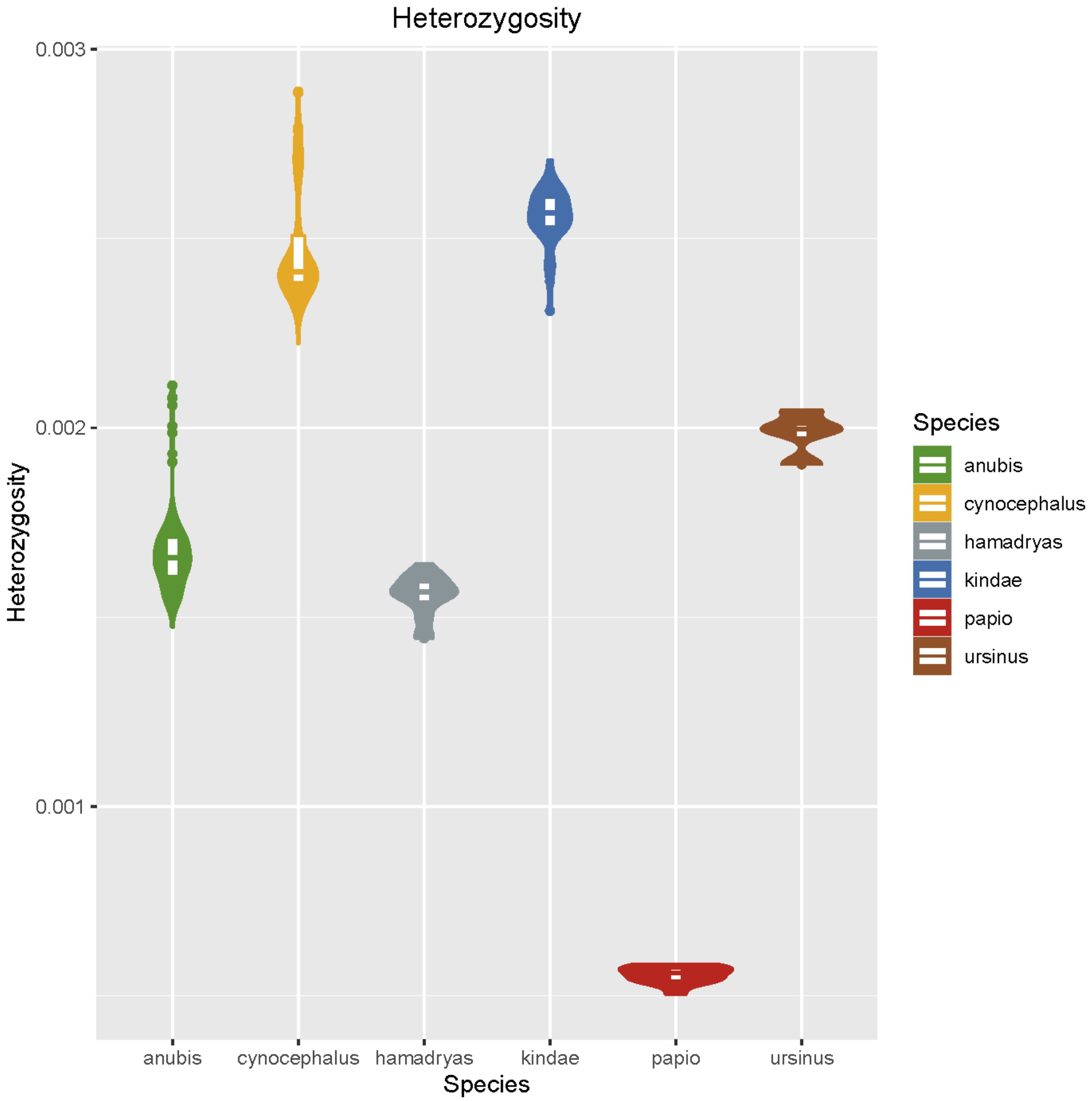


**Fig. S14.**
**Genome-wide autosomal heterozygosity of baboon species**. Heterozygosity was calculated as ((autosomal heterozygous SNV calls – missing SNV calls)/ungapped autosomal assembly length)) for each individual and then plotted separately for each species. Guinea baboons show relatively lower heterozygosity suggesting reduced diversity.

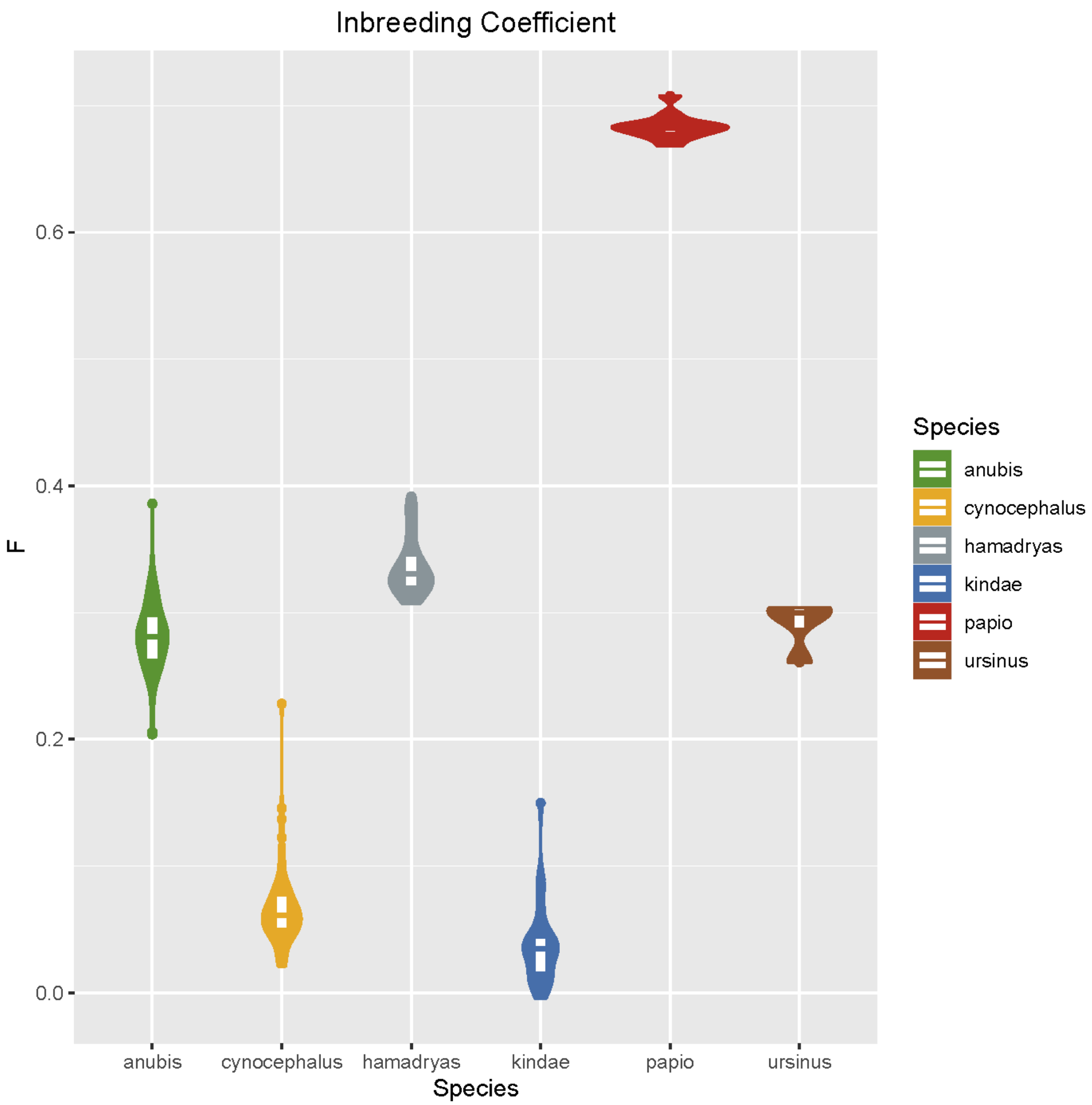


**Fig. S15.**
**Inbreeding coefficient estimates of baboon species.** Method-of-moments F inbreeding coefficient estimates were calculated with PLINK as (observed homozygous SNVs - expected homozygous SNVs) / (total called SNVs - expected homozygous SNVs) for each individual and then plotted separately for each species. Guinea baboons show relatively higher inbreeding coefficients suggesting reduced diversity.

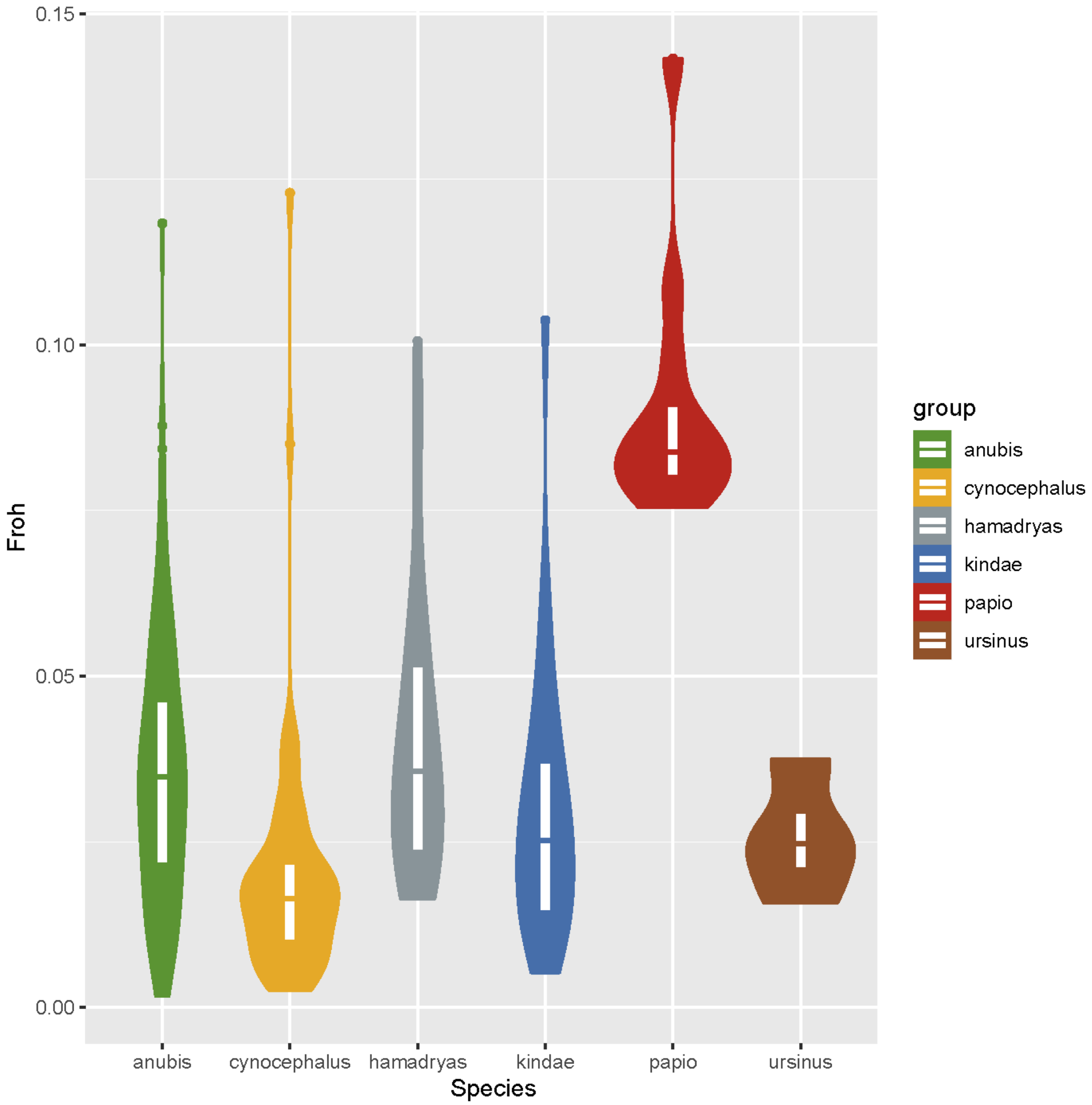


**Fig. S16.**
**$F_{ROH}$ estimates of baboon species**. ROHs were calculated by PLINK and $F_{ROH}$ was calculated as the proportion of runs of homozygosity (ROH) in the autosomal genome per individual and then plotted separately for each species. The guinea baboons show relatively higher $F_{ROH}$ suggesting reduced diversity.

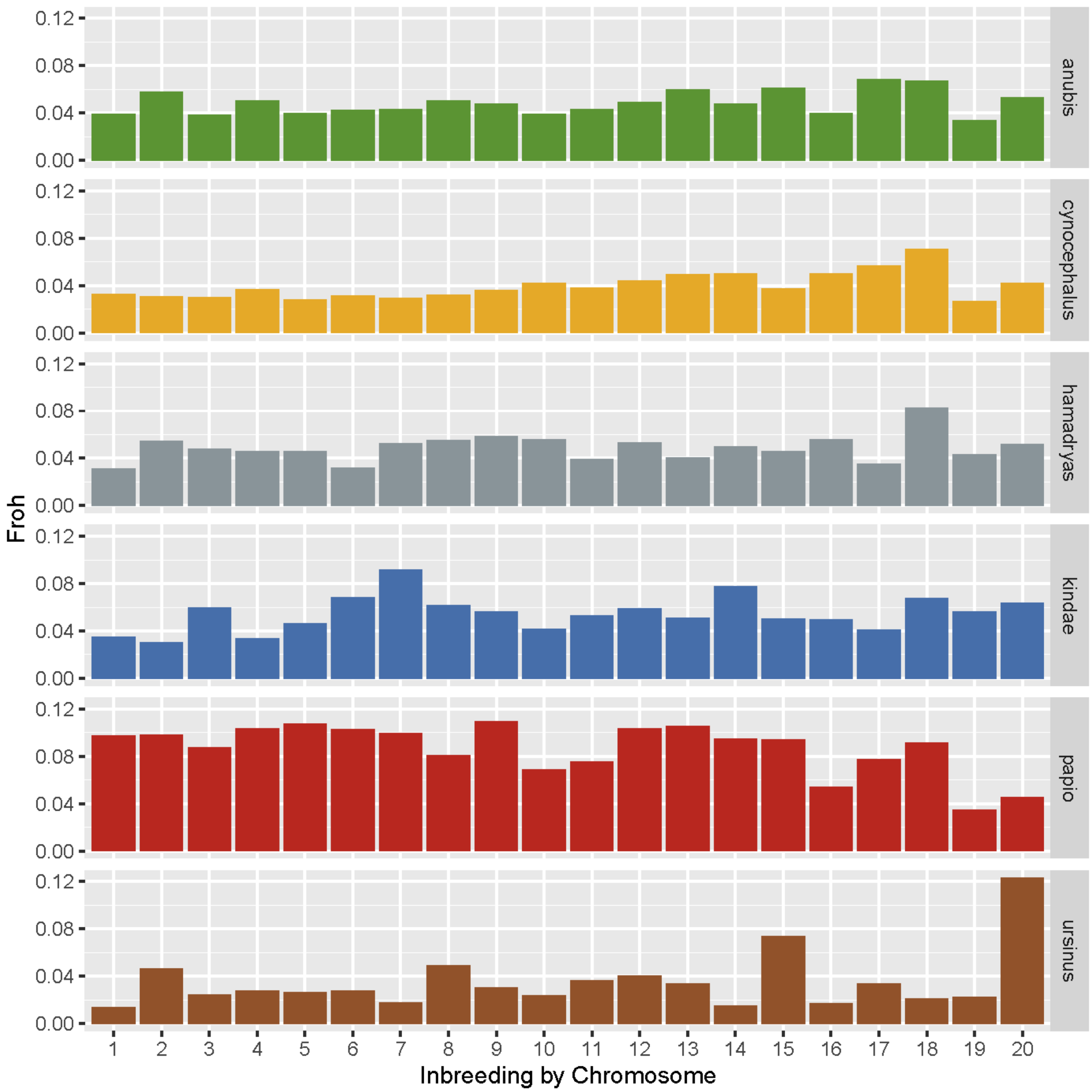


**Fig. S17.**
**$F_{ROH}$ estimates of baboon species by chromosome**. ROHs were calculated by PLINK and $F_{ROH}$ was calculated as the proportion of runs of homozygosity (ROH) in each chromosome per individual and then plotted separately for each species. The Guinea baboons show relatively higher $F_{ROH}$ in most chromosomes suggesting reduced diversity.

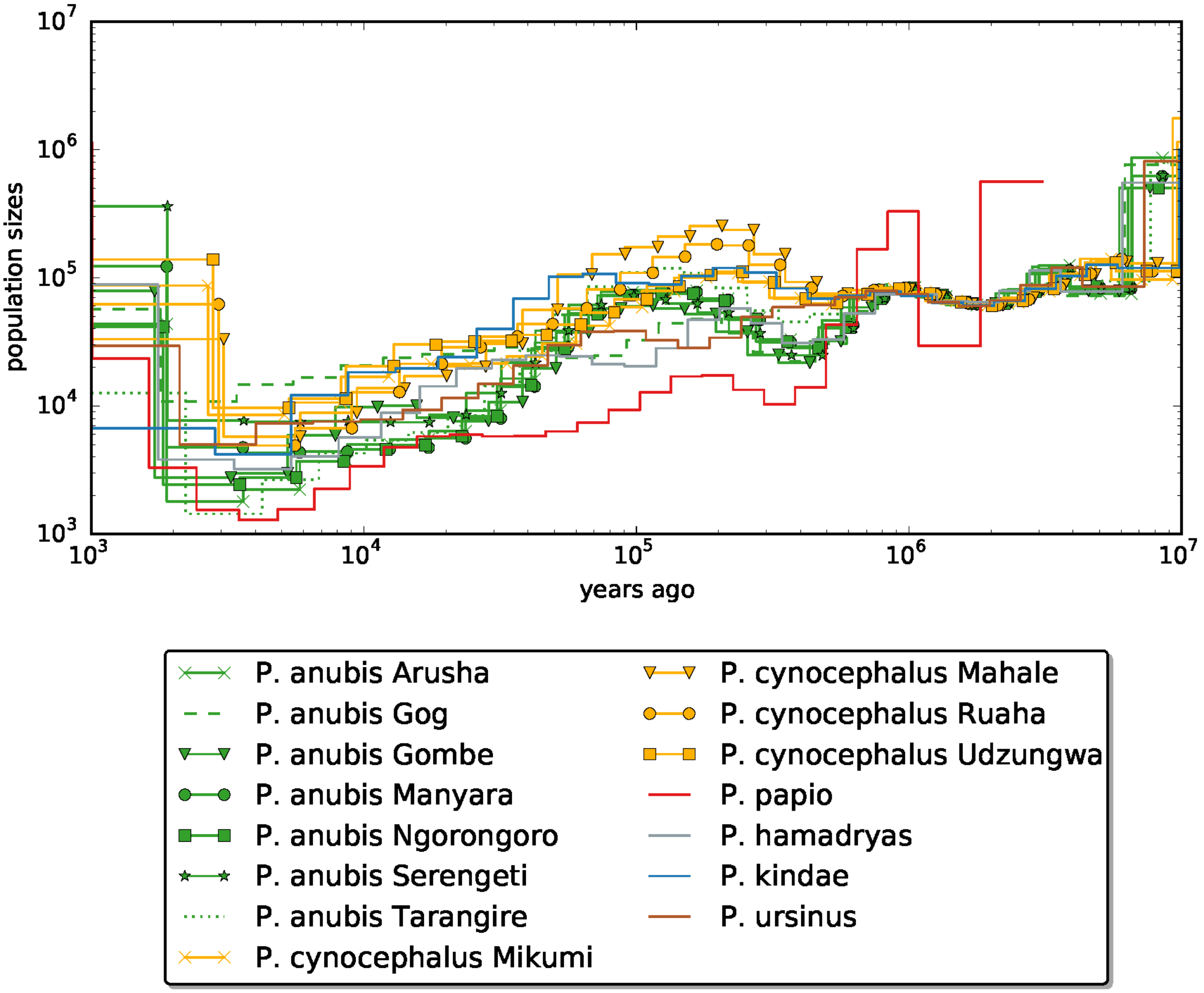


**Fig. S18.**
**MSMC2 plot of all species including populations for olive and yellow baboons**. The plots use a mutation rate of 0.9x10-8 and a generation time of 11 years (*23*). The plots were generated using four phased individuals per species or population.

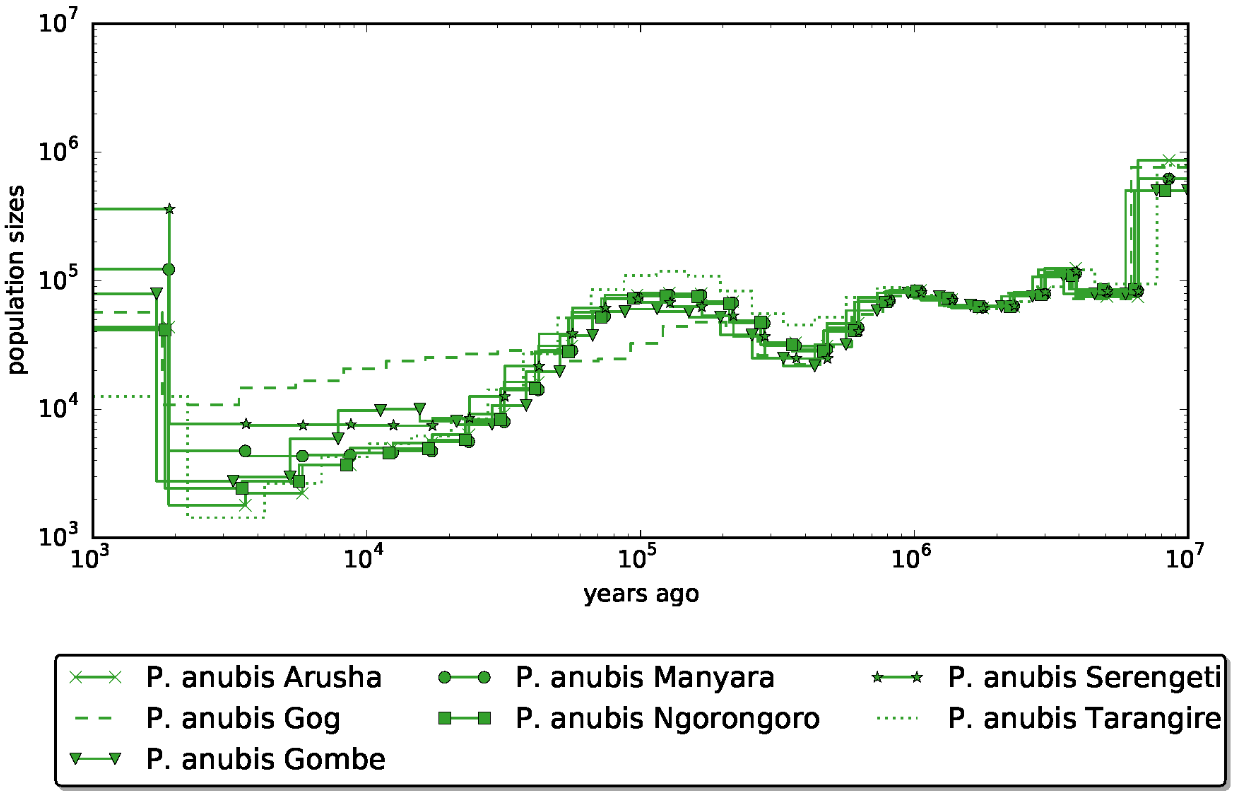


**Fig. S19.**
**MSMC2 plot of olive baboon populations**. The plots use a mutation rate of 0.9x10-8 and a generation time of 11 years (*23*). The plots were generated using four phased individuals per species or population.

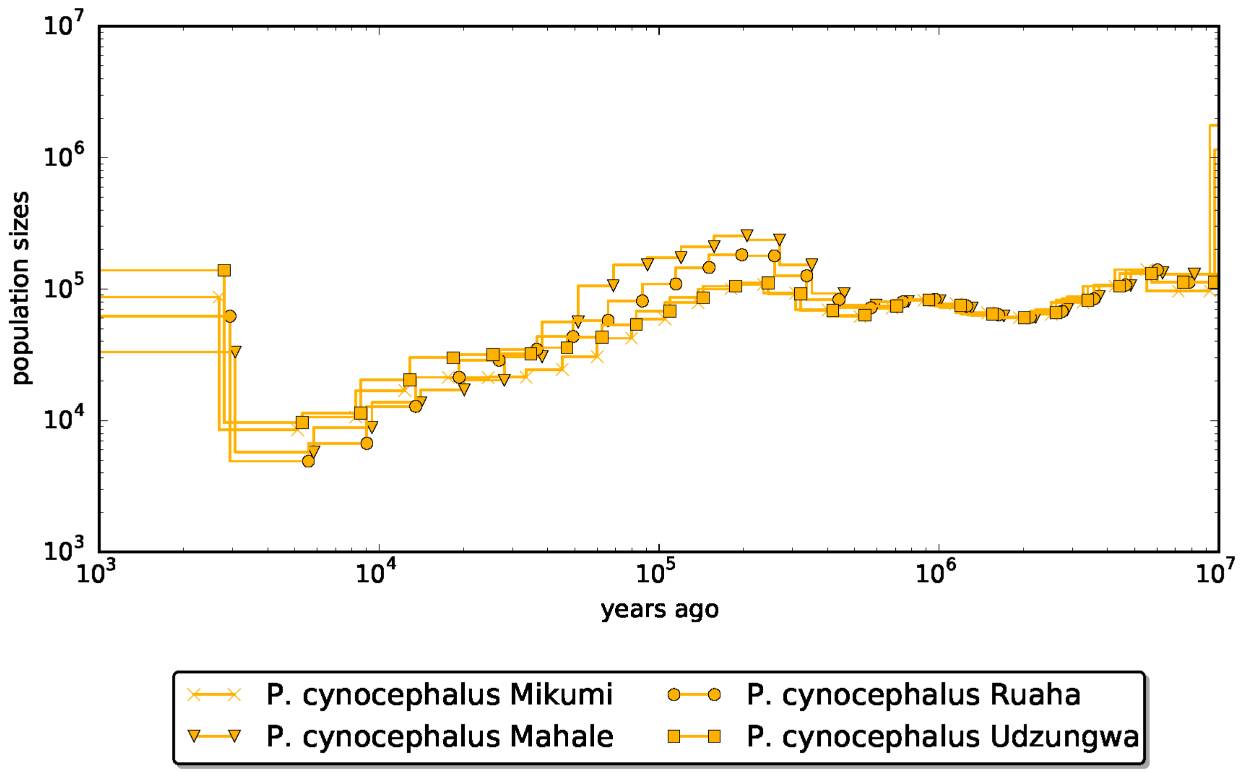


**Fig. S20.**
**MSMC2 plot of yellow baboon populations**. The plots use a mutation rate of 0.9x10-8 and a generation time of 11 years (*23*). The plots were generated using four phased individuals per species or population.

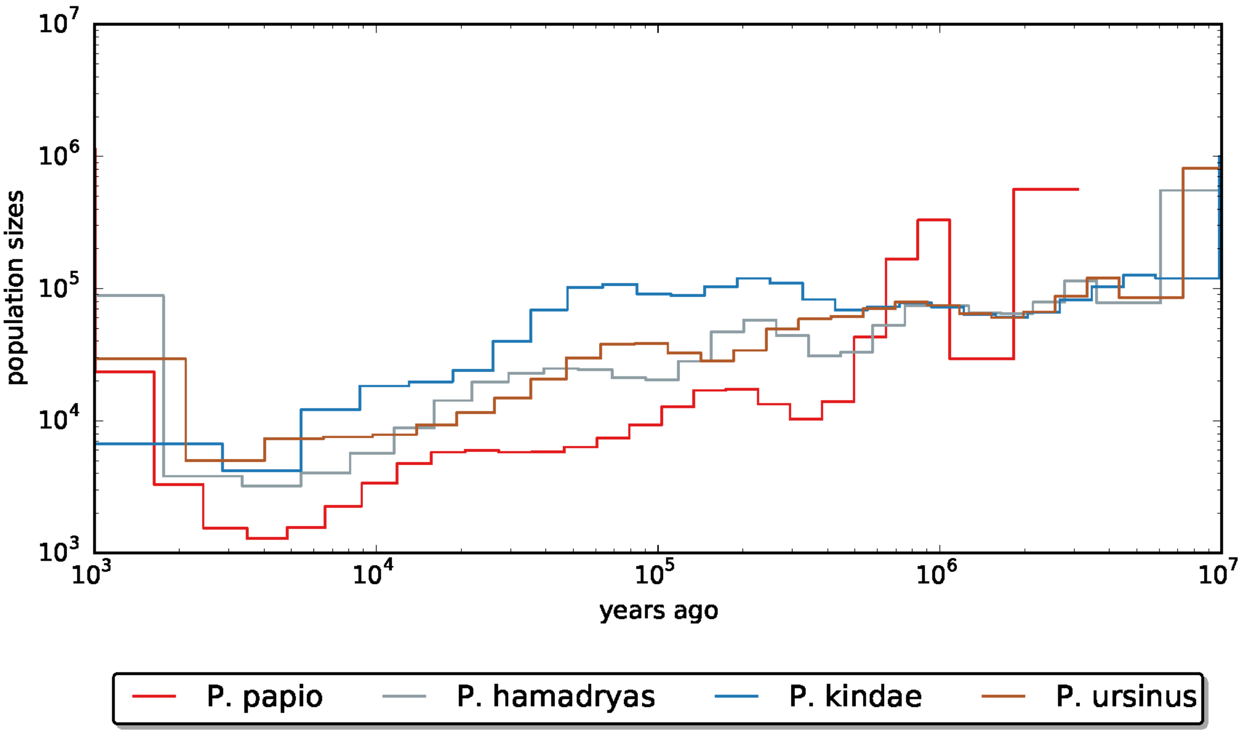


**Fig. S21.**
**MSMC2 plot of Guinea, hamadryas, Kinda and chacma baboons**. The plots use a mutation rate of 0.9x10-8 and a generation time of 11 years (*23*). The plots were generated using four phased individuals per species or population.

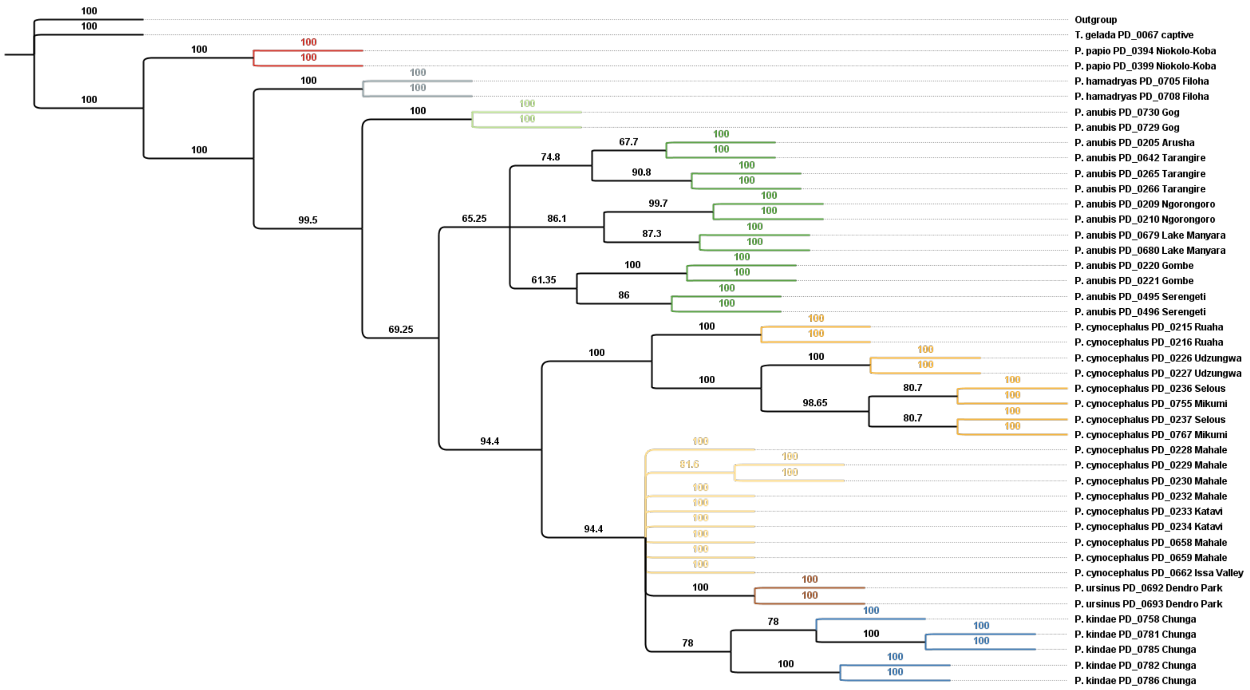


**Fig. S22.**
***Alu* insertion-based phylogeny.** A Dollo parsimony tree was generated in PAUP for 42 baboons and one gelada using 246,219 full-length *Alu* insertions from subfamily *Alu*Y. All loci were set to Dollo.up for 1,000 bootstrap replicates with maxtrees set to 100. Numbers above branches are bootstrap values. The number of parsimony-informative characters was 177,339. Branches are consistent with sampling locations. Consistency index (CI): 0.141; Retention index (RI): 0.724; Homoplasy index (HI): 0.859.

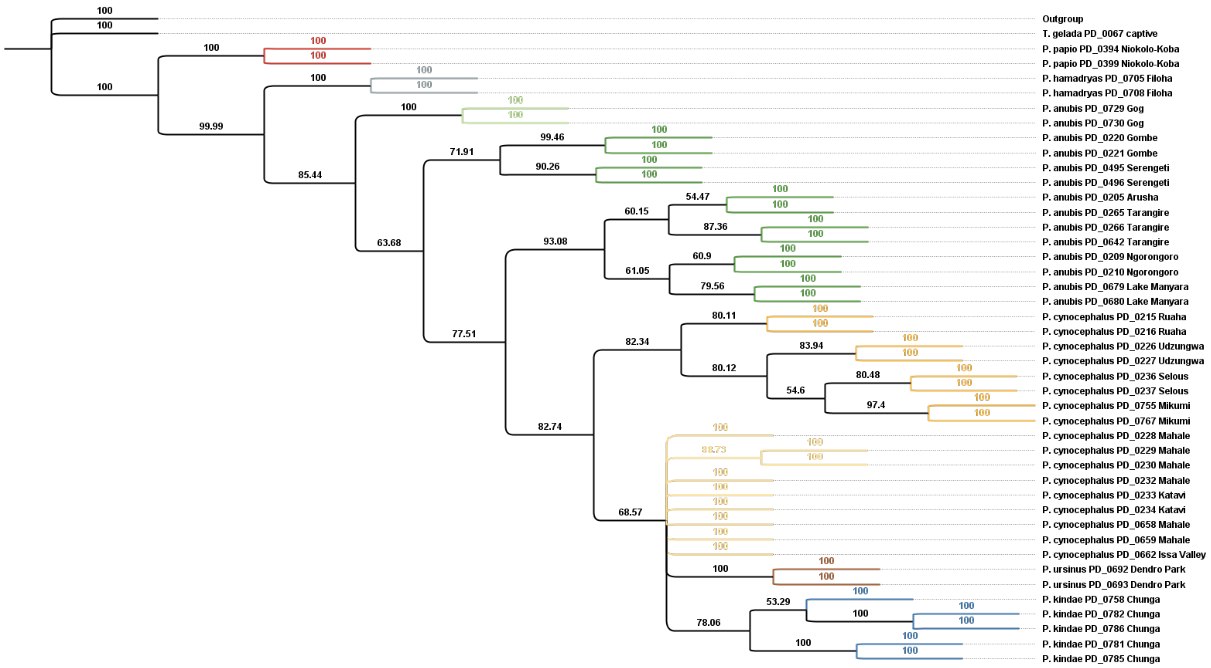


**Fig. S23.**

**L1 insertion-based phylogeny.** A Dollo parsimony tree was generated in PAUP for 42 baboons and one gelada using 35,913 L1PA6 insertions. All loci were set to Dollo.up for 10,000 bootstrap replicates with maxtrees set to 100. Numbers above branches are bootstrap values. The number of parsimony-informative characters was 24,546. Branches are consistent with sampling locations. Consistency index (CI): 0.157; Retention index (RI): 0.760; Homoplasy index (HI): 0.843.

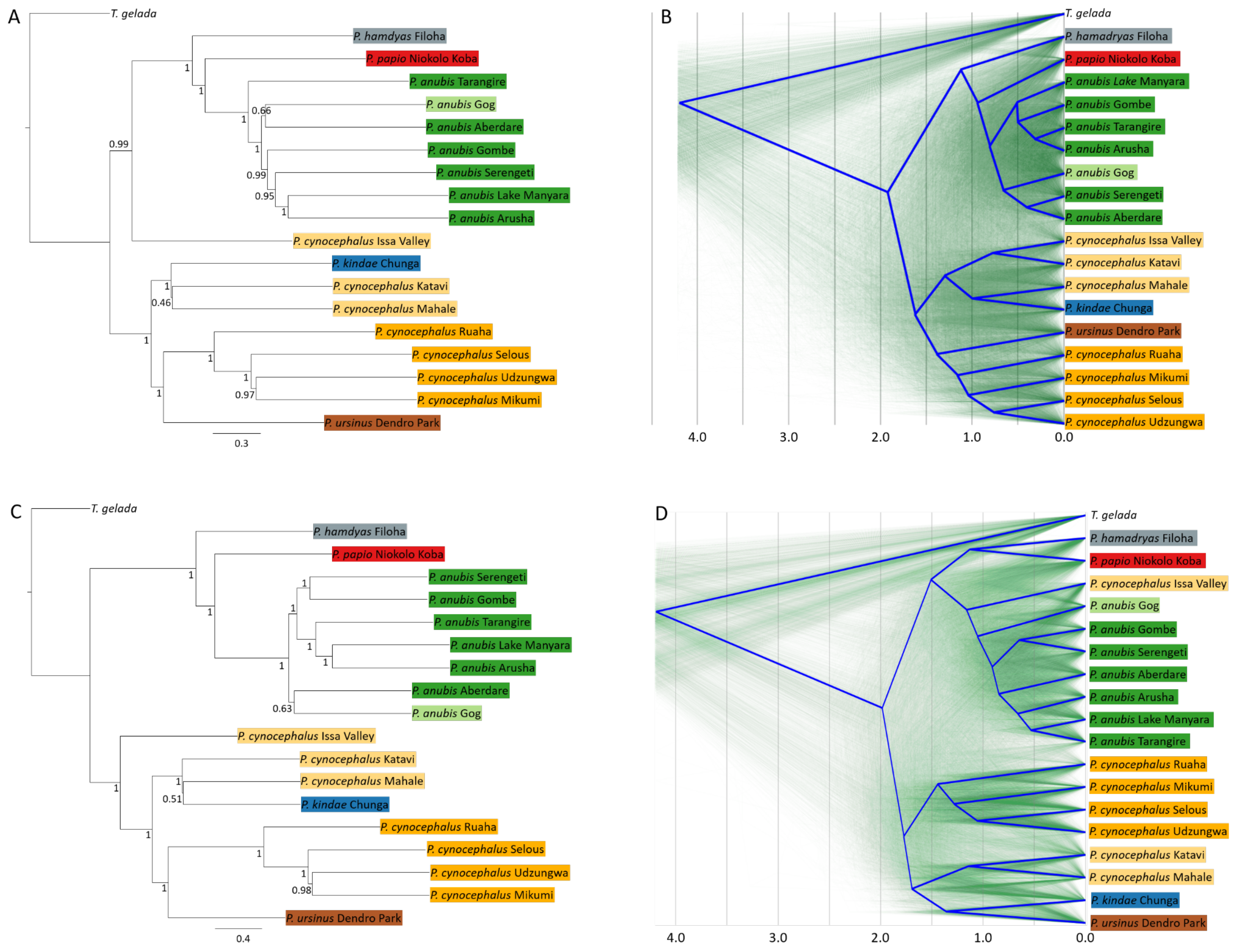


**Fig. S24.**

**Species tree and DensiTree visualization of autosomal window trees**. Trees were generated using 28,700 50kb- (**A**, **B**) and 3129 500kb-windows (**C**, **D**). In the species trees (**A**, **C**), numbers at nodes refer to local posterior probabilities and branch lengths are given in coalescent units. The normalized quartet scores, i.e. the proportion of input gene tree quartet trees satisfied by the species tree, are 0.58 and 0.70 for 50kb- and 500kb-windows, respectively. In the DensiTree visualization (**B**, **D**), green lines refer to individual gene trees and the consensus tree is highlighted in blue. The time scale below refers to million years ago.

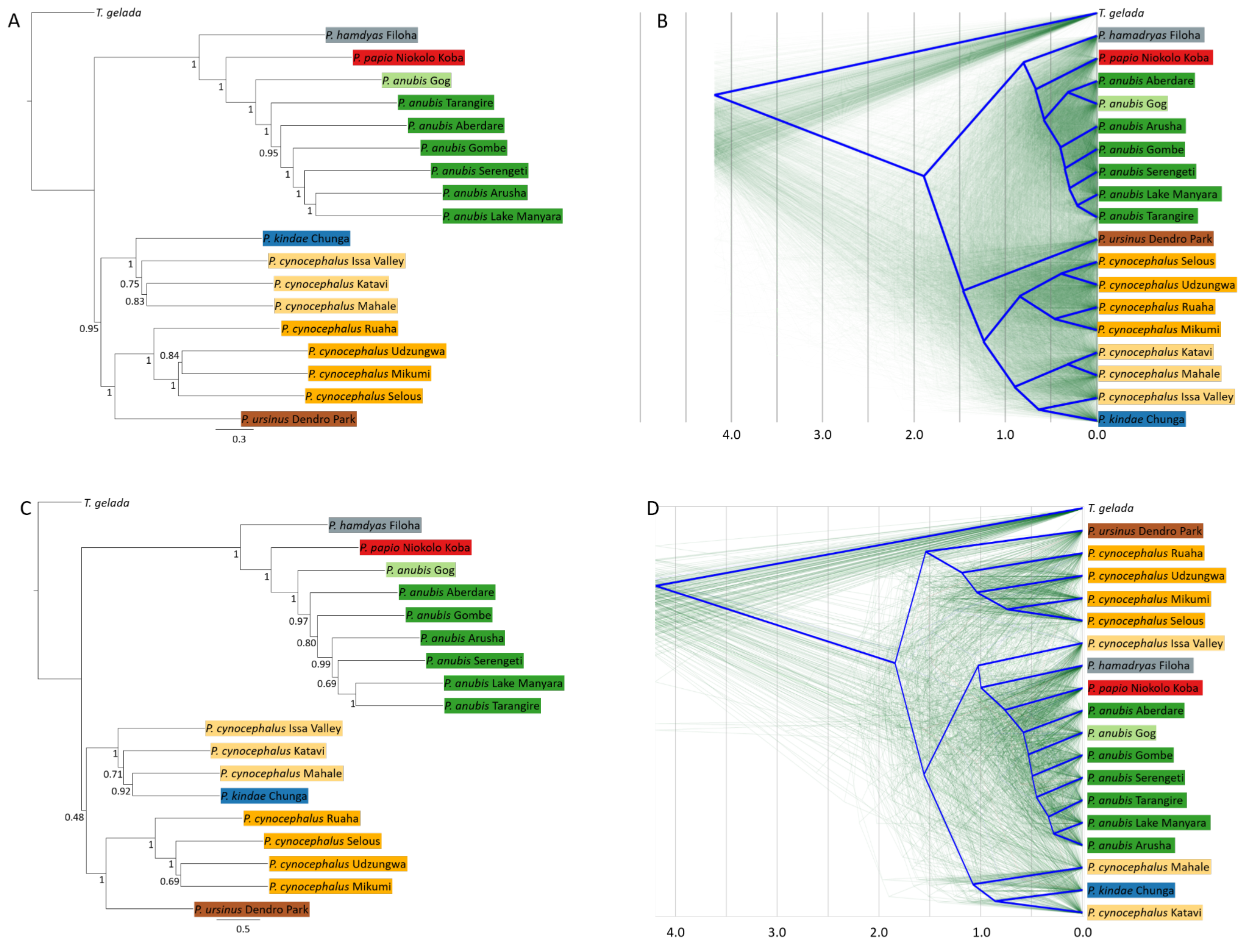


**Fig. S25.**

**Species tree and DensiTree visualization of X-chromosomal window trees**. Trees were generated using 1553 50kb- (**A**, **B**) and 165 500kb-windows (**C**, **D**). In the species trees (**A**, **C**), numbers at nodes refer to local posterior probabilities and branch lengths are given in coalescent units. The normalized quartet scores, i.e. the proportion of input gene tree quartet trees satisfied by the species tree, are 0.61 and 0.70 for 50kb- and 500kb-windows, respectively. In the DensiTree visualization (**B**, **D**), green lines refer to individual gene trees and the consensus tree is highlighted in blue. The time scale below refers to million years ago.

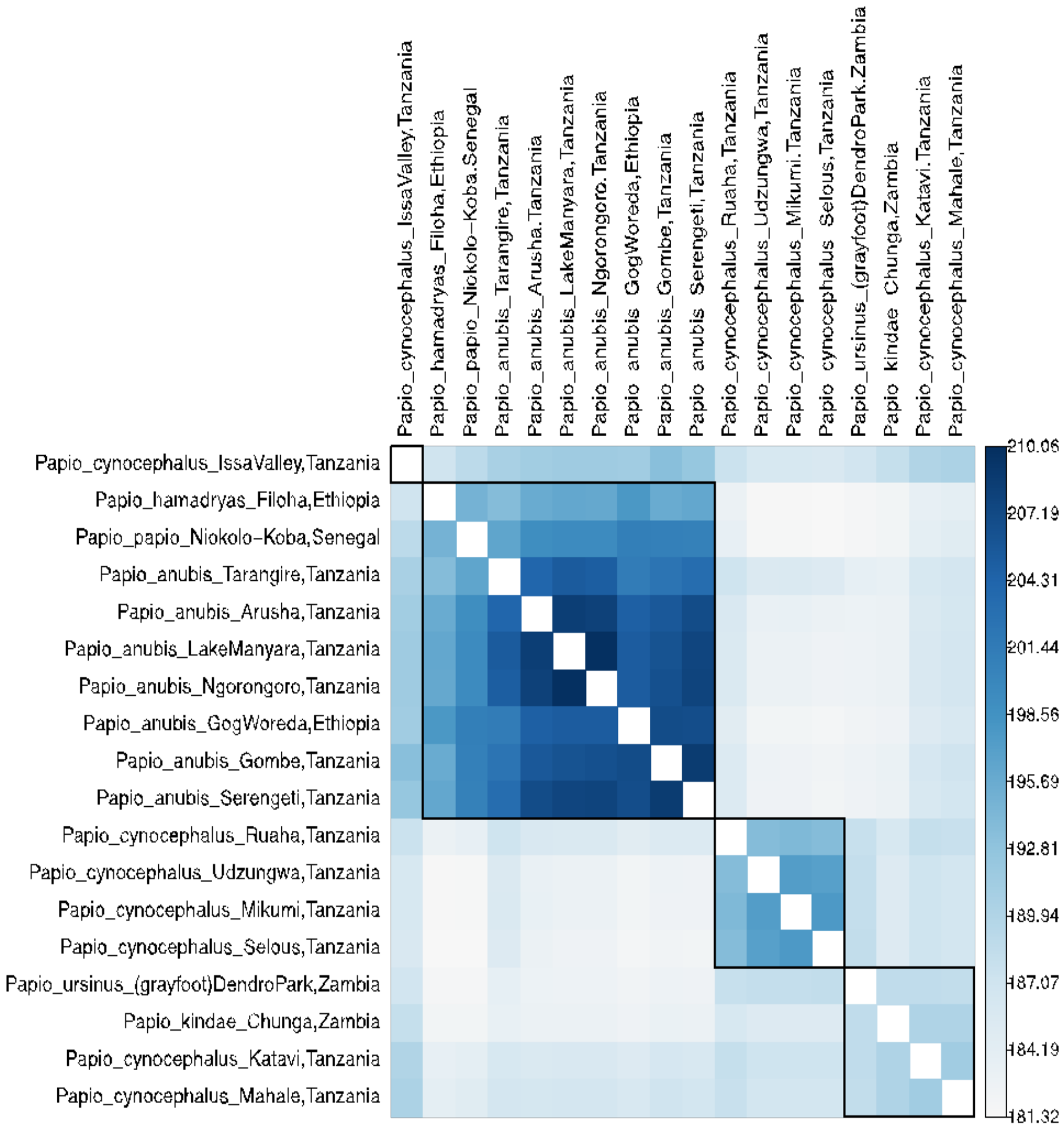


**Fig. S26.**
**F3 outgroup statistics with baboons stratified by locality and species.**

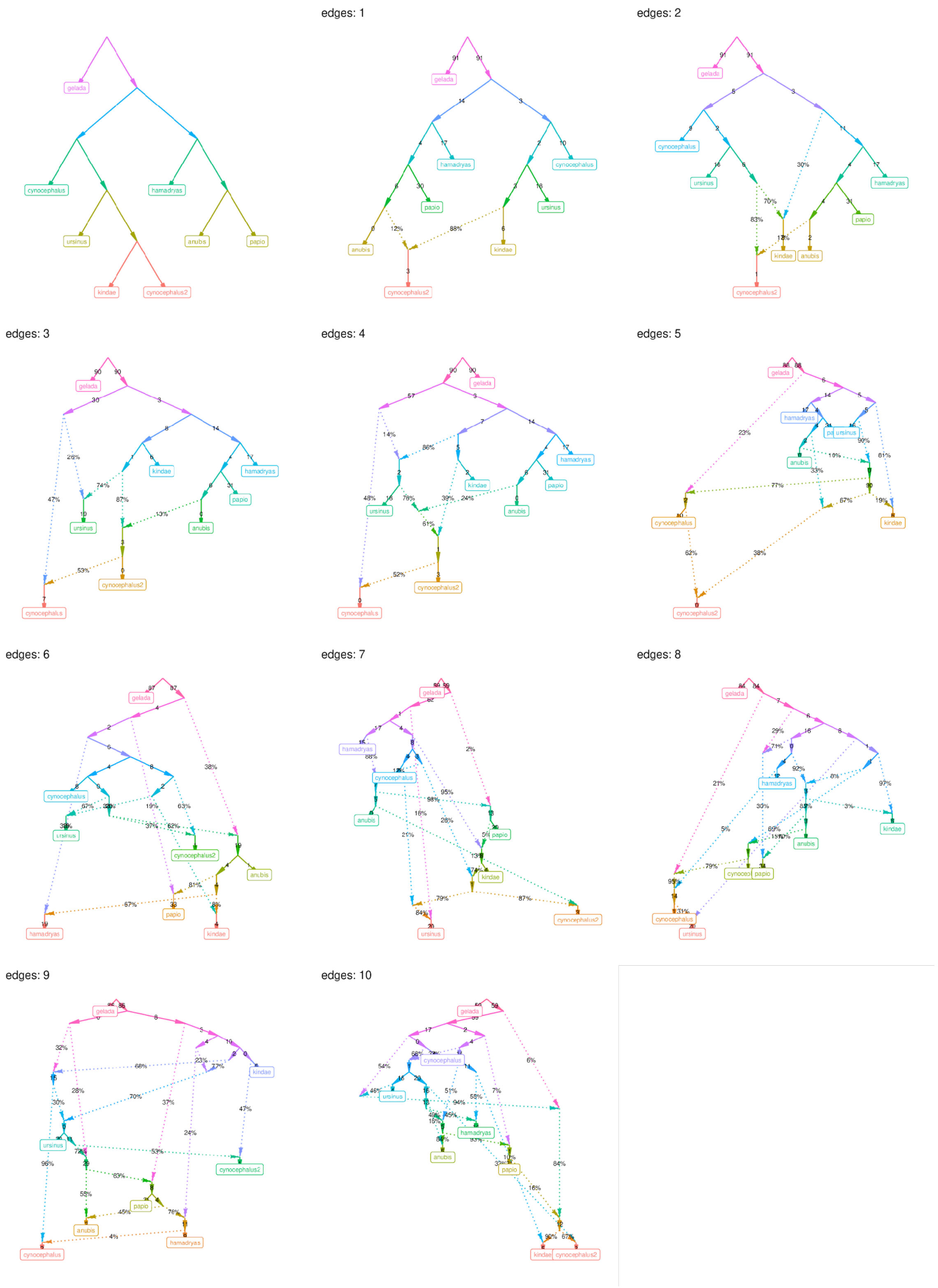


**Fig. S27.**
**ADMIXTOOLS admixture graphs.** Results of automatically retrieved graphs with up to 10 admixture events.

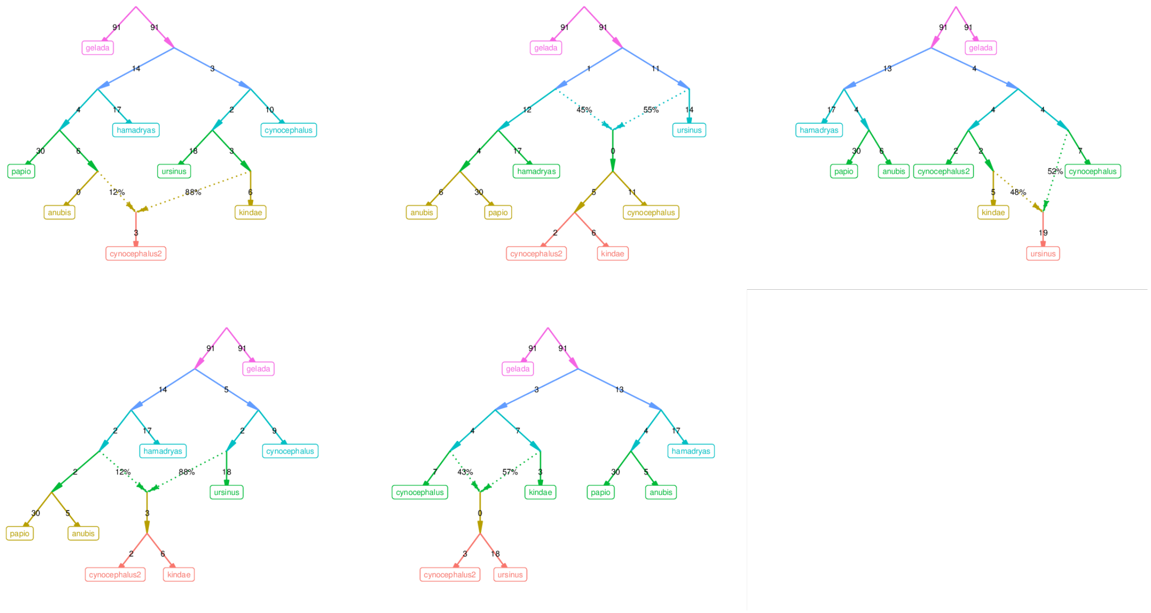


**Fig. S28.**
**ADMIXTOOLS admixture graphs.** Different admixture graphs obtained from repeating automatic graph fitting 25 times using the same dataset with one admixture event.

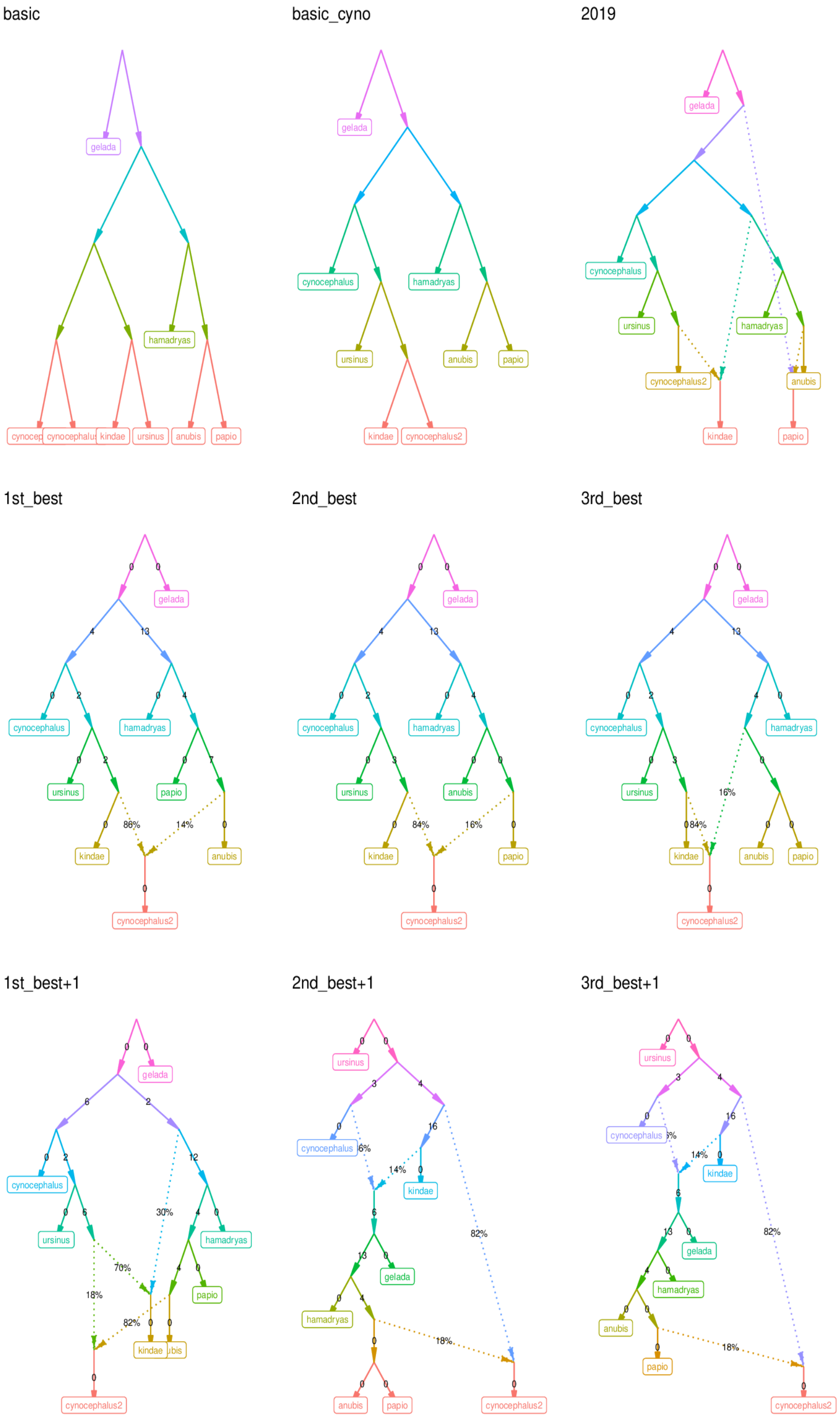


**Fig. S29.**
**ADMIXTOOLS admixture graphs.** Admixture graphs tested with f4-statistics, and best-fitting graphs with one and two admixture edges.

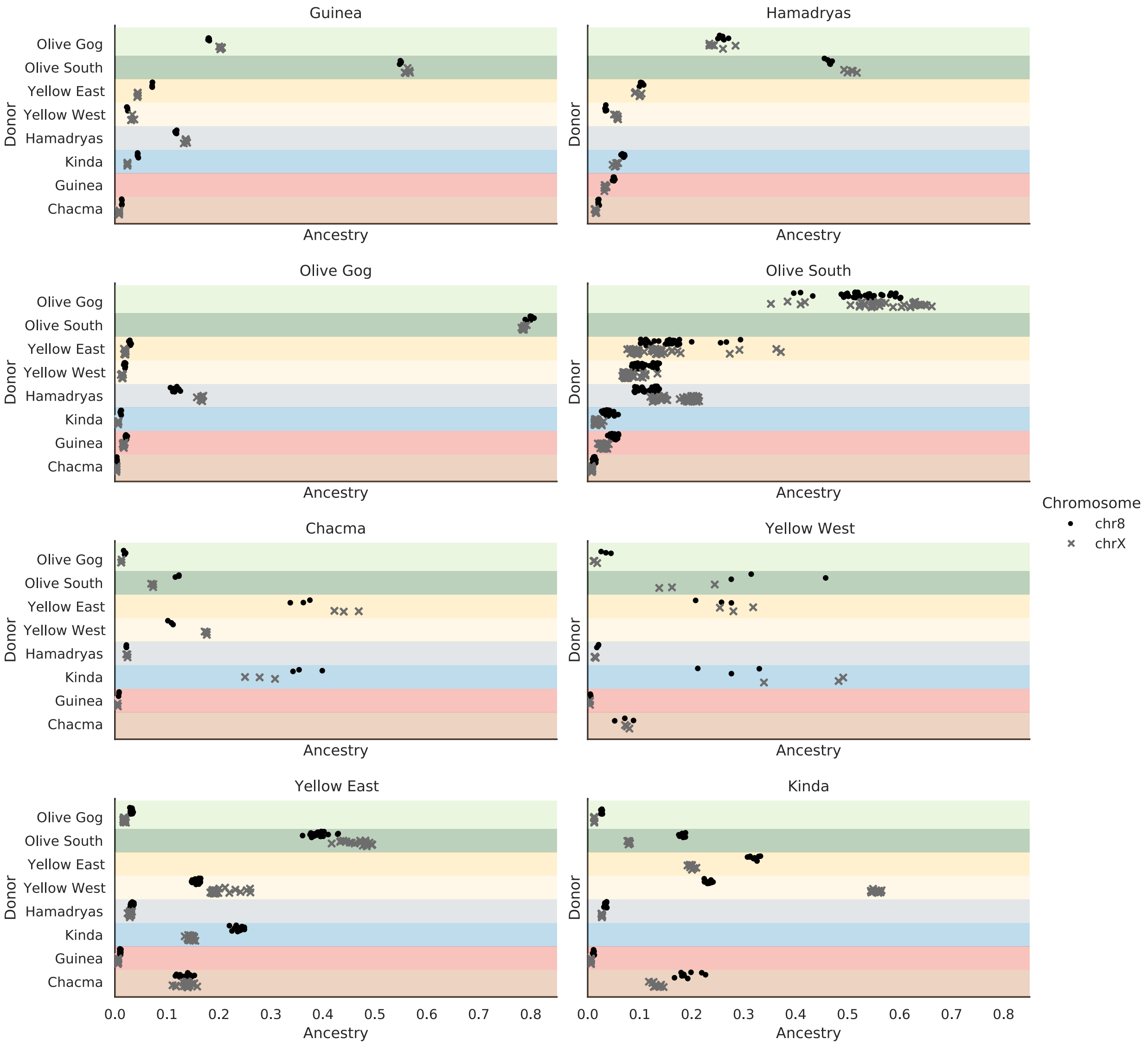


**Fig. S30.**

**Ancestry proportions of female baboons.** Each marker represents the fraction of total chromosome ancestry of one individual that is assigned to each of the remaining donor groups. Circles and crosses represent ancestry proportions of chromosomes 8 and X, respectively. The y-axis depicts the available donors in the analysis - in this figure, it is the seven other identified populations, as copying ingroup is not allowed. The x-axis shows the percentage of ancestry attributed to each other population per individual.

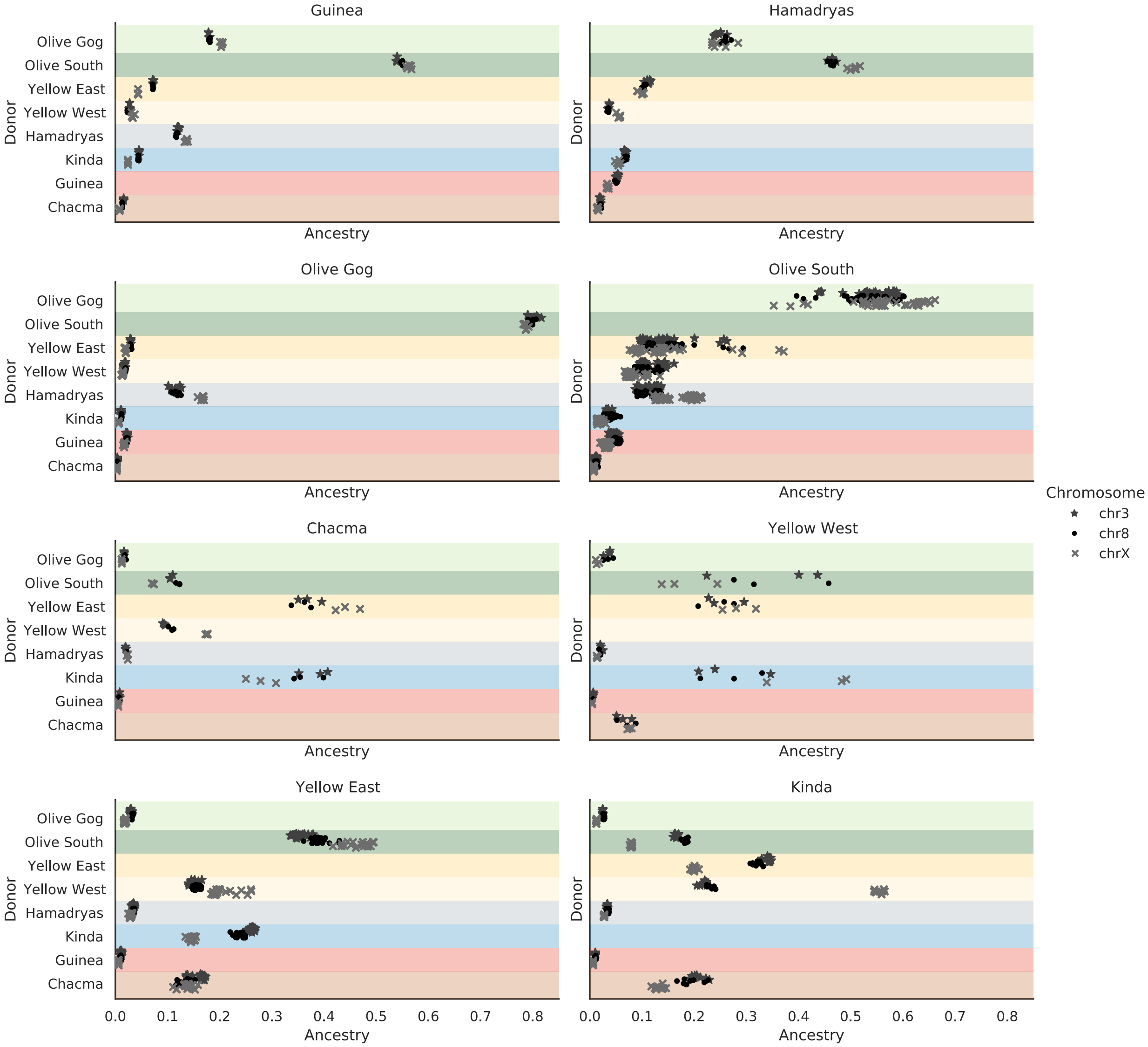


**Fig. S31.**
**Ancestry proportions of female baboons.** Same as fig. S30, but including chromosome 3 in addition to chromosome 8 for comparison to chromosome X.

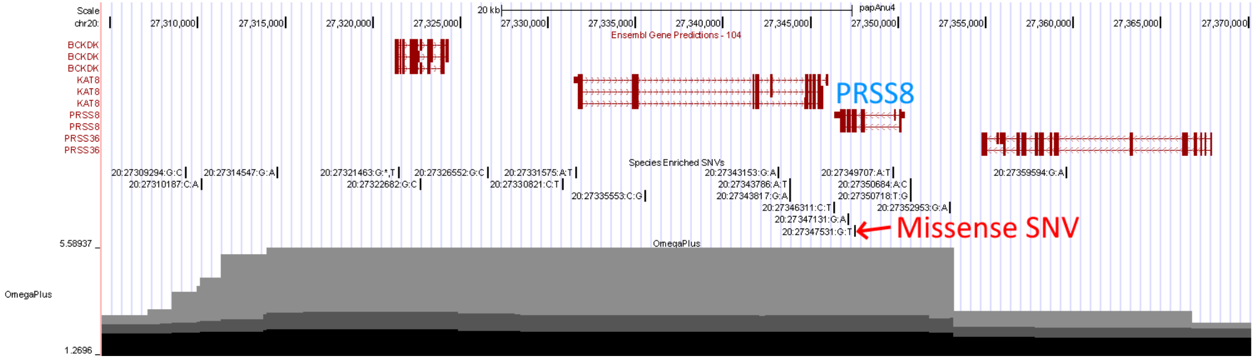


**Fig. S32.**

**Region surrounding SNV_1 (20:27347531:G:T) in hamadryas baboons**. UCSC Browser view on Panu_3.0/papAnu4 of hamadryas enriched SNVs and OmegaPlus sweep results for the region surrounding a missense SNV in Serine Protease 8 (*PRSS8*). This browser session can be viewed here: https://genome.ucsc.edu/s/Rharris1/hamadryas.PRSS8

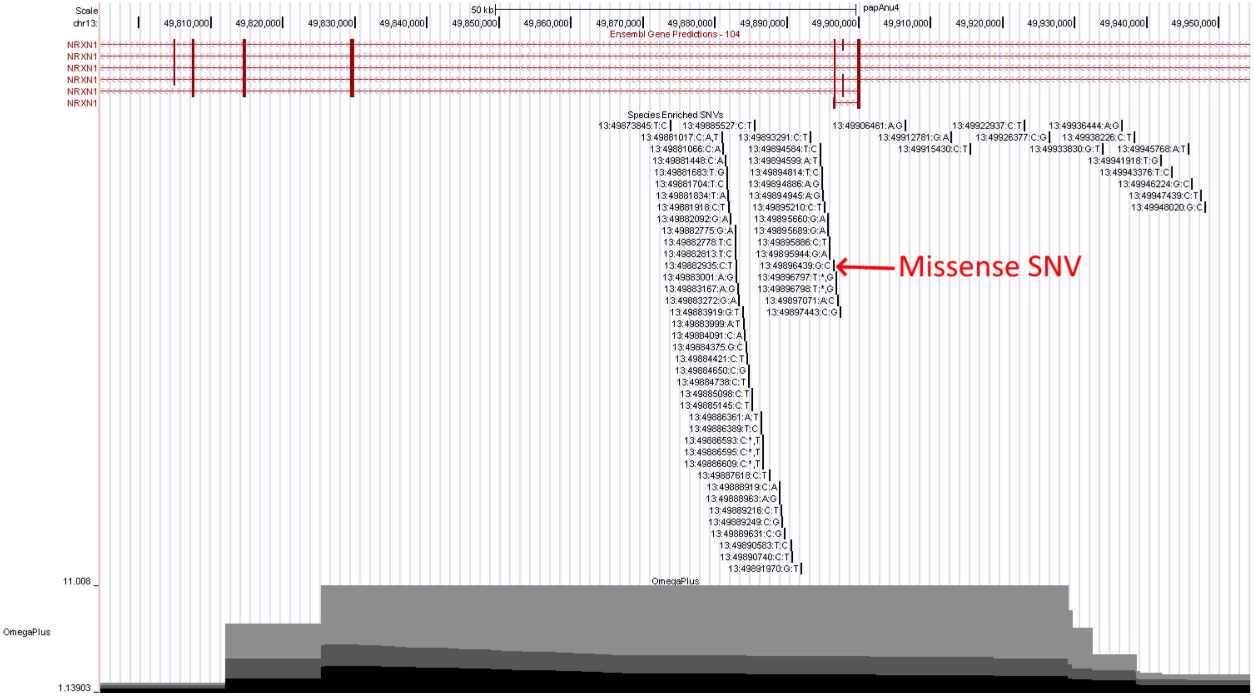


**Fig. S33.**
**Region surrounding SNV_2 (13:49896439:G:C) in hamadryas baboons.** UCSC Browser view on Panu_3.0/papAnu4 of hamadryas enriched SNVs and OmegaPlus sweep results for the region surrounding a missense SNV in Neurexin 1 (*NRXN1*). This browser session can be viewed here: https://genome.ucsc.edu/s/Rharris1/hamadryas.NRXN1

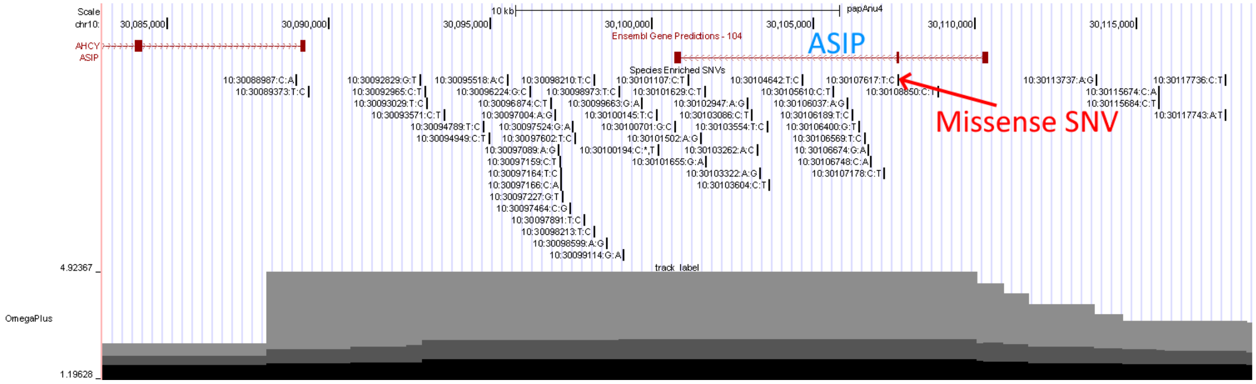


**Fig. S34.**

**Region surrounding SNV_3 (10:30107617:T:C) in Kinda baboons**. UCSC Browser view on Panu_3.0/papAnu4 of Kinda enriched SNVs and OmegaPlus sweep results for the region surrounding a missense SNV in Agouti Signaling Protein (*ASIP*). This browser session can be viewed here: https://genome.ucsc.edu/s/Rharris1/kinda.ASIP

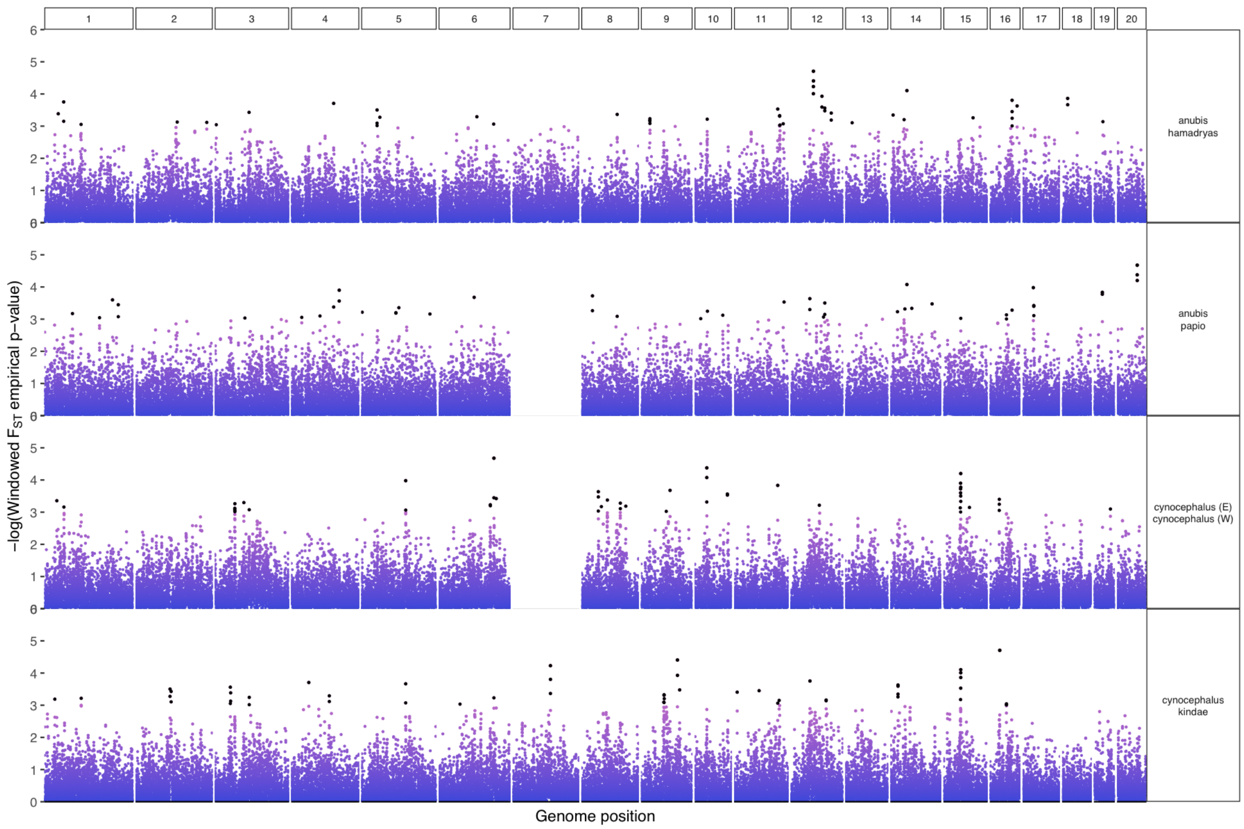


**Fig. S35.**
**Empirical p-values for differentiation metric (windowed $F_{ST}$) between pairs of species are plotted on a negative logarithmic scale**. $F_{ST}$ was computed in sliding windows of size 100kb across the genome, stepping 50kb each time. Darker colors indicate higher values of differentiation and chromosomes are listed at the top.

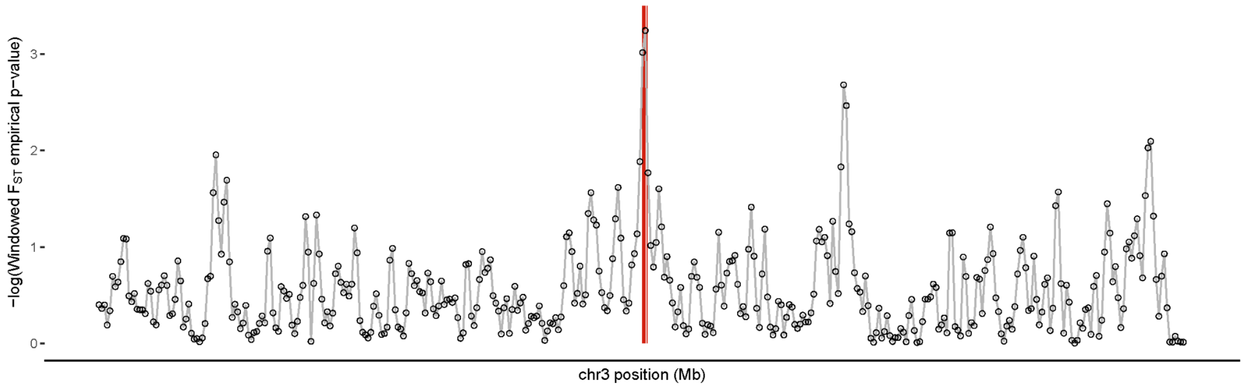


**Fig. S36.**
**Zoomed plot of empirical p-values for differentiation metric (windowed $F_{ST}$) between Kinda and yellow baboons for a portion of chromosome 3, plotted on a negative log scale.** The red line indicates the position of HOXA genes that drive the enrichment of the skeletal development/morphology-related Gene Ontology (GO) terms among genes in the most differentiated genomic regions (highest 1%).

#### Supplementary Tables

**Table S1.**
**Sample statistics based on BCFTools stats of Panu_3.0.** (.xlsx)

**Table S2.**
**List of samples with individual IDs, origin and sex.** (.xlsx)

**Table S6.**
**Globetrotter admixture dates and confidence intervals.** (.xlsx)

**Table S7.**
**Identical mitochondrial haplotypes.** (.xlsx)

**Table S9.**
**Outlier elevated FST windows (top 0.1%) for different comparisons.** Kinda versus yellow baboons, olive versus hamadryas baboons, olive versus Guinea baboons, and eastern yellow versus western yellow baboons. (.xlsx)

**Table S10.**
**Genes in outlier elevated FST windows (top 0.1%) for different comparisons.** Kinda versus yellow baboons, olive versus hamadryas baboons, olive versus Guinea baboons, and eastern yellow versus western yellow baboons. (.xlsx)

**Table S11.**
**Enriched Gene Ontology (GO) terms (FDR < 10%) for genes in outlier elevated FST windows (top 0.1%) for different comparisons.** Kinda versus yellow baboons, olive versus hamadryas baboons, olive versus Guinea baboons, and eastern yellow versus western yellow baboons. (.xlsx)

| Species or Population | Samples | Average heterozygosity |
|---|---|---|
| ***P. anubis*** | 94 | 0.001680 |
| Aberdare | 2 | 0.001768 |
| Arusha | 4 | 0.001743 |
| Gog | 25 | 0.001633 |
| Gombe | 17 | 0.001564 |
| Lake Manyara | 19 | 0.001695 |
| Ngorongoro | 6 | 0.001623 |
| Serengeti | 14 | 0.001697 |
| Tarangire | 7 | 0.002037 |
| ***P. cynocephalus*** | 62 | 0.002480 |
| Issa Valley | 1 | 0.002885 |
| Katavi | 2 | 0.002836 |
| Mahale | 7 | 0.002675 |
| Mikumi | 38 | 0.002403 |
| Ruaha | 6 | 0.002623 |
| Selous | 3 | 0.002372 |
| Udzungwa | 5 | 0.002453 |
| ***P. hamadryas*** | 26 | 0.001558 |
| ***P. papio*** | 12 | 0.000556 |
| ***P. kindae*** | 27 | 0.002553 |
| ***P. ursinus*** | 4 | 0.001986 |

**Table S3.**
**Heterozygosity estimates by species and population**. Heterozygosity was calculated as ((autosomal heterozygous SNV calls – missing SNV calls)/ungapped autosomal assembly length)) for each individual and then averaged across individuals for each species or population.

| Graph | Likelihood score | Out-of-sample score | Bootstrap p-value |
|---|---|---|---|
| edges: 0 | 15,910.1 | 8319.9 | NA |
| edges: 1 | 10,339.5 | 5729.3 | 8.34E-33 |
| edges: 2 | 6319.7 | 3074.9 | 9.8E-33 |
| edges: 3 | 1237.4 | 696 | 1.63E-80 |
| edges: 4 | 968.7 | 528.2 | 1.2E-15 |
| edges: 5 | 949.7 | 428.8 | 0.55 |
| edges: 6 | 311.7 | 177.9 | 7.12E-09 |
| edges: 7 | 38.6 | 16.3 | 9.67E-12 |
| edges: 8 | 1.3 | 1 | 0.048145 |
| edges: 9 | 0.6 | 2.3 | 0.67 |
| edges: 10 | 0.2 | 0.1 | 0.89 |

**Table S4.**
**ADMIXTOOLS admixture graphs statistics.** Likelihood and out-of-sample scores for automatically computed admixture graphs (fig. S27), as well as p-values for bootstrap-resampled graph fits.

| **Graph** | **Score f4** | **Score f3** |
|---|---|---|
| Basic | 920,874 | 27,220 |
| Basal cynocephalus | 733,376 | 15,910 |
| 2019 | 706,068 | 13,671 |
| 1_best | 356,965 | 10,951 |
| 2nd_best | 413,994 | 13,083 |
| 3rd_best | 413,994 | 13,083 |
| 1st_best+1 | 166,118 | 6737 |
| 2nd_best+1 | 141,343 | 5244 |
| 3rd_best+1 | 141,343 | 5244 |

**Table S5.**
**ADMIXTOOLS admixture graphs statistics.** Residual errors from f4-statistics and likelihood scores for f3-statistics for the models presented in fig. S29.

| Taxon | Total SNVs | Missense SNVs | Stop Gained SNVs |
|---|---|---|---|
| All 6 species | 1,342,371 | 4337 | 76 |
| Olive baboons | 27,105 | 105 | 0 |
| Hamadryas baboons | 172,002 | 555 | 6 |
| Guinea baboons | 840,650 | 2886 | 56 |
| Yellow baboons | 14,513 | 42 | 0 |
| Kinda baboons | 65,817 | 187 | 3 |
| Chacma baboons | 222,284 | 562 | 11 |

**Table S8**.

**Species Enriched SNVs**. SNVs with a PLINK p-value < $5x10^{-8}$ were filtered for those with an allele frequency > 0.9 in the target species and allele frequency < 0.1 in the other species generating a list of species enriched SNVs.
